# Comprehensive characterization of tissue-specific chromatin accessibility in L2 *Caenorhabditis elegans* nematodes

**DOI:** 10.1101/2020.09.15.299123

**Authors:** Timothy J. Durham, Riza M. Daza, Louis Gevirtzman, Darren A. Cusanovich, William Stafford Noble, Jay Shendure, Robert H. Waterston

## Abstract

Recently developed single cell technologies allow researchers to characterize cell states at ever greater resolution and scale. *C. elegans* is a particularly tractable system for studying development, and recent single cell RNA-seq studies characterized the gene expression patterns for nearly every cell type in the embryo and at the second larval stage (L2). Gene expression patterns are useful for learning about gene function and give insight into the biochemical state of different cell types; however, in order to understand these cell types, we must also determine how these gene expression levels are regulated. We present the first single cell ATAC-seq study in *C. elegans*. We collected data in L2 larvae to match the available single cell RNA-seq data set, and we identify tissue-specific chromatin accessibility patterns that align well with existing data, including the L2 single cell RNA-seq results. Using a novel implementation of the latent Dirichlet allocation algorithm, we leverage the single-cell resolution of the sci-ATAC-seq data to identify accessible loci at the level of individual cell types, providing new maps of putative cell type-specific gene regulatory sites, with promise for better understanding of cellular differentiation and gene regulation in the worm.

## 1 Introduction

Critical cellular processes are dependent on fine-tuned control of gene expression levels. From properly responding to environmental stimuli, to progressing through the stages of development and differentiation, specific changes in gene expression play an important role in facilitating precise changes in cellular state. Recent advances in single cell transcriptomics have enabled the massively-parallel measurement of gene expression, giving unprecedented genome-wide insight into which genes are regulated together in the same cell, and into expression dynamics over time. Determining which genes are important in which cells and under which conditions is critical to attaining a deeper understanding of gene function. However, in order to truly understand how gene expression reflects and influences cell state, we must also understand how it is controlled. The nematode *Caenorhabditis elegans* is a particularly powerful system in which to apply single cell genomics technologies because it has limited cell numbers that nonetheless form diverse tissue and cell types; it is very amenable to genetic manipulation; and the developmental lineage of every cell is known and invariant. In the past few years, the worm has been the subject of perhaps the most comprehensive cell type-specific metazoan gene expression atlas by single cell RNA-seq (scRNA-seq) (Cao et al., 2017; Packer et al., 2019). These studies of the *C. elegans* embryo (Packer et al., 2019) and second larval stage (L2) (Cao et al., 2017) provide a survey of the full complement of genes expressed in each major cell type, and even some cells present only once in the worm (e.g. the ASEL and ASER gustatory neurons). Now, in order to understand how these tissue-specific expression patterns arise, we also need to have a similarly comprehensive catalog of regulatory elements to map their activity in different cell types and at different stages of the life cycle.

Several efforts have been undertaken to map regulatory DNA in the worm (Araya et al., 2014; Ho et al., 2017; Daugherty et al., 2017; Kudron et al., 2018; Jänes et al., 2018). Collectively, these studies have identified tens of thousands of chromatin accessibility regions and transcription factor binding sites, using DNase-seq (Ho et al., 2017), ATAC-seq (Daugherty et al., 2017; Jänes et al., 2018), and ChIP-seq (Araya et al., 2014; Kudron et al., 2018) to assay developmental stages throughout the worm life cycle. The results convincingly show that the activity at many regulatory sites changes dramatically over the worm’s lifespan. However, the data from all of these studies is from whole worms and thus does not resolve differences in regulatory activity across cell types. The lack of cell type resolution is problematic for three main reasons. First, gene regulation is often highly cell type-specific, and even when different cell types express the same gene, they may use different enhancers or promoters to regulate that gene. In the case of two sites regulating the same gene in different cell types at the same stage, a whole worm chromatin accessibility data set would only show that both sites are accessible at the same time, but would not reveal whether the sites act in concert in the same cell type or if they affect the same gene but act in different cell types. Distinguishing between these cases is critical for understanding and modeling gene regulation. The second reason is that whole worm data may lack the sensitivity to detect regulatory events that occur in cell types that make up small fractions of the whole worm. Single cell RNA-seq data (Cao et al., 2017; Packer et al., 2019) reveal important differences in gene expression that distinguish even individual cells. Such differences are presumably driven in part by regulatory regions that are only accessible in those cells; in a whole worm assay the signal from these highly cell type-specific regions could be drowned out by the noise generated from more populous cell types. Third, the lack of cell type resolution on these regulatory DNA maps confounds our ability to draw conclusions about differential activity across development. During development, the number of cells, and with them the diversity and proportion of cell types, is constantly changing. Thus, if an accessible site is less prominent in a later larval stage compared to an embryonic stage, this change could mean the site is more accessible in embryogenesis than in later development, or it could reflect that the site is more specialized in later stages and is accessible in a smaller fraction of the cells. Given these important limitations in the available data on *C. elegans* gene regulation, we sought to generate cell type-resolved chromatin accessibility maps.

Over the past few years, the technology to collect chromatin accessibility profiles of single cells has improved greatly. This technology relies on the assay for transposase-accessible chromatin followed by high throughput sequencing (ATAC-seq) (Buenrostro et al., 2013), which treats permeabilized nuclei with a hyperactive Tn5 transposome from prokaryotes (Adey et al., 2010) to simultaneously cut accessible sites in the genome and ligate sequencing adapters onto the fragment ends on either side of the cut site (a reaction referred to as “tagmentation”). The resulting library is then amplified and sequenced. The simplicity of the assay significantly reduces the requirements for input material compared to DNase-seq (Song and Crawford, 2010), and protocols have adapted ATAC-seq to work on single cells (Buenrostro et al., 2015; Cusanovich et al., 2015; Chen et al., 2018). These and other studies have shown in multiple systems that single cell ATAC-seq (scATAC-seq) can measure thousands of sites per cell type and can identify distinct cell populations with high sensitivity. In particular, we apply the single cell combinatorial indexing variant of single cell ATAC-seq (sci-ATAC-seq). This assay probabilistically identifies DNA fragments isolated from single cells by first sorting 2500 nuclei per well into a 96 well plate and treating the nuclei in each well with a Tn5 enzyme loaded with adapters that contain a unique barcode sequence, and then pooling and re-sorting 25 nuclei per well into new 96-well plates in which a second set of barcodes is incorporated by using well-specific primers during library amplification. After sequencing, the reads can be assigned to a particular cell based on their combination of Tn5 and PCR barcodes. The sci-ATAC-seq assay has been successfully leveraged in several previous studies, including the identification of differences in gene regulation across germ layers in *Drosophila* embryogenesis (Cusanovich et al., 2018b); the generation of an atlas of 85 different clusters of cells from 13 different mouse tissues (Cusanovich et al., 2018a); and the identification of cell types in hippocampal tissue from mice (Sinnamon et al., 2019).

Here we present a comprehensive cell type-resolved map of the regulatory DNA in a whole metazoan. We collected sci-ATAC-seq data from 30,930 nuclei (hereafter referred to as cells for simplicity) isolated from a synchronized population of second larval stage (L2) *C. elegans* nematodes. To contend with low library complexity, we implement a latent Dirichlet allocation (LDA) model (Blei et al., 2003; Griffiths and Steyvers, 2004) that can parallelize the training process across multiple cores and thus is significantly faster than a previously published LDA implementation called cisTopic (González-Blas et al., 2019), although we note that the cisTopic developers have recently released an improved version. We found 37 clusters of cells that represent distinct tissue types based on mapping 38,017 peak regions to their nearest downstream genes and assessing the tissue-specific expression of those genes in L2 sci-RNA-seq data (Cao et al., 2017). Our maps of chromatin accessibility validate many previously reported regulatory sites, and also annotate almost 10,000 additional novel candidate regulatory sites, most of which are accessible only in a subset of cell types. We anticipate that these data will provide a valuable resource for studying regulatory biology in the worm and set the stage for future single cell ATAC-seq experiments on additional life stages. In conjunction with cell type-specific gene expression data, these maps of candidate regulatory regions will help reveal the gene regulatory networks driving development in *C. elegans*.

## 2 Results

### 2.1 Single cell chromatin accessibility in *C. elegans* with sci-ATAC

In order to match the sci-RNA-seq data, we grew wild-type worms to the middle of the second larval (L2) stage. At this stage, the majority of the 959 cells in the adult hermaphrodite have been produced and are terminally differentiated, but the development of the gonad has not progressed far enough to begin producing the thousands of germline nuclei that eventually outnumber the somatic nuclei in the adult and would bias our collection of tissue types. After harvesting the worms, we fixed and isolated the nuclei, froze them in aliquots, and used these wild-type nuclei as input to the single cell combinatorial indexing assay for transposase-accessible chromatin (sci-ATAC-seq) (Cusanovich et al., 2015, 2018b,a). We collected sci-ATAC-seq data for 30,930 cells with at least 150 unique reads per cell (median 672 reads per cell), which represents about 40x sampling of each cell in the L2 worm (Fig. S1). Note that we expect thousands of genes to be expressed per cell, and thus at 672 reads per cell we are only sampling a small fraction of the accessible regulatory sites in any given cell.

The post-sequencing pipeline consists of aligning the paired-end reads to the WS235/ce11 version of the *C. elegans* genome, and identifying cut sites as the 60 bp regions centered on the ends of the DNA fragments identified by the mapped mate pairs. (See methods for details.) Next we needed to identify which loci were accessible in each of our cells. There exists no unbiased annotation of cell type-resolved regulatory regions in *C. elegans*, so we called peaks directly from the sci-ATAC-seq data in an iterative fashion (Fig. 1). The first step was to call peaks with MACS2 (Zhang et al., 2008) using all of the reads together, as with a bulk ATAC-seq data set. To detect some more cell type-specific peaks that could be obscured by the background reads coming from the most abundant cell types in this complex cell mixture, we clustered the cells based on the output of the latent Dirichlet allocation (LDA) modeling technique (Blei et al., 2003) using the bulk peaks, pooled the cut sites from the cells in each cluster, and called peaks for each cluster using MACS2 (“Primary LDA” in Fig. 1b). Finally, in order to refine our peak set we repeated the clustering, pooling, and peak calling once more (“Refinement LDA” in Fig. 1b).

**Figure 1:**
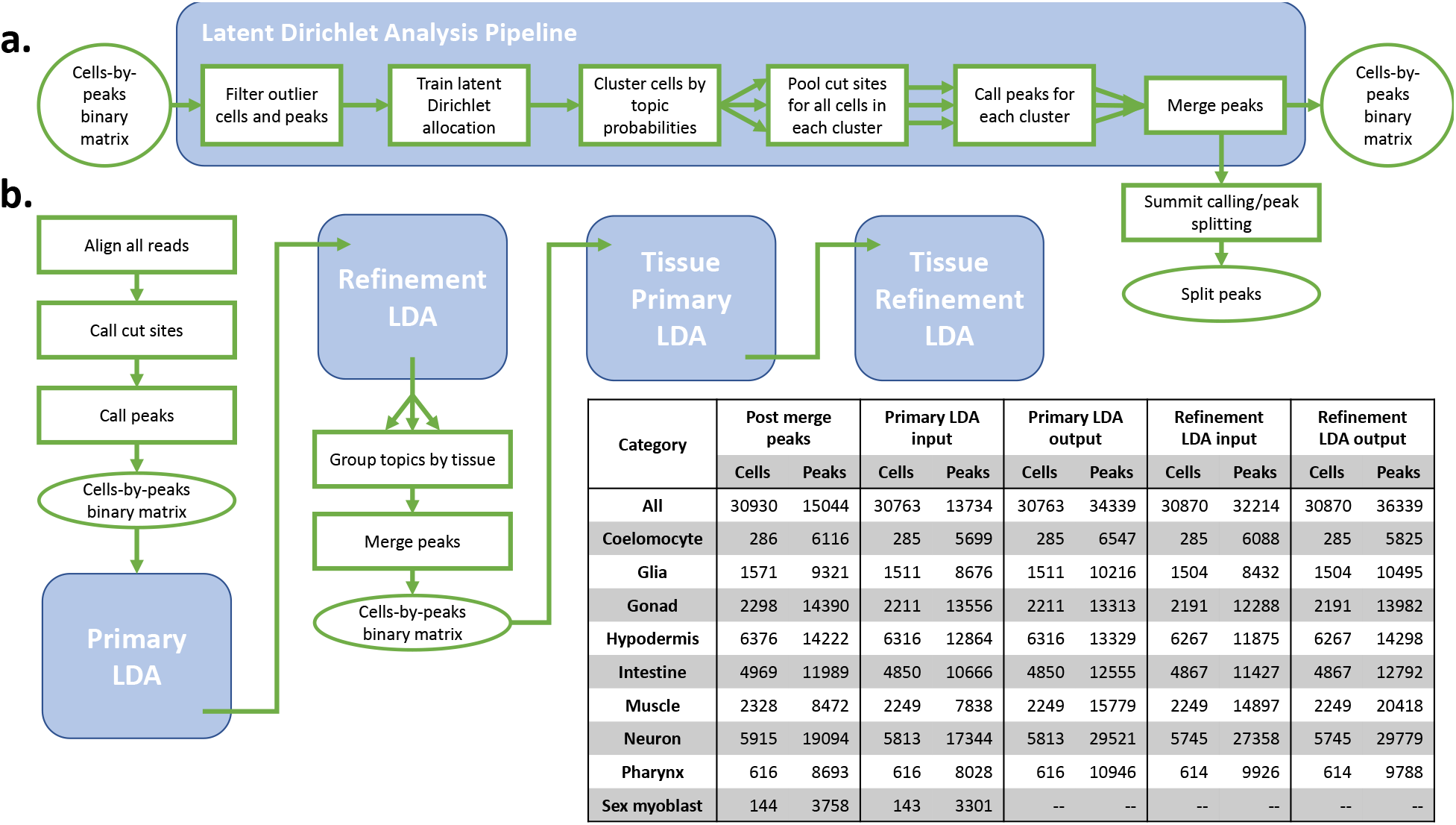
An iterative peak-calling procedure yields more peaks from the complex mix of worm cell types. **a.** The core peak-calling procedure is to model the data using latent Dirichlet allocation, cluster the cells, and call peaks based on the clusters. **b.** A flow chart represents the overall peak calling strategy. First, bulk peaks are called, followed by two iterations of clustering and peak calling based on an LDA model. Then, we group the cells by tissue and repeat the two steps of clustering and peak calling. The number of cells included and the number of peaks called at each step is given in the inset table.

Each peak calling iteration centers around modeling the peak distribution in individual cells, as represented by a cells-by-peaks matrix containing binary values indicating whether or not a cell reports a read overlapping a particular peak. This binary matrix is input into LDA, a Bayesian modeling approach that was originally developed in the field of semantic analysis of text documents, and has recently been used to model single cell genomics data (González-Blas et al., 2019; Kim et al., 2019; Dey et al., 2017). We discuss this modeling approach in detail below and in Methods. After our iterative clustering and peak-calling procedure, we post-processed the merged peaks from the refinement LDA step by detecting local maxima (summits) in the signal over each peak and splitting any peaks with multiple summits into separate contiguous segments. This step splits wide peaks to better capture accessible regions that may contain multiple binding sites. We share the pipeline output in a UCSC track hub, which can be accessed at the following URL: http://genome.ucsc.edu/cgi-bin/hgTracks?db=ce11&hubClear=http://waterston.gs.washington.edu/atacTissue/Durham_hub.txt.

### 2.2 Single cell peaks are concordant with regulatory regions from published bulk chromatin assays

After applying our iterative peak calling procedure, the whole worm refinement LDA (Fig. 1) identified 36,339 peak regions, and splitting the multi-summit peaks resulted in a total of 38,017 peaks. To compare our peak calls to other maps of regulatory DNA in *C. elegans*, we intersected our regions with peaks from two other data sets: bulk, whole worm ATAC-seq data from samples spanning the *C. elegans* life cycle (Jänes et al., 2018), and transcription factor (TF) binding site peaks identified from 427 whole-worm TF ChIP-seq data sets from the modERN consortium (Kudron et al., 2018) (Fig. 2).

**Figure 2:**
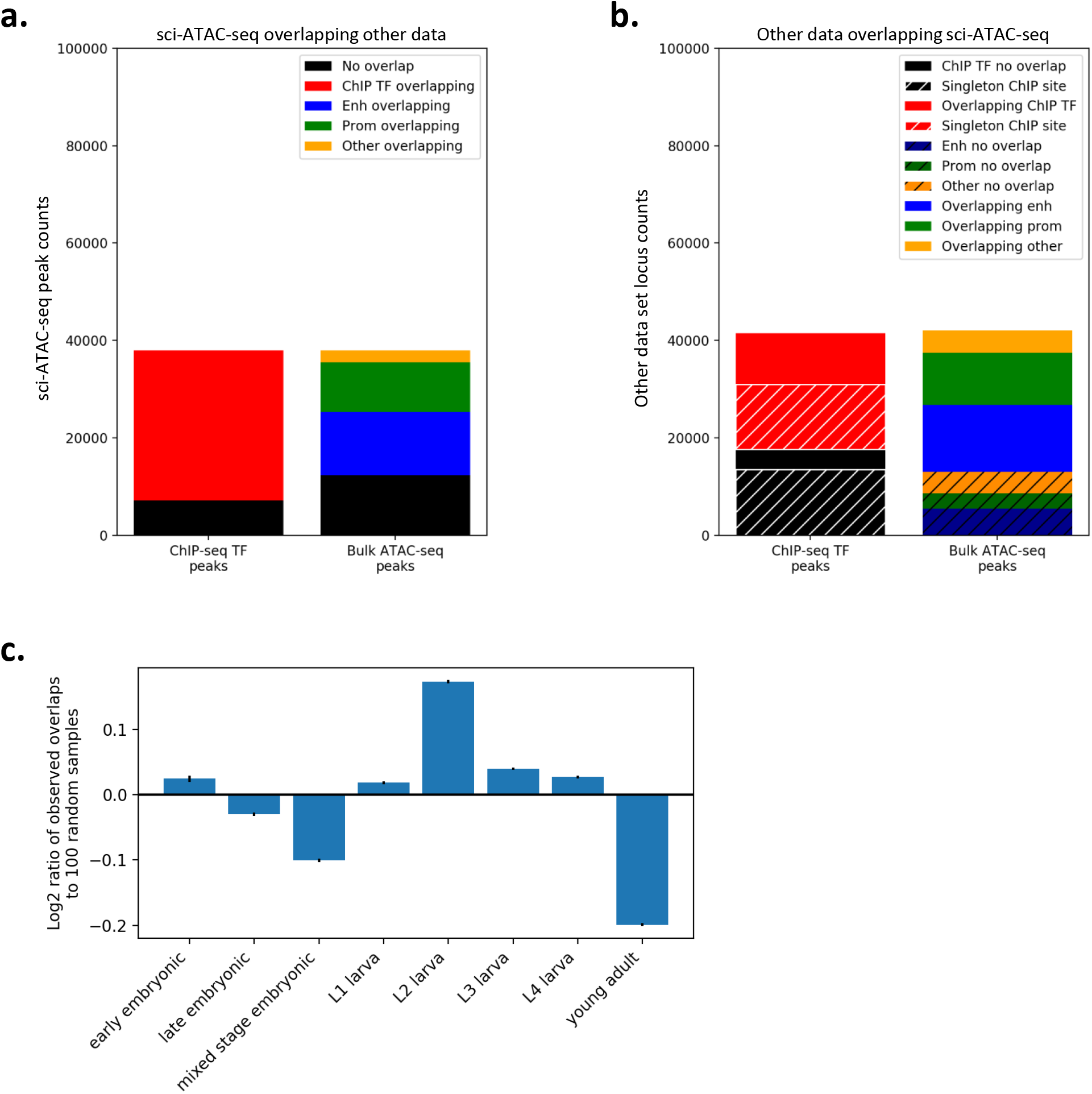
The peaks called from sci-ATAC-seq data exhibit substantial overlap with existing chromatin data collected from whole worms. **a.** The vast majority of sci-ATAC-seq peaks overlap sites called in the TF ChIP-seq peaks from modERN or the bulk ATAC-seq peaks from Jänes, et al. 2018. **b.** Peaks from the other data sets show substantial overlap with sci-ATAC-seq peaks. Most of the ChIP-seq TF peaks that do not overlap a sci-ATAC-seq peak are singleton peaks that are only found in a single experiment. **c.** Breaking out the ChIP-seq peak overlaps by the developmental stage of the worms assayed and comparing the distribution across stages of the peaks with overlaps compared to the stage distribution for randomly selected ChIP-seq peaks shows an enrichment for peaks found in larval stage L2. Error bars indicate the 95% confidence interval.

As expected, we find good overlap with both published data sets but also some differences (Fig. 2a). First, intersecting the sci-ATAC-seq peaks with the bulk ATAC-seq data set shows 25,675 peaks of 38,017 (~ 66%) overlapping a bulk ATAC-seq site overall. Note that our sci-ATAC-seq peaks cover 20,234,260 bp in total, which is about 20.2% of the genome, while the bulk ATAC-seq peaks cover 6,376,655 bp, or only about 6.4% of the genome; so this overlap is highly significant (Fisher’s exact test *p* = 0). About 50% (12,960) of the overlaps were with bulk ATAC-seq sites classified as enhancers (Jänes et al., 2018), about 40% (10,219) were with sites classified as promoters, and 10% (2,496) were with other kinds of sites (e.g. non-coding RNAs). We find even more extensive overlap with TF ChIP-seq sites; 30,886 of the 38,017 sci-ATAC-seq peaks (~ 81%) overlap TF ChIP-seq peaks from modERN (Fig. 2a). The ChIP TF peaks cover 23,225,218 bp (~ 23.2% of the genome), and similarly to the bulk ATAC-seq overlaps the number of TF ChIP-seq peakd overlaps is highly significant (Fisher’s exact test p = 0).

Looking at the data from the opposite perspective, almost three quarters of bulk ATAC-seq sites overlap a sci-ATAC-seq peak (29,021 of 42,102, ~ 69%), and these overlaps are fairly evenly split between sites classified as promoters (10,684 of 13,833, ~ 77%) and enhancers (13,715 of 19,195, ~ 71%), with the remaining overlaps (4,622 of 9,074, ~ 51%) involving other smaller categories of regulatory elements. In the case of the modERN TF sites, we find that the majority overlap a sci-ATAC-seq peak (23,863 of 41,542, ~ 57%). Most of the ChIP-seq sites that do not overlap a sci-ATAC-seq peak are “singletons” that were only observed in one of the 427 ChIP-seq data sets (13,579 of 17,679, ~ 77%). Previous work suggests that singleton sites are enriched in false positives (Araya et al., 2014), which would be less likely to appear in an orthogonal assay like sci-ATAC-seq. Alternatively, some of these ChIP sites could be tissue- or stage-specific ChIP-seq sites that may not be accessible in the L2 stage that we assayed here. Indeed, we find that the set of ChIP-seq sites that overlap sci-ATAC-seq peaks is enriched for those found in larval samples, especially L2, and depleted for sites observed in embryo and young adult (Fig. 2c).

In addition, other important technical differences in the data collection methods likely also contribute to differences in the regulatory sites detected. First, we expect that whole worm bulk assays have less sensitivity to detect cell type-specific sites than sci-ATAC-seq. Second, in the case of the ChIP-seq data, the recovered sites are biased for the particular set of TFs that are assayed; in contrast, sci-ATAC-seq is relatively unbiased, but is subject to the Tn5 sequence bias (Green et al., 2012). Third, ChIP-seq data is known to have artifacts associated with the antibody used for immunoprecipitation (Araya et al., 2014; Kudron et al., 2018).

### 2.3 Latent Dirichlet allocation modeling reveals 37 clusters of cells

In order to interpret our data at the level of tissues and cell types, we applied latent Dirichlet allocation (LDA) (Blei et al., 2003), a statistical modeling technique that is particularly well-suited to finding patterns in sparse data from single cell genomics assays (Fig. 3a). LDA has been successfully used to analyze single cell chromatin accessibility (González-Blas et al., 2019), single cell chromatin conformation (Kim et al., 2019), and single cell gene expression data (Dey et al., 2017). LDA is a generative Bayesian modeling approach that was developed in the context of document classification. In the document classification task, the model is trained to identify information-rich words in a document corpus and to associate those words with latent topics that can distinguish the documents. The output consists of two matrices: one that captures the probability distribution of each topic over all words, and another that captures the probability distribution of each document over all topics. Thus, each topic is defined as some combination of words, and each document is modeled with some combination of topics based on its word content.

**Figure 3:**
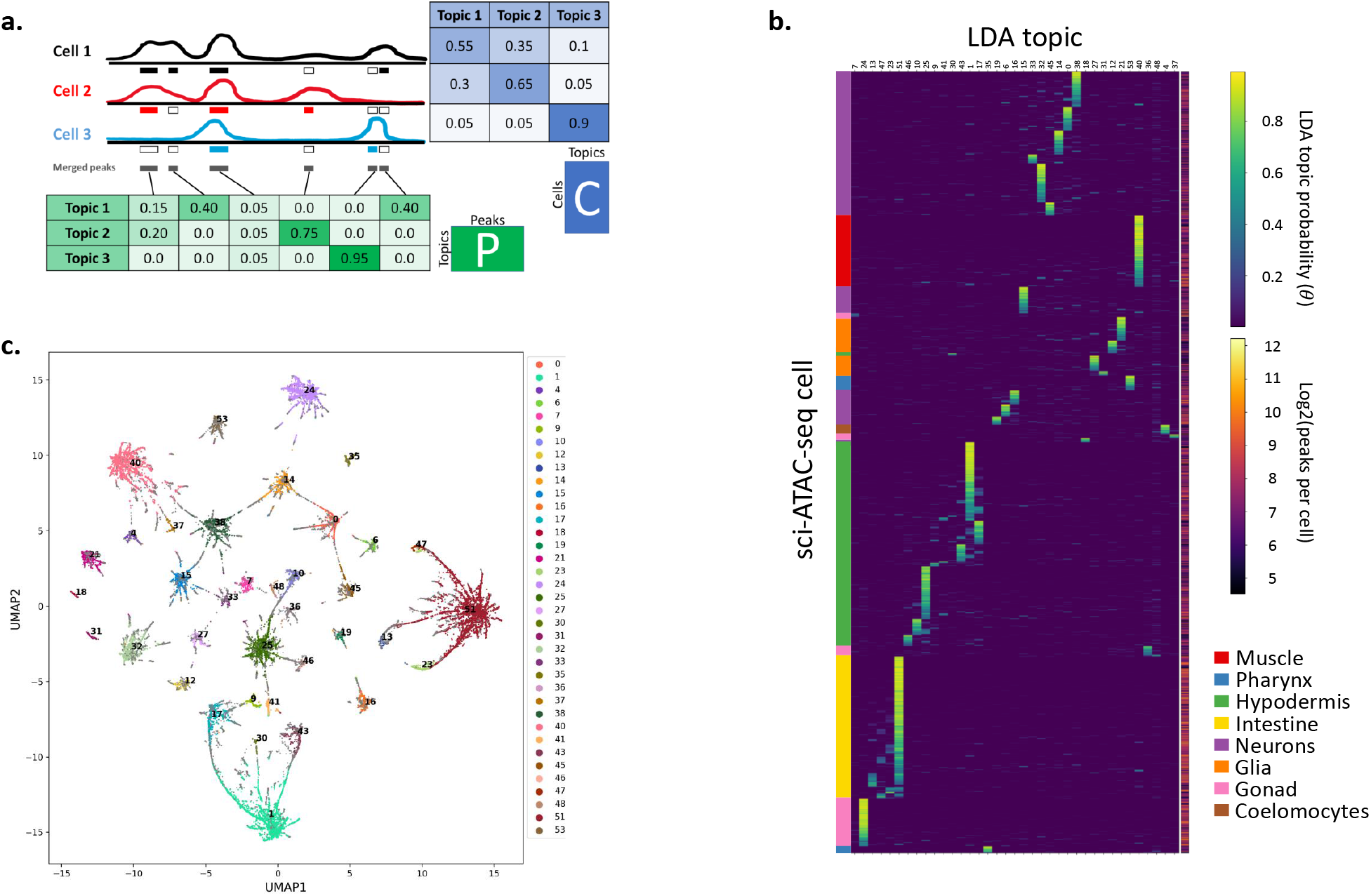
Latent Dirichlet allocation modeling yields 37 major cell clusters that are characterized mostly by a single topic each. **a.** LDA modeling learns latent topics that explain the data and return two matrices, here designated *P* and *C*. Matrix *P*, referred to in the text as the peaks-by-topics matrix, captures the probability distribution of each topic over all peaks, while matrix *C*, referred to in the text as the cells-by-topics matrix, captures the probability distribution of each cell over all topics. **b.** Heatmap showing the normalized C matrix values for the 37 topics associated with clusters; this plot highlights that most cells have probability concentrated in one or a few topics. Cell types determined for the topics based on analysis of the P matrix are annotated on the left, and the number of peaks per cell is shown to the right. **c.** UMAP embedding of the C matrix colored to indicate the 37 cell clusters. Any cells that are not assigned to a cluster are plotted as small gray dots and are mostly found on the periphery of the clusters.

When applied to scATAC-seq data, cells are treated as documents, and peaks are treated as words. The model learns the peaks associated with latent “regulatory topics” that capture patterns that discriminate among regulatory states and cell types. The output consists of two matrices: one representing the distribution of peaks over topics, and another representing the distribution of topics over cells. A key advantage of LDA in this setting is that it handles sparse data quite well and leverages information from all cells at once to assign peaks to topics and all peaks at once to assign topics to cells.

We trained an LDA model with 55 topics, choosing 55 topics based on a 5-fold cross-validation hyperparameter search procedure (see Methods, Fig. S4). This model yielded a cells-by-topics matrix with 30,870 rows (one for each cell after filtering out cells with too few peaks detected) and 55 columns, and a peaks-by-topics matrix with 32,214 rows (one for each peak after filtering out outlier high- and low-coverage peaks) and 55 columns (Fig. 3a). [Note that in the text we transpose the topics-by-peaks matrix and refer to it as the peaks-by-topics matrix for consistency with the cells-by-topics matrix.] We expected that differences in chromatin accessibility among cell types would be the largest source of covariation in the data, and thus that there would be many topics that corresponded to distinct cell types.

In order to look for topics that might distinguish cell types, we first tried to identify topics that did not seem to have particularly high probability in any subset of cells and remove those from consideration. For each topic, we ranked the cells by their probability for that topic, then took the mean topic probability distribution for the top 50 cells. We then calculated the similarity between the mean topic distribution and that of each of the top 50 cells by taking the dot product, and then took the average of those dot products to yield an average similarity for the top 50 cells. After examining the similarity distribution across all topics, we set a cutoff of 0.2 and removed 15 topics, leaving 40 topics (Fig. S6). Next, we assigned cells to clusters based on their highest-probability topic, and we refer to the resulting clusters of cells as “topic clusters”. Any cell with greater than 50% probability for one of the 40 topics was automatically added to that topic’s cluster. For any topic cluster with fewer than 150 cells, we attempted to assign unassigned cells (i.e. that had less than 50% probability for all topics) from the periphery based on their Euclidean distance from the centroid of the topic in LDA cells-by-topics coordinate space. After attempting to add unassigned cells, we removed three topics that still had fewer than 50 cells because they would have too little coverage for peak calling. Thus, using this procedure we assigned a total of 24,503 cells to 37 topic clusters for further analysis (Table 1).

**Table 1:**
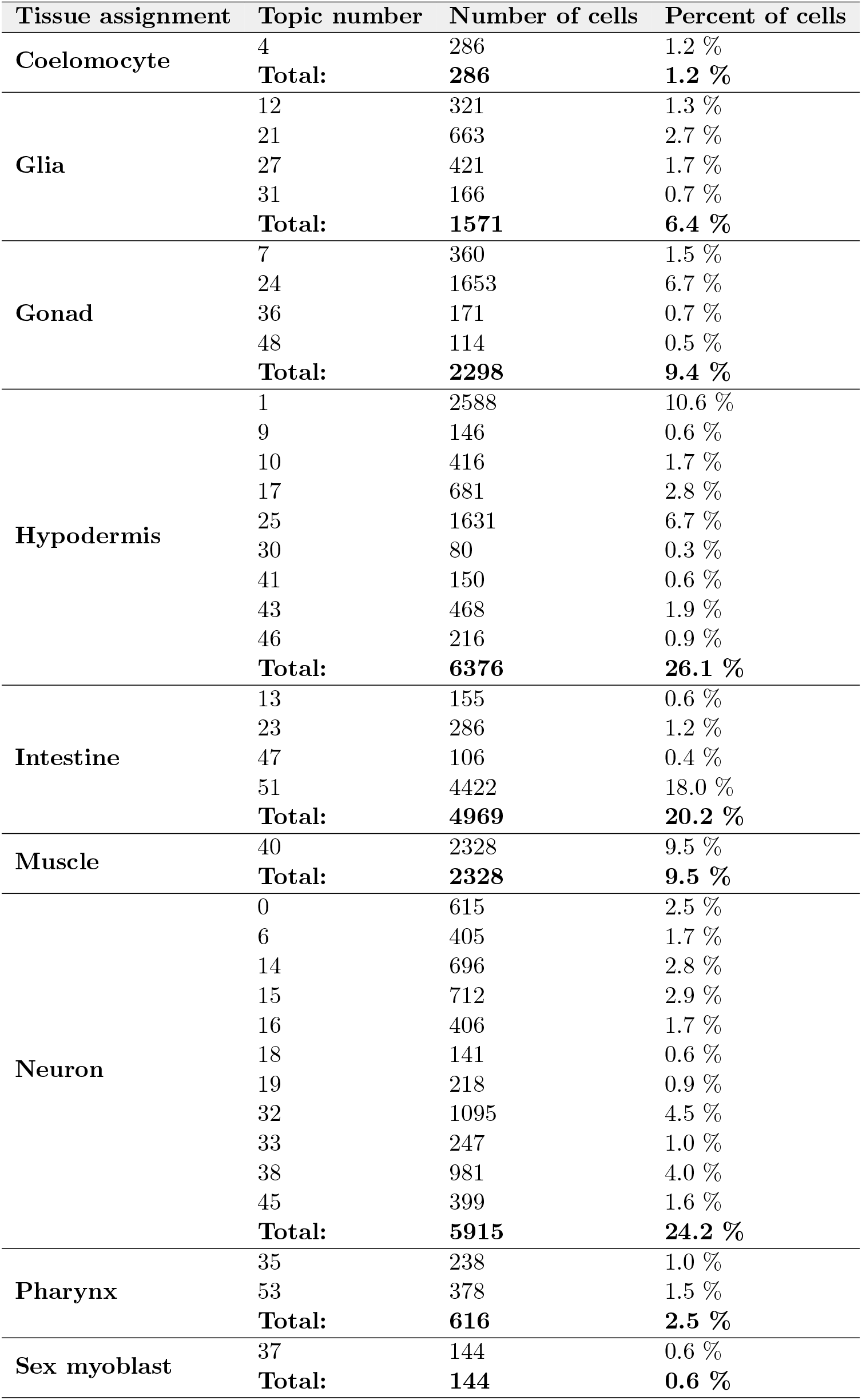
Results of iterative LDA procedure, with the number cells assigned to each topic cluster, the tissue assignment, and the percentage of all 24,503 cells that were assigned to a topic cluster.

| Tissue assignment | Topic number | Number of cells | Percent of cells |
| --- | --- | --- | --- |
| Coelomocyte | 4 | 286 | 1.2 % |
|  | **Total:** | **286** | **1.2 %** |
| Glia | 12 | 321 | 1.3 % |
|  | 21 | 663 | 2.7 % |
|  | 27 | 421 | 1.7 % |
|  | 31 | 166 | 0.7 % |
|  | **Total:** | **1571** | **6.4 %** |
| Gonad | 7 | 360 | 1.5 % |
|  | 24 | 1653 | 6.7 % |
|  | 36 | 171 | 0.7 % |
|  | 48 | 114 | 0.5 % |
|  | **Total:** | **2298** | **9.4 %** |
| Hypodermis | 1 | 2588 | 10.6 % |
|  | 9 | 146 | 0.6 % |
|  | 10 | 416 | 1.7 % |
|  | 17 | 681 | 2.8 % |
|  | 25 | 1631 | 6.7 % |
|  | 30 | 80 | 0.3 % |
|  | 41 | 150 | 0.6 % |
|  | 43 | 468 | 1.9 % |
|  | 46 | 216 | 0.9 % |
|  | **Total:** | **6376** | **26.1 %** |
| Intestine | 13 | 155 | 0.6 % |
|  | 23 | 286 | 1.2 % |
|  | 47 | 106 | 0.4 % |
|  | 51 | 4422 | 18.0 % |
|  | **Total:** | **4969** | **20.2 %** |
| Muscle | 40 | 2328 | 9.5 % |
|  | **Total:** | **2328** | **9.5 %** |
| Neuron | 0 | 615 | 2.5 % |
|  | 6 | 405 | 1.7 % |
|  | 14 | 696 | 2.8 % |
|  | 15 | 712 | 2.9 % |
|  | 16 | 406 | 1.7 % |
|  | 18 | 141 | 0.6 % |
|  | 19 | 218 | 0.9 % |
|  | 32 | 1095 | 4.5 % |
|  | 33 | 247 | 1.0 % |
|  | 38 | 981 | 4.0 % |
|  | 45 | 399 | 1.6 % |
|  | **Total:** | **5915** | **24.2 %** |
| Pharynx | 35 | 238 | 1.0 % |
|  | 53 | 378 | 1.5 % |
|  | **Total:** | **616** | **2.5 %** |
| Sex myoblast | 37 | 144 | 0.6 % |
|  | **Total:** | **144** | **0.6 %** |

Visualizing these clusters with Uniform Manifold Approximation and Projection (UMAP) (McInnes et al., 2018; Becht et al., 2019) shows clear separation among groups of cells (Fig. 3c). We also note that, although we focus on the 37 topics that drive our cell clusters, the remaining topics could still contain useful information. Some topics with high probability in too few cells to be used for clustering may characterize cell types that are rare or that were less-successfully isolated. In addition, topics with more diffuse probability across cells could correspond to other phenomena, for example, to different kinds of regulatory activity (e.g. promoters or enhancers (González-Blas et al., 2019)), cells with more complex patterns of regulatory activity, cells with noisy signal, or cell types with insufficient signal to be confidently clustered.

### 2.4 Topics correspond to specific tissue identities

After clustering our cells based on 37 topics, we sought to determine whether these clusters of cells represent different cell types. As with other dimensionality reduction techniques (e.g. principal component analysis), LDA is an unsupervised algorithm with no restrictions on what qualities of the data it uses to determine the topics, and interpretation of the topics can be challenging.

One way to assess whether the topics show some tissue-specificity is by cross-referencing the sci-ATAC-seq peaks with what is known about those loci in the literature, similarly to how marker genes are identified for clusters in scRNA-seq data (Cao et al., 2017; Packer et al., 2019). In the absence of broad data sets for cell-specific regulatory elements, we began by looking for overlap of the topic specific ATAC-seq peaks with the ChIP-seq peaks from cell type-specific transcription factors. For each of the 37 topics that we used to cluster the cells, we found all peaks in the peaks-by-topics matrix with probability greater than zero for that topic and overlapped them with all available ChIP-seq peaks from sites found in 40 or fewer other ChIP-seq data sets (i.e. non-high-occupancy-target, or non-HOT, sites (Kudron et al., 2018)) for three transcription factors with known cell type-specific expression patterns: HLH-1, a master regulator for body wall muscle (Krause et al., 1990); ELT-1, a master regulator for hypodermis in embryos and seam cells in L2 larvae (Page et al., 1997); and ELT-2, a transcription factor important in intestine development (Fukushige et al., 1998). We compared the number of overlaps in each topic to the number we would expect if the overlaps were random (i.e. if topics were not cell type-specific), and expressed this comparison as a log_2_ ratio between observed and random overlap counts (Fig. 4). We find topics with specific enrichment for overlaps with peaks from each transcription factor (95% confidence intervals are provided in Fig. 4). Peaks that characterize topics 37 and 40 are most enriched for overlaps with HLH-1 sites; peaks for topics 9, 10, 25, and 46 are particularly enriched for overlaps with ELT-1 sites; and peaks for topics 13, 23, 47, and 51 are most enriched for overlaps with ELT-2 sites. This analysis suggests that at least some of the topics are representing different tissues, and that the cells enriched in topics characterized by many peaks overlapping HLH-1, ELT-1, and ELT-2 sites are likely muscle, hypodermis, and intestine cells, respectively. We show the same overlap analysis for all 283 transcription factors in Figure S8.

**Figure 4:**
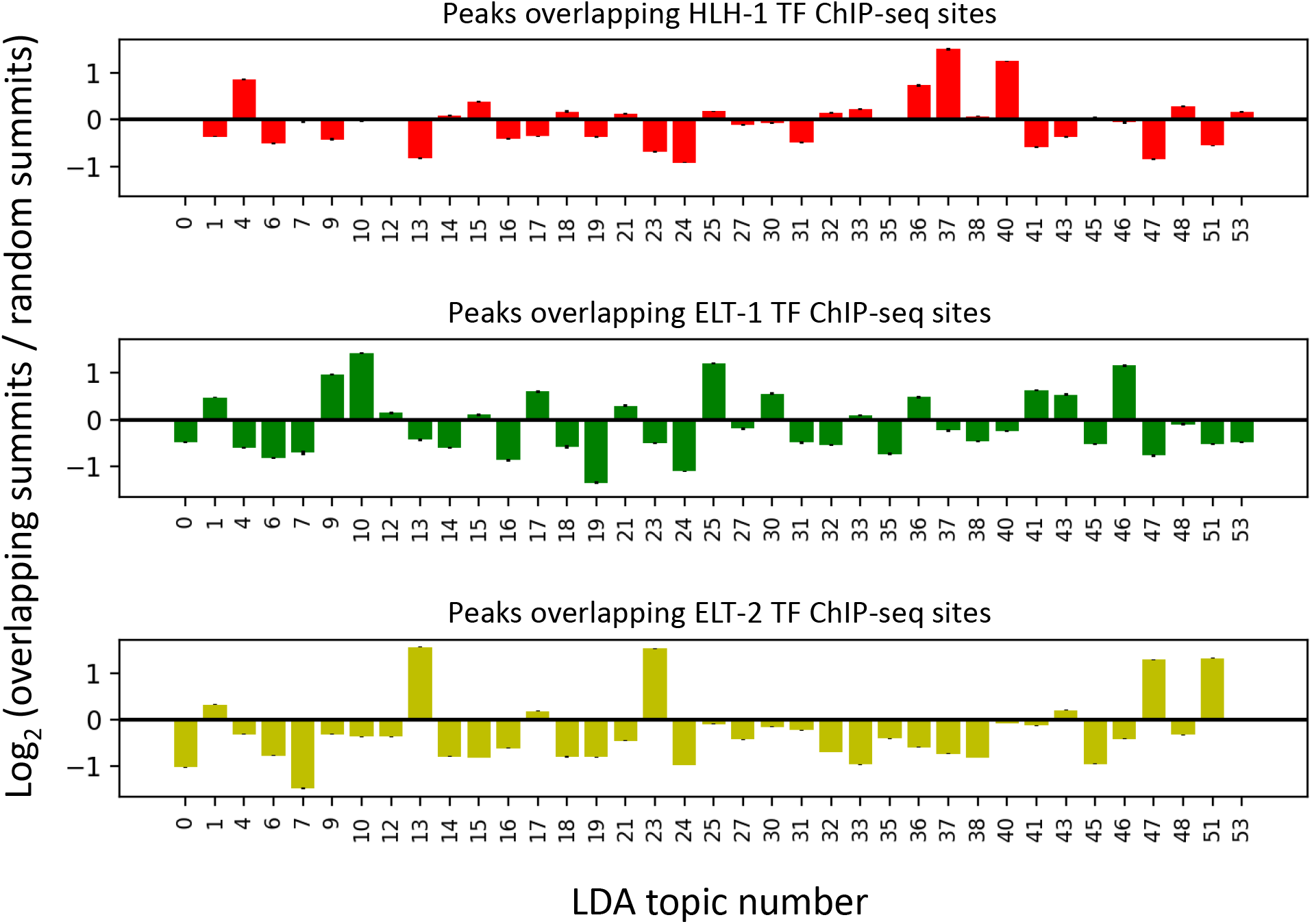
Overlapping peaks important for each topic with ChIP-seq peaks collected from cell type-specific TFs suggests at least some topics represent tissue types. Peaks associated with each topic were overlapped with ChIP-seq peaks for three cell type-specific transcription factors: HLH-1, which is specific for muscle (top plot); ELT-1, which is specific for seam cells (middle); and ELT-2, which is specific for intestine (bottom). Topic distributions for peaks with ChIP-seq site overlaps were compared to the topic distribution for randomly sampled peaks, and the results are plotted here as the log_2_ ratio of the overlap topic distribution to the random topic distribution. Error bars represent the 95% confidence interval for 100 random samples.

Encouraged by the analysis of overlaps with ChIP-seq data from cell type-specific TFs, we sought to leverage the L2 sci-RNA-seq data (Cao et al., 2017) to do a more comprehensive analysis of all 37 topics. In order to do this, we mapped the sci-ATAC-seq peaks to the nearest downstream exon within 1200 bp, thereby accounting for peaks at alternative promoters and within introns. Assigning regulatory regions to their nearest gene is a commonly used (Araya et al., 2014) but rather naïve heuristic that works well in *C. elegans*. In contrast to mammals and other more complex animals (Pliner et al., 2018), there is little evidence that regulation by distal sites plays a prominent role in *C. elegans* (Reinke, 2013). Furthermore, the *C. elegans* genome is compact and gene-dense, meaning that most regulatory sites are found close to genes, either in intergenic sequences or in introns (Reinke, 2013). In total, we were able to assign 19,532 peaks of our 36,339 total peak calls to 17,389 genes. The number of genes we associate with accessible regions is higher than the number of genes known to be expressed in L2 worms (Boeck et al., 2016; Cao et al., 2017). We believe there are two reasons for this discrepancy: first, the RefSeq database that we used includes non-coding genes that are not found in the sci-RNA-seq data (and thus excluded from the following analysis), and second, some accessible sites are likely hosting regulatory factors playing a repressive role and thus are not necessarily associated with productive transcription that would be detected by RNA-seq.

With these caveats, we used these peak-gene assignments to associate genes with topics and thereby infer whether or not the topics are clustering cells by tissue type. For each topic, we computed the mean expression distribution across tissues for the genes associated with the top 250 most topic-specific peaks and then calculated the log_2_-ratio of that to the mean expression distribution of 250 randomly selected genes. Our results show that these topic-specific peaks are near genes with tissue-specific expression patterns that suggest specific cell types (Fig. 5, S9). In fact, the genes associated with many of the topics show evidence of enrichment for tissue subtypes, including distinct kinds of neurons and hypodermis, and even some small but clearly distinct cell populations, such as sex myoblasts. At this resolution, there appears to be no distinction between body wall muscle and intestinal/rectal muscle (topic 40 encompasses both), and it is not possible in all cases to assign a subtype to topics enriched in neuronal genes (for example, in topic 38). This limited cell type resolution could be due to instances of differing patterns of accessibility marking alternative promoters for the same set of genes that are expressed in many neuron subtypes (i.e. a common neuronal program); pan-neuronal, non-productive accessibility near genes that ultimately are expressed in only a specific subset of neurons; or noise in the LDA resulting in poor separation of cells belonging to neuronal subtypes.

**Figure 5:**
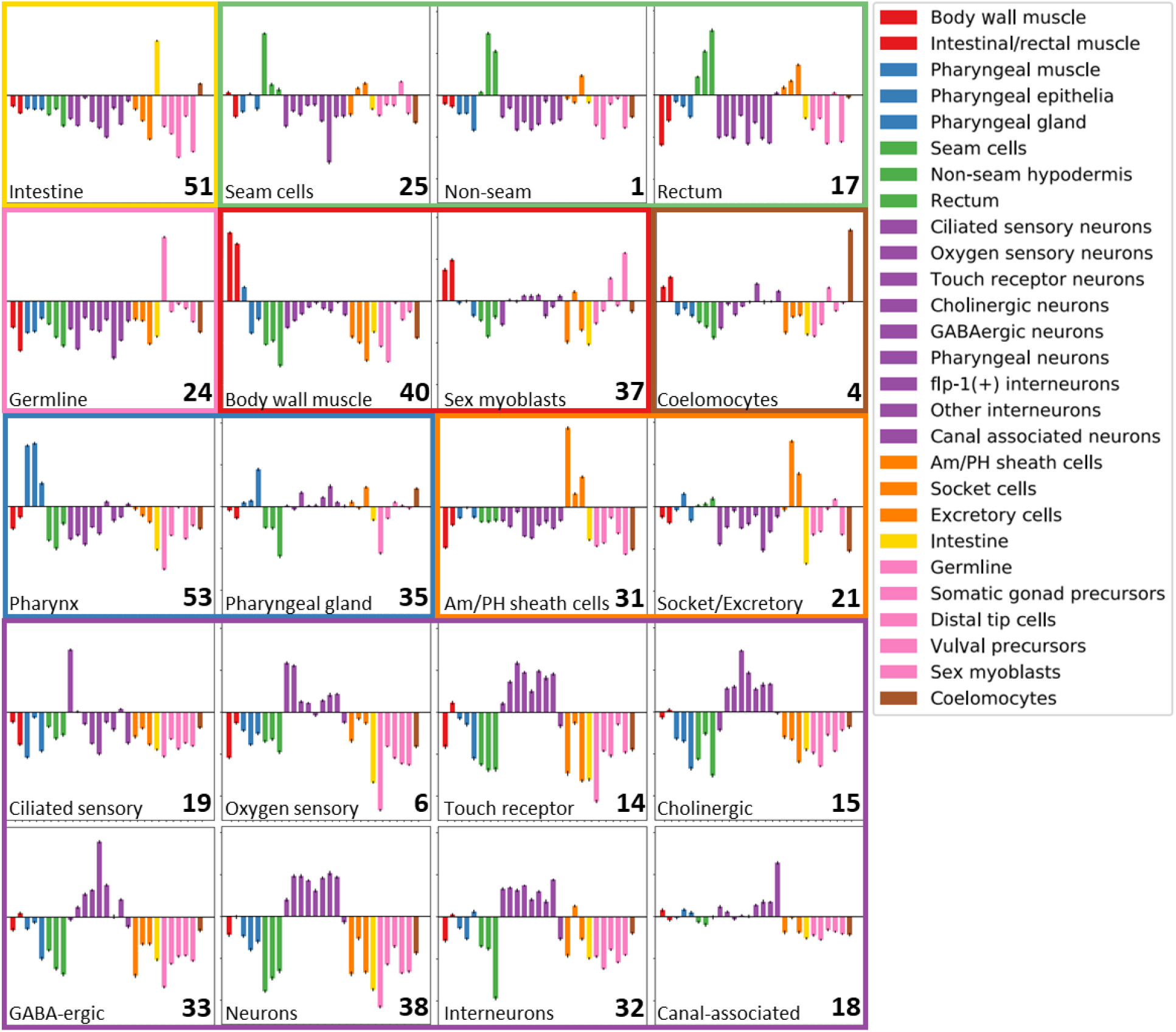
Topic-specific peaks tend to be near tissue-specific genes. Peaks associated with each topic were mapped to the nearest downstream gene, and the tissue expression distribution of the genes near the top 250 most-specific peaks for each topic was compared with the tissue expression distribution of 250 randomly-selected genes. Here we plot the results as the log_2_ ratio of the topic-associated tissue expression distribution to that of randomly selected genes. Error bars represent the 95% confidence interval after comparing to the tissue expression distribution of 100 random samples. LDA topic numbers are shown in the bottom right corner of each plot. Topics with similar tissue specificity patterns are grouped together, and the tissue type names and colors are as in Cao, et al. 2017. Tissue assignments were made by eye based on the tissue with maximal fold-change, and are written in the bottom left corner of each plot. If there was no single tissue was clearly the maximum then a more general tissue annotation was chosen (e.g. “Neurons” for topic 38). These annotations may need to be revisited with new data. Note that for concise visualization in this figure we display just 20 of our 37 topics, but we report a version of this figure with all 37 topics in Fig. S9.

We also investigated the sci-ATAC-seq signal at known tissue-specific genes, and we find strong tissue-specific chromatin accessibility that is consistent with the known expression patterns of these genes (Fig. 6). The genes *hlh-1, pha-4* (a master regulator of pharyngeal tissue), *elt-1, col-160* (a collagen gene that is expressed in non-seam hypodermis in L2), *bbs-8* (a gene encoding a receptor expressed in ciliated sensory and oxygen sensory neurons), *unc-47* (a gene expressed in GABA-ergic neurons), *elt-2*, T02B11.3 (a gene that is specifically expressed in sheath glial cells), and *glh-1* (a gene expressed specifically in the germline), all show accessibility enriched in the expected tissue types. The data also suggest patterns of differential isoform expression; for example, the 5’ end of the long *pha-4* isoform is most accessible in intestine cells, the 5’ end of a medium isoform has almost no accessibility, and the shortest isoform has pharyngeal accessibility. There are also several sites downstream of the *pha-4* gene that are strongly accessible in pharynx and perhaps represent other sites that play a role in regulation of this locus.

**Figure 6:**
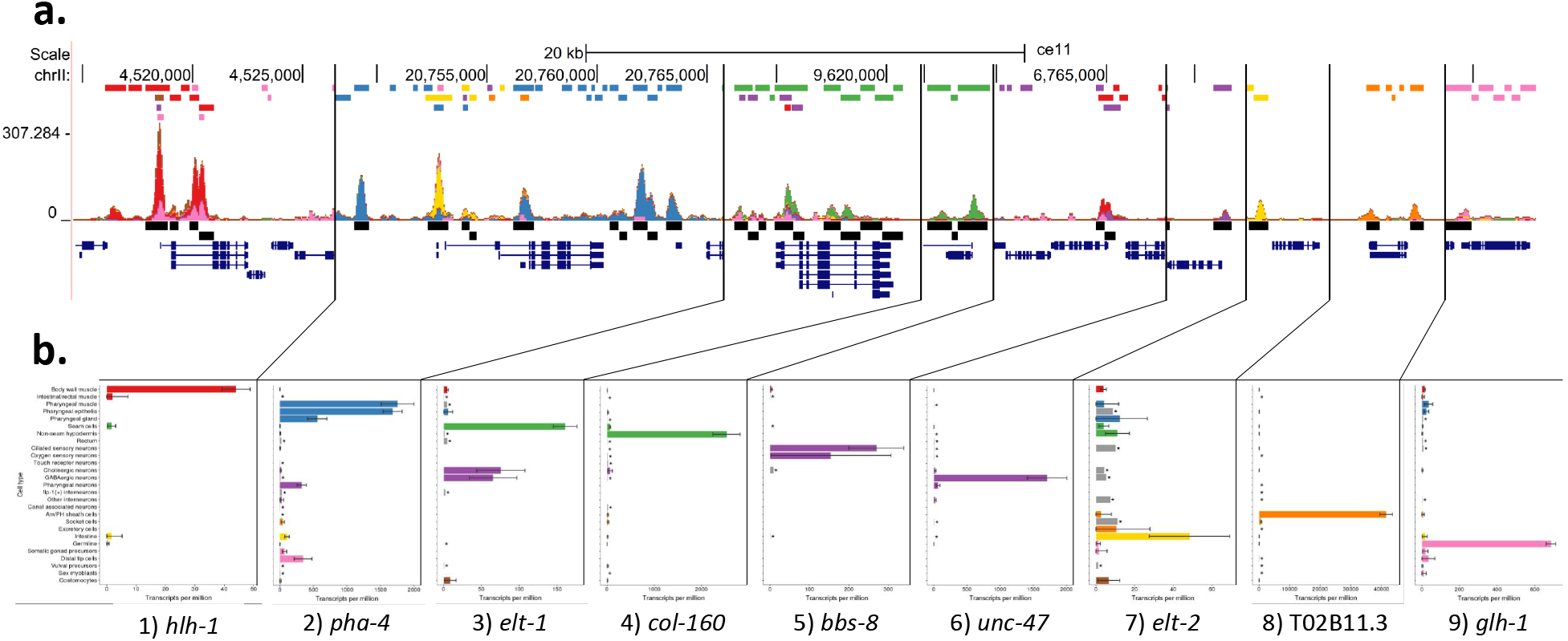
Known tissue-specific genes show topic-specific chromatin accessibility. UCSC Genome Browser multi-locus view of the regions surrounding nine known tissue-specific genes (**a.**), and the tissue expression patterns from scRNA-seq (**b.**). In the genome browser view, the top track shows the locations of sci-ATAC-seq peaks colored by tissue type, the second track shows the stacked sci-ATAC-seq signal from each tissue, the third track shows consensus peak regions around local maxima in the signal track, and the fourth track shows the gene models.

After identifying cell types associated with our topics, we revisited the list of peaks that had no overlap with TF ChIP-seq sites and looked for differences between those with overlaps and those without (Fig. 7). In general, most peaks contribute to primarily one topic with appreciable contributions to a few additional topics. The peaks that overlap TF ChIP-seq sites (Fig. 7a) tend to be found in many more cells in the L2 animal than the peaks without TF ChIP-seq overlaps (Fig. 7b). Nevertheless, the peaks without ChIP-seq overlaps still show clear topic-specificity, suggesting that they are informative peaks. In particular, over half of the peaks with no ChIP-seq overlaps contribute to neuron or gonad topics. Neurons are the most diverse tissue type in the worm, with scRNA-seq capable of identifying transcriptional signatures consistent with specific neuron cells (Packer et al., 2019); their absence in the ChIP-seq data suggests that whole-worm ChIP-seq lacks the sensitivity to find the cell-type-specific regulatory sites present in only a few L2 cells. On the other hand, transcription factors for gonad tissues are not well-characterized by the current modERN ChIP-seq compendium. Thus, we conclude that the sci-ATAC-seq peaks that do not overlap TF ChIP-seq sites are most likely either highly cell-type-specific, or specific to transcription factors that have not yet been tested with ChIP-seq.

**Figure 7:**
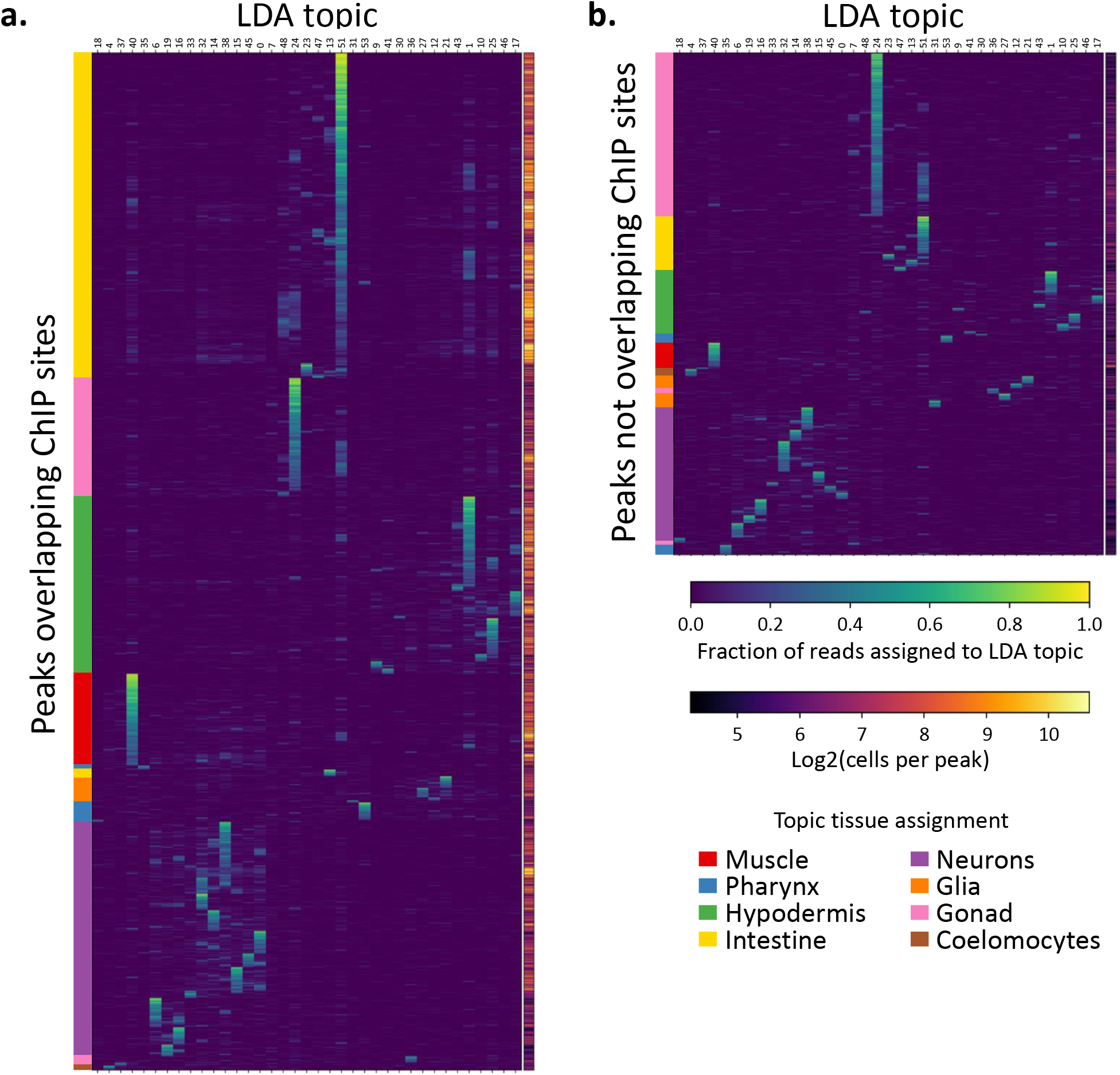
Novel sites of accessible chromatin with no overlapping modERN ChIP-seq peaks exhibit convincing topic patterns. We compare the normalized peak-by-topic matrix values between the peaks that overlap a ChIP-seq site (**a.**) and those that do not (**b.**). The nonoverlapping peaks are enriched for topics associated with gonad (especially germline/topic 24) and topics associated with neurons. The non-overlapping peaks also tend to be observed in fewer cells.

The proportion of cells assigned to the different tissue types generally aligns with the number of cells of each tissue type per worm, except for hypodermis and intestine. To understand this, it is important to remember that sci-ATAC-seq assays nuclei, not cells per se. We find 6,376 hypodermal nuclei, about ~ 26.1% of the total (Table 1), even though hypodermal cells make up far less than that percentage in the worm. Development of the hypodermis is a complex process that results in multiple large syncytial cells (Altun and Hall, 2002), and thus the number of nuclei is far larger than the number of cells. In addition, some of these nuclei are tetraploid, and we expect that nuclei with more copies of the genome will have more opportunities for the ATAC Tn5 enzyme to cut at a given genomic locus, yielding more data per cell. Similarly, although there are only 20 intestinal cells in the L2 worm (~ 2.7%) we recover 4,969 intestinal nuclei (~ 20.2%). As with hypdermis, some of intestinal cells are multinucleate and all of them are tetraploid (McGhee, 2007). Simply having more nuclei per cell will result in an over-representation of intestine and hypodermis in our data, but the polyploid nature of these tissues contributes by providing more Tn5 cutting opportunities as well for two reasons: first, more cut sites produce more reads and nuclei with more reads are more likely to pass the threshold for having enough reads to be analyzed (Fig. S1); and second, more cut sites are likely to result in more complex libraries in those cells, which will then be more confidently clustered and more likely to be assigned to a tissue type.

### 2.5 LDA modeling of cells from individual tissue types detects fine-grained cell types

While the topics we identified can distinguish cells at the level of tissue type, to yield more specific cell identities we tried a more focused analysis of cells from a particular cluster/tissue. We re-ran our LDA-based clustering procedure (Fig. 1) for cells of each tissue type to identify subclusters that correspond to more fine-grained cell types. We grouped the topics into eight major tissue types (coelomocyte, glia, gonad, hypodermis, intestine, muscle, neuron, and pharynx), and iteratively trained LDA models for each tissue, clustered the cells, and called peaks for each cluster. Similarly to the whole worm analysis, we provide the results as a UCSC Genome Browser track hub, viewable here: http://genome.ucsc.edu/cgi-bin/hgTracks?db=ce11&hubClear=http://waterston.gs.washington.edu/atacCellType/Durham_hub.txt.

Body wall muscle cells are quite similar to each other, despite differentiating from four different embryonic lineages. In previous single cell RNA-seq studies, the body wall muscle cells all clustered together, without much separation (Cao et al., 2017; Packer et al., 2019). Nevertheless, within the body wall muscle cluster the cells were found to group by anatomical position, setting up an anterior-posterior axis through the cluster that was identified by looking for the expression of specific marker genes. Similarly, the intestine cells showed evidence of expression differences in sci-RNA-seq by anatomical position. We looked for a similar anatomical pattern in the sci-ATAC-seq muscle and intestinal cell clusters(Figs. 8, S10, S11). We found cells with peaks associated with marker genes that are expressed throughout muscle (*hlh-1* and *myo-3*) and intestine (*end-1* and *elt-2*) distributed throughout the clusters (Fig. 8a), whereas those cells with peaks associated with marker genes with anatomically-biased expression showed distinct patterns (Fig. 8b). The subclustering thus appears to reveal still finer distinctions between cells.

**Figure 8:**
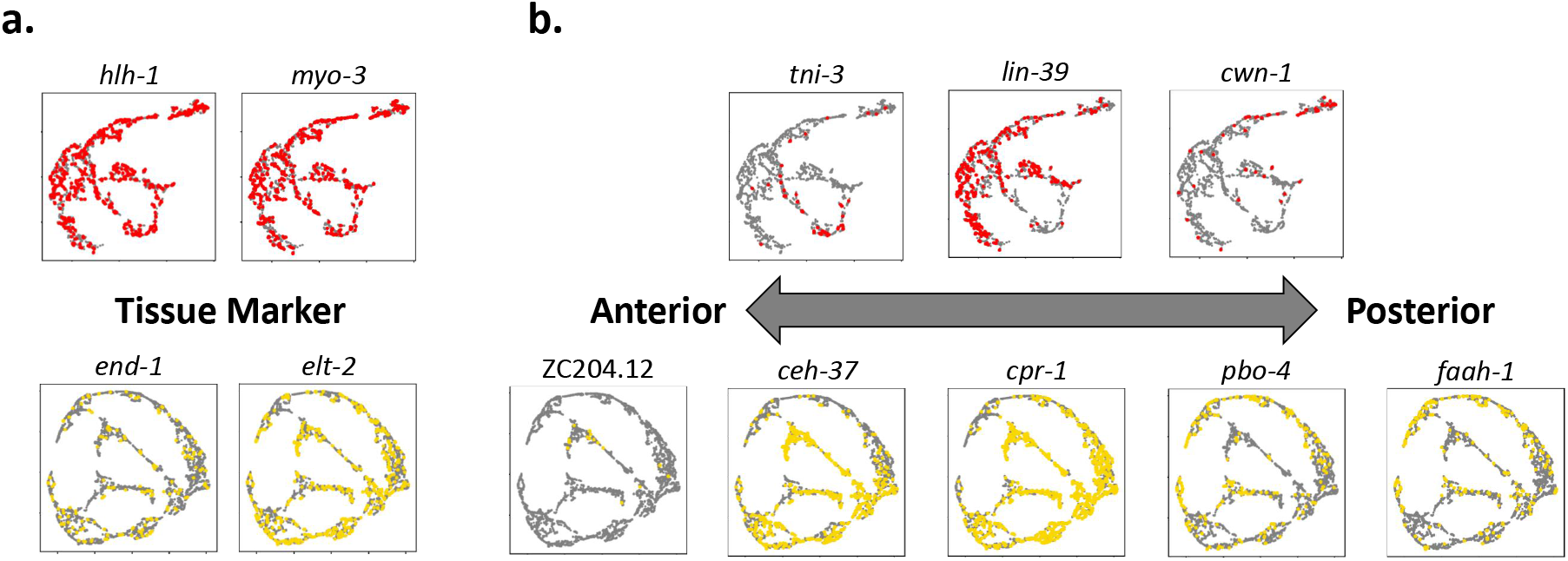
Subclustering of muscle and intestinal cells separates them by position along the anterior-posterior body axis. **a.** Peaks near genes that should be expressed throughout a tissue, like *hlh-1* and *myo-3* in body wall muscle or *end-1* and *elt-2* in intestine, show accessibility in cells throughout the UMAP. **b.** In both muscle and intestine data, we can detect subclusters of cells that show peaks near genes that mark the anterior or posterior regions of these tissues based on literature and microscopy data (Packer et al., 2019).

Next, we performed a similar analysis on all cells from neuron-enriched clusters. While the tissue type categories based on the L2 scRNA-seq data (Cao et al., 2017) already break gene expression distributions down into nine different neuronal subtypes, there are many more specific neuron types revealed by anatomical analysis and by additional scRNA-seq data sets (Packer et al., 2019; Taylor et al., 2019). (Fig. 5). In order to take a closer look at the neuron cells, we gathered all cells from topic clusters showing neuronal enrichment and performed the same analysis that we did for body wall muscle and intestine above. The neuron LDA model yielded 36 topic clusters – only one fewer than the number of clusters found by the whole worm model, highlighting the diversity of neuronal cell types. We evaluated five marker genes chosen for their tissue specific expression patterns based on the scRNA-seq data: *bbs-8* is expressed in ciliated sensory neurons; *gcy-32* is expressed in oxygen sensory neurons; *unc-30* is expressed in GABA-ergic neurons; *mec-7* is expressed in touch sensitive neurons; and *ceh-24* is expressed in cholinergic neurons. We find that the cells with chromatin accessibility near these genes are associated with distinct, well-separated clusters in UMAP space (Fig. 9a), suggesting cell type identities for these clusters.

**Figure 9:**
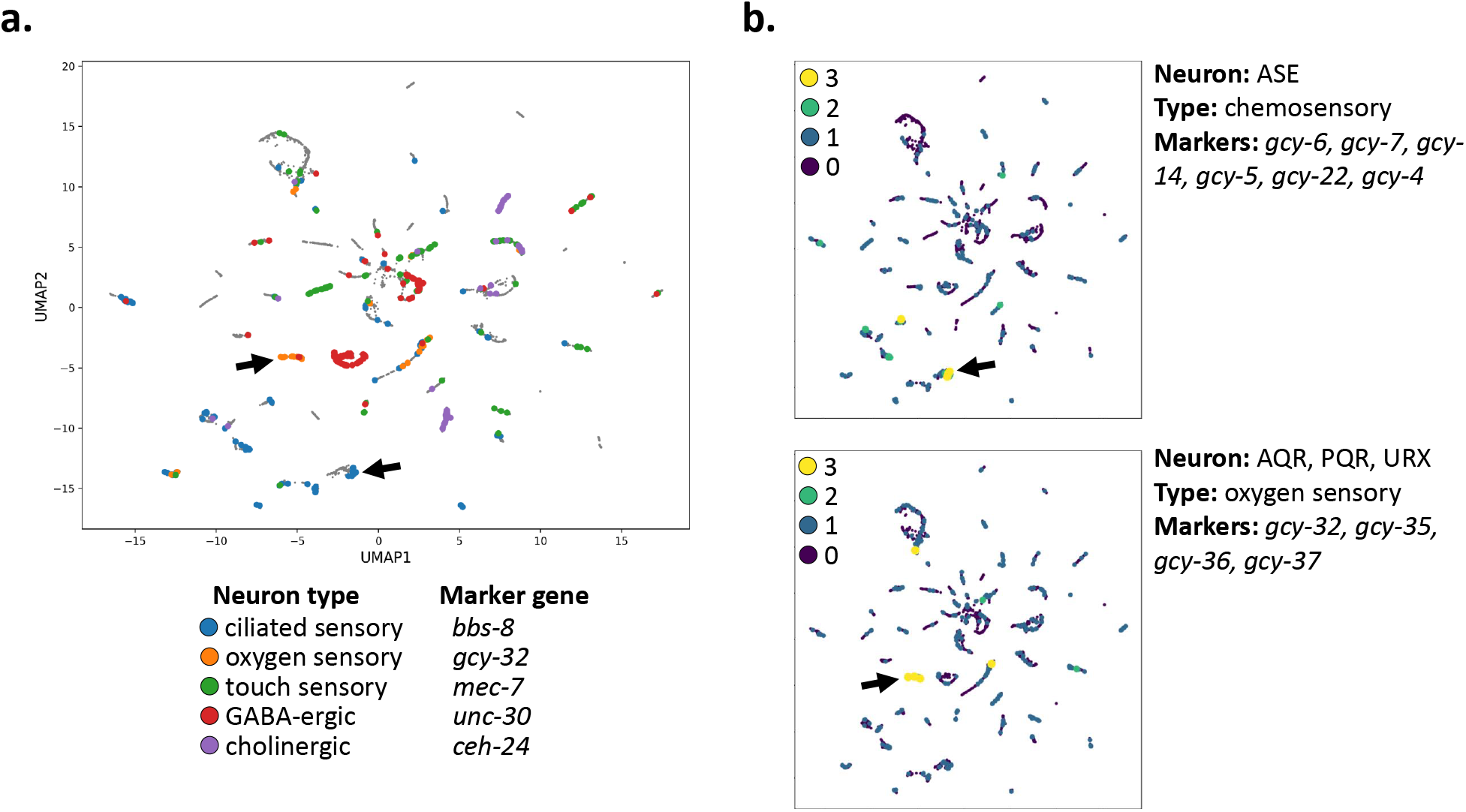
Subclustering of neurons reveals finer structure that distinguishes different types of neurons. **a.** Cells with reads in peaks near genes with expression patterns specific to neuron subtypes cluster together (*bbs-8*: ciliated sensory neurons, *gcy-32*: oxygen sensory neurons, *unc-30*: GABA-ergic neurons, *mec-7*: touch receptor neurons, *ceh-24*: cholinergic neurons). **b.** Cells in the UMAP plot are colored by the number of marker genes with nearby co-accessible peaks. Here, we show marker genes for the ASE neurons, a specific pair of ciliated sensory neurons, which are identified in one of the *bbs-8* clusters from **a.** (marked by the left-facing arrow); and marker genes shared by the oxygen sensory neurons AQR, PQR, and URX, which further support the cluster marked with *gcy-32* in **a.** (marked by the right-facing arrow).

In order to verify the neuron subtypes identified by single marker genes and also attempt to identify additional subtypes for the other clusters of cells, we followed up by checking additional marker genes for specific neurons (Fig. 9b). The ASE neurons are a pair of ciliated chemosensory neurons that detect water-soluble attractants like potassium and sodium ions. They express *bbs-8*, along with a highly specific repertoire of guanylyl cyclase (*gcy*) genes that encode sensory receptors. We found chromatin accessibility peaks near six such receptor genes specific to ASE neurons: *gcy-4, gcy-5, gcy-6, gcy-7, gcy-14*, and *gcy-22* (Etchberger et al., 2009). In order to identify the ASE neurons in the UMAP, we colored cells by the number of co-accessible peaks they have that are near the ASE-specific *gcy* genes, which identifies one of the *bbs-8* subclusters as likely to be ASE neurons. We performed a similar analysis to confirm the UMAP cluster containing the AQR, PQR, and URX oxygen-sensory neurons. Oxygen-sensory neurons express their own repertoire of *gcy* genes, including *gcy-32, gcy-35, gcy-36*, and *gcy-37* (Packer et al., 2019), and peaks near these genes are co-accessible primarily in the cells marked by *gcy-32* in Fig. 9a. We identified additional examples of neuron subtypes, including mechanosensory neurons, two types of motor neurons, and interneurons (Fig. S16). We show the neuron UMAP colored by LDA topic probabilities in Supplementary Figures S12, S13, S14, and S15. Finally, we also subclustered coelomocyte (Figs. S17, S18), glia(Figs. S19, S20), gonad (Figs. S21, S22), hypodermis (Figs. S23, S24), and pharynx (Figs. S25, S26), finding evidence for additional specific cell types in each of them. Thus, sci-ATAC-seq can make fine-grained distinctions among cell types at a resolution that approaches scRNA-seq and thereby begin to associate specific regions of accessible chromatin with specific cell types.

## 3 Discussion

We used the sci-ATAC-seq assay to assemble the first cell type-resolved map of regulatory elements in *C. elegans*. We found 38,017 peaks, which we used to assign 24,503 of our 30,930 cells to one of 37 different clusters (Fig. 3) that represent distinct, differentiated tissues in the L2 nematode (Fig. 5). Our map, derived from data collected in essentially a single experiment, recovers the vast majority of L2 regulatory sites detected by hundreds of individual ChIP-seq experiments (Fig. 2), and accessibility at these sites can distinguish among cell types with small populations in the worm, such as specific types of muscle, intestine, and neurons (Figs. 5, 8, 9).

In addition to producing tissue-specific maps of regulatory DNA in L2, another significant contribution of our work is our implementation of the latent Dirichlet allocation code. LDA is a modeling approach that handles sparse data very well, and has previously been successfully applied to single cell genomics data (González-Blas et al., 2019; Kim et al., 2019; Dey et al., 2017). However, the initial implemtation of cisTopic, which is perhaps the most high-profile LDA implementation in the field, trains each model on a single CPU and is too slow to be practical for exploring data sets with tens of thousands of cells. Furthermore, its training procedure effectively throws out information from all training iterations except the last one. The cisTopic developers have just recently released an updated version of their code that addresses some of these issues, but in the meantime we wrote our own implementation of LDA that parallelizes the model training across multiple CPUs to vastly increase training speed. We also added the option to make a point estimate of the posterior distribution by taking the mean or mode of the model parameters across all training iterations after model burn-in instead of just reporting the parameters from the final training iteration. The third key feature of our implementation is a principled cross validation procedure for choosing the best number of topics to use that helps to avoid over-fitting the model (see Methods). Our implementation is written in Java, making for easy installation with no dependencies aside from Java itself, and can train a model on our full data set with 30,000 cells in about two hours on a machine with 32 GB of RAM and 8 cores.

The cell type resolution of sci-ATAC-seq data approaches that of scRNA-seq (Cao et al., 2017; Packer et al., 2019), despite the extreme sparsity and low dynamic range of the data in individual cells. Single cell ATAC-seq suffers generally from sparsity and a low dynamic range because there are at most two chances in a diploid cell to sample a given accessible locus, depending on whether or not both alleles are accessible. This contrasts to single cell RNA-seq, which has hundreds or even thousands of chances for measuring the mRNA of highly expressed genes. The sparsity of scATAC-seq was made more extreme in our data set because the worm samples yielded sci-ATAC-seq libraries with many fewer unique fragments per cell than other organisms. In the first reported sci-ATAC-seq results, human and mouse cell lines yielded a median of 2,503 fragments per cell (Cusanovich et al., 2015), and in more recent work on fly and mouse cells the median yield was over 10,000 fragments per cell (Cusanovich et al., 2018a,b). In contrast, despite using the most up-to-date protocol (Cusanovich et al., 2018a,b), the median *C. elegans* cell yielded only about 700 fragments (Fig. S1). We hypothesize that this could be improved by optimizing the nuclear isolation and permeabilization conditions — L2 nuclei are extremely small, compact, and dense (about 2μm in diameter), and possibly after formaldehyde fixation Tn5 has only limited access to the chromatin. As single cell technology advances and we are able to generate more complex libraries we expect that scATAC-seq will provide even higher resolution of individual cell types in the worm.

We also expect to see improved results from applying more advanced computational techniques, especially new approaches that more tightly integrate the analysis of the sci-ATAC-seq and scRNA-seq data sets. There now exist multiple approaches for projecting single cell data from different modalities into the same embedding space (Cusanovich et al., 2018a; Welch et al., 2017; Stuart et al., 2018). By jointly analyzing the sci-ATAC-seq data and sci-RNA-seq data with one of these methods it may be possible to improve the cell type resolution of our chromatin accessibility maps.

Such accessibility maps with high cell type resolution will be important for understanding gene regulation on the scale of the whole genome across the whole organism. In addition, regulatory sites are hypothesized to play a major role in common disease and evolutionary adaptation, so maps of regulatory sites will aid in interpreting the effects of genetic variation. For example, many mutations that are linked to some phenotype by approaches like GWAS do not fall in genes. The implication is that, if one of the mutations is indeed causal, it must fall in a regulatory sequence of DNA. Thus, maps of cell type-specific regulatory regions can help interpret and prioritize candidate causal variants, and will be a useful complement to genetic resources in *C. elegans*, including the *C. elegans* Natural Diversity Resource (Cook et al., 2017) and the Million Mutations Project strains (Thompson et al., 2013). Better annotations of regulatory DNA can also help with understanding comparative genomics and the evolution or conservation of stretches of non-coding DNA.

Given the importance of mapping regulatory sites for understanding genome structure and function, and the power of *C. elegans* as a model organism, improving and expanding our maps of regulatory regions should be a high priority. In particular, collecting accessibility data for additional developmental stages in worm will provide valuable insight into the dynamics of gene regulation over the course of development as cells differentiate. These data can be paired with new scRNA-seq data collected from throughout *C. elegans* embryogenesis (Packer et al., 2019), moving the field closer to having a truly comprehensive map of gene expression and regulation for every cell throughout development in *C. elegans*.

## 4 Methods

### 4.1 Nuclear isolation from whole L2 worms

We grew wild-type *Caenorhabditis elegans* worms (VC2010 strain) at 21°C on nine 150 mm plates and synchronized the population by bleaching (2% bleach, 0.5 M KOH) young adults with 8-12 embryos to isolate embryos, hatching them at room temperature in egg buffer (118 mM NaCl, 48 mM KCl, 2 mM CaCl_2_, 2mM MgCl_2_, HEPES 25 mM at pH 7.3) for 12-16 hours, and re-plating the L1 hatchlings onto nine more 150 mm plates at a density of approximately 60,000 worms per plate. After two rounds of this bleach synchronization and plating, the L1 worms were allowed to grow at 21°C for 19 hours after plating to reach the middle of the L2 stage. The worms were washed off eight of the plates with M9 buffer (22 mM KH_2_PO_4_, 22 mM Na_2_HPO_4_, 85 mM NaCl, 1 mM MgSO_4_ at pH 6.5) into a 50 ml conical tube. Bacteria were removed from the suspension by spinning the tube at ~ 3,000 × *g*, aspirating the supernatant, resuspending in fresh M9. The M9 wash was repeated, and the supernatant was aspirated, leaving a worm pellet in ~ 1ml of M9. The worm pellet was flash frozen by using a P1000 to transfer the worms drop by drop into a mortar containing liquid nitrogen. The frozen worms were crushed into powder with a pestle such that each worm broke into 3-4 chunks, and the powder was transferred to a 50 ml falcon tube containing 8.75 ml of 1.1% formaldehyde in egg buffer supplemented with 1x protease inhibitor. Worms were rocked at room temperature for 10 min before the fixation reaction was quenched by adding 1.25 ml 1M glycine (final concentration ~ 125mM) and incubated another 5 min at room temperature. The fixed worms were pelleted at 3220 × *g* for 5 min at 4°C, the supernatant was removed, and the pellet was resuspended in 10ml ice cold egg buffer. Fixed worms were pelleted again by spinning at 3220 × *g* for 5 min at 4°C. The egg buffer supernatant was aspirated, and the pellet was resuspended in ice cold 2x nuclear preparation buffer (20 mM HEPES pH 7.6, 20 mM KCl, 3 mM MgCl_2_, 2 mM EGTA, 0.5 M sucrose, 0.05% Triton X-100 in egg buffer) supplemented with protease inhibitor (NPB+PI). The following steps were all performed at 4°C or on ice: The solution was transferred to a 7 ml Dounce homogenizer and the fixed worm chunks were homogenized with 20 loose pestle strokes followed by ten tight pestle strokes. The Dounced suspension was spun for 90 seconds at ~ 200 × *g* in a swing-arm centrifuge to loosely pellet debris, and the top 1000 *μl* of supernatant (containing the nuclei) was removed to a 15 ml falcon tube on ice. 1 ml of fresh NPB+PI was added to the Dounce, the debris pellet was gently resuspended, and the Douncing and spinning were repeated three more times, resulting in the collection of 4 ml of nuclei. The suspension of nuclei was cleaned by gently passing through a 10μm syringe filter pre-wetted and chased with 1 ml ice cold NPB+PI into a new 15 ml falcon on ice. The nuclei were split evenly into 1.5 ml eppendorf tubes and pelleted at 2000 × *g* for 10 min at 4°C. All supernatant was removed, and the pellets were each gently resuspended in 1 ml freezing solution (50 mM Tris at pH 8.0, 25% glycerol, 5 mM Mg(OAc)_2_, 0.1 mM EDTA, 5 mM DTT, 1× protease inhibitor cocktail (Roche), 1:2,500 superasin (Ambion)) (Cusanovich et al., 2018b). The resuspended nuclei were transferred to 2 ml cryotubes, flash frozen in liquid nitrogen, and stored at −80°C.

### 4.2 Single cell ATAC-seq via single cell combinatorial indexing

The sci-ATAC-seq protocol was as described Cusanovich, et al. 2018 (Cusanovich et al., 2018b). Briefly, flash-frozen VC2010 nuclei were thawed in a 37°C water bath and put immediately on ice. The nuclei were transferred to a 1.5 ml eppendorf tube and spun at 2000 × *g* for 10 min. The supernatant was aspirated, and the pellet was resuspended in 200 μl of ATAC-OMNI (Corces et al., 2017) RSB (10 mM Tris-HCl pH 7.4, 10 mM NaCl, and 3 mM MgCl_2_ in water) supplemented with 0.01% Digitonin, 0.1% IGEPAL-630, and 0.1% Tween-20, allowed to stand for 3 min on ice, and then quenched by adding 1 ml of RSB supplemented with 0.1% Tween-20. The resuspended and lysed nuclei were stained with 1x Hoechst and a BD FACS Aria II was used to distribute 2500 nuclei into each well of a 96-well v-bottom plate (Eppendorf twin.tec LoBind skirted 96 well PCR plate) prepared with 19μl of tagmentation reaction solution (10μl 2x Nextera TD buffer, 3.3μl 1X DPBS, 0.2μl 1% Digitonin, 0.2μl 10% Tween-20, 5.3μl H_2_O) (Corces et al., 2017). After sorting, 1.0 μl of 2.5 μM uniquely-barcoded Tn5 from Illumina (Cusanovich et al., 2015) was pipetted into each well of the 96 well plate, and the transposition reaction was allowed to proceed at 55°C for 30 minutes. Next, 20 μl of STOP reaction buffer (40 mM EDTA and 1 mM Spermidine) was added to quench the reaction, and the plate was put at 37°C for 15 min. After stopping transposition, all nuclei were pooled into a 15 ml conical tube, re-stained with 1x Hoechst, and distributed by FACS into twenty eight 96-well v-bottom plates at 25 nuclei per well. The 96-well plates contained 12μl per well of reverse cross-linking buffer (0.83 mg/ml Proteinase K and 0.042% SDS in Qiagen EB buffer), and were put on ice, spun down, and frozen at −20°C in batches during sorting. Later, these plates were thawed in groups of four for reversing crosslinks by incubating at 65°C for 16 hours, after which the transposed and un-crosslinked fragments were amplified using uniquely-barcoded PCR primers. We ran four wells as test reactions in qPCR and monitored the libraries for saturation of SYBR-Green signal to identify the number of cycles required for appropriate amplification (Cusanovich et al., 2018b), and then amplified the rest of the wells for either 22 cycles with Illumina NPM 2x PCR master mix or 23 cycles with NEBNext 2x PCR master mix (PCR Reaction: 12.0 μl of nuclei in reverse crosslinking buffer; 2.5 μl of 5 μM Nextera v2 barcoded P7 PCR primer; 2.5 μl of 5 μM Nextera v2 barcoded P5 PCR primer; 1.0 μl of 100X BSA; 25.0 μl of 2x NEBNext PCR Master Mix (NEB cat M0541); and 7.0 μl nuclease free H_2_O) (NPM PCR protocol: 72°C for 3:00, 98°C for 0:30, repeat×22(98°C for 0:10, 63°C for 0:30, 72°C for 1:00), 4°C HOLD; NEBNext PCR protocol: 72°C for 5:00, 98°C for 0:30, repeat×23(98°C for 0:10, 63°C for 0:30, 72°C for 1:00), 4°C HOLD). After amplification, the fragments were cleaned up by pooling the contents of all wells and splitting across four Zymo Clean and Concentrate columns (cat. D4014), eluted each in 25 μl Qiagen EB, combined the eluates, and then further cleaned and concentrated with 1x Ampure XP magnetic beads, and finally eluted in 25 μl. Library quality was assessed using the Agilent TapeStation D5000 kit (Screentape cat. 5067-5588, and reagents cat. 5067-5589), and molarity was quantified for fragments between 200 and 1000 bp. Last, the libraries were combined into an equimolar pool at 2 nM for sequencing. Libraries were sequenced using the manufacturer’s denaturation conditions, and loaded either on a Illumina MiSeq 300 cycle v2 kit (cat. MS-102-2002) at an input concentration of 15 pM, or on a Illumina Nextseq Mid-Output 300 cycle v2.5 kit (cat. 20024905) at an input concentration of 1.8 pM, and using custom sequencing primers and recipe from Illumina.

### 4.3 Generation of genomic DNA input control

In order to control for the sequence cutting bias of Tn5 (Green et al., 2012), we treated naked *C. elegans* genomic DNA with the bulk ATAC-seq protocol (Buenrostro et al., 2013). We isolated genomic DNA with phenol:chloroform extraction and ethanol precipitation. In order to keep the Tn5:DNA ratio similar to a bulk ATAC-seq experiment with 50,000 cells, we estimated that a typical *C. elegans* nucleus will contain 1e^6^bp×2genomes×660MW/bp× 1.67e^-12^pg/MW ≈ 0.22pg/nucleus, or ~ *11ng* in 50,000 nuclei. We diluted the DNA to a concentration of ~ 0.87 ng/μl, as measured with the Qubit High Sensitivity assay (Invitrogen), and used 11.5 μl as input to a 25 μl reaction with 12.5 μl of Nextera TD buffer and 1.0 μl of Nextera Tn5 enzyme. The reaction was incubated at 37°C for 30 min, cleaned up with a Qiagen MinElute column, and amplified using NEBNext 2x PCR Mix with primers from Buenrostro, et al. 2013 (Buenrostro et al., 2013). The libraries were cleaned up with 1:1 AMPure XP magnetic beads, and sequenced on the Illumina NextSeq platform.

### 4.4 ATAC-seq alignment pipeline

Initial processing of the sequencing results was done as reported in Cusanovich, et al. 2018 (Cusanovich et al., 2018b), with some changes. Sequencing results were converted to FASTQ format with the Illumina `bcl2fastq` program (v. 2.19). First, the integrity of the barcode sequences was checked for each of the four components of the barcode (tagmentation barcodes from both sides of the cut and the P5 and P7 primer indices added during PCR amplification) by matching the sequencing results to the known barcode sequences. Any read that had three or fewer edits compared to the best-matching known barcode sequence and that had no other known barcode sequences matching with five or fewer edits were corrected and assigned to the best-matching barcode sequence. Any read-through of short templates was corrected by trimming adapter sequences from reads using Trimmomatic (Bolger et al., 2014) (v0.36) with the options `ILLUMINACLIP:NexteraPE-PE:2:30:l0:l:true, TRAILING:3, SLIDINGWINDOW:4:10`, and `MINLEN:20`. Next, read sequences were aligned to the WS235/ce11 build of the *C. elegans* genome with bowtie2 (Langmead and Salzberg, 2012) with options `-X 2000` and `−3 1`, properly paired reads with mapping scores greater than 10 were kept, any reads mapping to the mitochondrial DNA were filtered out, and read pairs with identical barcode sequence and identical starting and ending mapping coordinates were identified as PCR duplicates and collapsed to 1 using a custom script (Cusanovich et al., 2018b). Next, read coverage for each cell was calculated, and cell barcodes with fewer than 150 reads were removed from further analysis (Fig. S1). The reads that made it through filtering for each batch of sequencing were merged into a single BAM file using the Picard MergeSamFiles program (http://broadinstitute.github.io/picard). Last, the reads in the merged BAM file were converted into cut sites by taking 60 bp intervals centered on the fragment ends, shifting those sites for reads mapping to the forward strand by +4 bp and the negative strand by −5 bp (to account for the shape of the Tn5 cut site (Buenrostro et al., 2013)), and writing the resulting coordinates to a BED file for peak calling.

We called peaks from the cut site data using MACS2 (Zhang et al., 2008) (v. 2.2.5) with options `--format=BED, -g 9e7, --nomodel, --qvalue=0.05, --SPMR, --tsize=60, --bdg, --keep-dup all`, and `--call-summits`. Additionally, we aligned and mapped cut sites for some bulk ATAC-seq data collected on naked *C. elegans* genomic DNA and provided this as an input control to correct our peak calls for sequence bias in Tn5 cutting. After peak calling, we merged any overlapping peaks to produce a single set of non-overlapping genome-wide peak calls. Finally, we generated a binary cells-by-peaks matrix that records which peaks were detected (i.e. were overlapped by a cut site) in each cell. This data structure was used for further cell clustering analysis with latent Dirichlet analysis (LDA), described below.

### 4.5 Cell clustering pipeline overview

In order to identify cell types from the sci-ATAC-seq data, we used an iterative clustering and peak calling approach. After investigating dimensional reduction methods including PCA and LSI, we found LDA provided the best separation between clusters of cells. The core pipeline consisted of three main steps: first, training a latent Dirichlet allocation model (LDA, see below) on the data; second, identifying LDA topics that corresponded to coherent groups of cells and using those topics to cluster the cells; and finally, calling peaks on pooled data for each cluster of cells. Calling peaks on the cell clusters increases sensitivity to detect cluster-specific peaks compared to the peak calling at the end of the alignment pipeline; cell type-specific regulatory sites may be weaker than more commonly-accessible peaks either because they are smaller or more transiently accessible, or because they may be specific to a small subset of the cells in the whole worm. Either way, the data for such sites might not rise above the background noise from other cell types in the whole worm data set, but are detectable in more homogeneous subsets of cells. We iterated this procedure twice: first, to generate a more sensitive set of peaks than the peak calls from the alignment pipeline (we call this first iteration the primary LDA), and a second time to train a new LDA model with the improved peak set, which also results in a third, refined peak set (we will refer to the second iteration as the refinement LDA). Last, MACS2 occasionally calls broad peaks that cover large regions of the genome of hundreds or even thousands of base pairs. The signal over these peaks is usually multimodal with multiple distinct “summits” that most likely represent distinct binding sites. In order to better capture the distinct nature of these summit regions in the refined peak set, we implemented a custom script that identifies the local summits within a MACS2 peak and splits the peak into multiple contiguous segments that each encompass a single summit region, and we report these split peaks in our UCSC Genome Browser track hubs. (Note that our peak-splitting procedure is similar to, but distinct from, the MACS2 summit-calling procedure, which reports the coordinate of the base pair with the highest signal and does not actually segment the peak around the summits.) For the peak-splitting code, see the `expand_summits2.py` script in the paper’s GitHub repository: https://github.com/tdurham86/L2_sci-ATAC-seq.

### 4.6 Latent Dirichlet allocation implementation

Inspired by the effectiveness of LDA as implemented in cisTopic (González-Blas et al., 2019), we decided to take this approach to analyzing the sci-ATAC-seq data. Briefly, LDA is a Bayesian modeling strategy that was originally developed in the setting of document classification. It assumes that each document is characterized by one or more latent “topics”, and that these topics are characterized by subsets of the words in the document. Documents are modeled as Dirichlet probability distributions over the set of topics, and the topics are modeled as Dirichlet probability distributions over the set of words. Training proceeds by iterating over the entire vocabulary defined by the documents it is modeling and proposing a topic for every instance of every word in every document. The probability of picking a topic for a given word and document is computed based on the current probability distribution of topics for that document and the probability of words for each topic. At the end of training, the probability distributions for the topics over the documents and the words over the topics can be calculated by summing the topic proposals for all peaks, and for all documents, respectively. Our implementation uses a collapsed Gibbs sampler to speed up training by sampling the latent parameters of the model from the full conditional posterior (Griffiths and Steyvers, 2004). When applied to single cell ATAC-seq data, as in cisTopic, cells are treated as the documents and peaks are treated as the words. The LDA model then learns topics that distinguish among the cells based on which peaks tend to be accessible in similar patterns across all cells, and outputs two matrices that capture the relationship between peaks and topics, and cells and topics: the first matrix contains the counts of the number of cells for which a given peak was assigned to each topic (the peaks-by-topics matrix), and the second matrix contains how many peaks from each cell were assigned to each topic (the cells-by-topics matrix).

We began by using cisTopic itself, but found the R implementation to take almost two days to process the full data set. (Note that a new recently released version of cisTopic is significantly faster and incorporates similar ideas to our implementation.) In order to speed up the modeling, we implemented a parallelized version in Java that can split the training of a single model across multiple cores, reducing the run time to just a couple of hours. In the end, for the whole worm primary LDA we used 34 topics and set the alpha parameter for the Dirichlet priors to 3.0 to concentrate the probability distributions into just a few peaks/topics. We set the beta parameter to 2000.0; the higher value allows the LDA model to spread the probability across more peaks. These alpha and beta values control the weight of the symmetric, or uniform, priors used for the cell-by-topic and topic-by-peak distributions – the higher the alpha or beta value, the more weight is given to the uniform prior, which encourages the model to spread probability across topics and peaks instead of concentrating it. The Java code can be accessed on GitHub at https://github.com/gevirl/LDA and is released under the GNU GPLv3 license.

### 4.7 Filtering the cells-by-peaks matrix for LDA

In order to improve the efficiency and effectiveness of LDA, it can help to remove cells that have very few peaks and to remove peaks that are either found in very few cells or too many cells. Sparse cells and peaks provide little useful information and are enriched for noise, and while peaks that are found in too many cells are likely real, they are not very helpful for distinguishing among cell types. We filtered the outliers by sorting the cells by the fraction of all peaks that was detected in each cell (or peaks by the fraction of all cells in which they were detected), mean centering the data, and then identifying the change points at the extremes of the resulting curve by doing a convolution with values from a sigmoid function to detect the areas where the slope is changing fastest. We set the filtering thresholds as four times the inter-quartile range above the mean of the convolution output (Figs. S2, S3). We applied LDA to the resulting filtered cells-by-peaks matrix.

### 4.8 Latent Dirichlet allocation model selection

Choosing hyperparameter values is one of the most challenging aspects of training models like LDA. In particular, using an appropriate number of topics is critical to getting good results, and picking this number requires an empirical approach. In cisTopic (González-Blas et al., 2019), the authors recommend training several model instances, each with a different number of topics, and choosing the number of topics that gives the best log likelihood of your input data. However, increasing the number of topics adds parameters to the model, which makes the model better able to fit the training data, even if it has already fit the true signal and begins to train on noise (i.e. it is overfitting). Since the cisTopic procedure uses the same data for training and evaluation, it does not test the generalizability of the model parameters, and cannot tell when the model starts to overfit. Ultimately, it will recommend using a higher number of topics than can be supported by the data. In order to identify a suitable number of topics that avoids overfitting, we implemented the following cross-validation procedure.

First, the cells are evenly and randomly split into five disjoint sets for five-fold cross validation. Then, for each number of topics that we would like to test, five LDA models are trained, with each model training on four of the folds and holding one out for evaluation. Once each model is done training, it estimates the likelihood of the data in the held-out test fold with a Chib-style estimator (Wallach et al., 2009). In this estimation procedure, the peak-topic probabilities learned from the training data are fixed, and then a cell-topic vector is trained for each held out cell based on the fixed peak-topic probabilities. The log likelihood of each held out cell is estimated based on sampling from the posterior of the model trained on that held out cell. We convert these log likelihoods to perplexity, which is defined as follows:

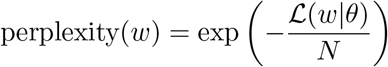

where *w* is a held out test cell, 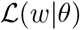 is the log likelihood of that test cell given the LDA model, and *N* is the number of peaks found in that cell. Because perplexity is inversely related to the log likelihood, smaller values are better. The best number of topics to use is the one that produces the lowest mean perplexity from the five held out sets of test data. It’s important to note that LDA is a stochastic modeling technique, and training on the same data with different random seeds will yield similar but different solutions. In addition, our sci-ATAC-seq data are by nature noisy and complex. Thus, the hyperparameter search procedure will not always result in a clear best number of topics to pick. We found that if we trained a model with a few extra topics beyond the optimal number, LDA would largely ignore the extra topics and still put almost all of the probability in a number of topics that approximated the underlying dimensionality of the data. Given that the model appeared robust to some extra topics, we ran our models with 1.5 times the number of topics recommended by the hyperparameter search (Figs. S4, S5).

### 4.9 Latent Dirichlet allocation training

To train the LDA model, we used our parallelized implementation of LDA, which could generally run on the full data set in about two hours on a machine with 8 cores and 32 GB of memory. Here is an example command line from the whole-worm refinement LDA analysis (see the GitHub repo for full documentation and usage information):

java -Xms32G -cp LatentDirichletAllocation.jar org.rhwlab.lda.cache.matrix. LDA_CommandLine -lda -a 3.0 -b 2000.0 scatac_data.bow -li 4000 -o ./out/dir -s 1 -t 55 -th 8 -tn 5 -ch 0 -rid 0000 -pe -d topic -st mode -sk 40 -v 1 -id ./out/dir/0000_topics55_alpha3.000_beta2000.000 -pr 1.0

The model has four main outputs: the docTopic matrix, which contains the raw counts of how many peaks were assigned to each topic in each cell; the theta matrix, which contains the probability distribution across topics for each cell and takes into account the full LDA probability, including the prior; the wordTopic matrix, which contains the raw counts of how many times each peak was assigned to each topic across all cells; and the phi matrix, which contains the probability distribution across peaks for each topic, and takes into account the prior. We used the theta matrix to cluster the cells, and the wordTopic matrix for identifying the cell type for each cluster.

### 4.10 Cell clustering by topics

Next, we sought to cluster the cells based on the LDA modeling results. We reasoned that differences among cell types would be the dominant source of informative variation in our sci-ATAC-seq data, and that, for this reason, many topics should mostly correspond to distinct cell types. In order to identify which topics were most likely to distinguish among cell types, we looked for topics that had high probability in subsets of cells that were close together in LDA topic-space and used these topics to define cell clusters as follows. In order to identify topics that corresponded to groups of cells in LDA topic-space, for each topic, we ranked the cells by their probability for that topic in the theta matrix, computed the centroid of the topic as the mean topic probability vector for the top 50 cells, and then scored the topic by calculating the average similarity of those top 50 cells to the centroid by averaging the dot products of the topic probability vectors and the mean topic probability vector. Then, we ranked the topics by this centroid similarity and identified a threshold of 0.2 to separate the candidate cell type topics from the others (Fig. S6).

Next, we assigned cells to clusters defined by these topics. Any cell with greater than 50% probability in one of the cell type topics was automatically assigned to that topic cluster. Some topic clusters had many high probability cells, while some had few. For any clusters with fewer than 150 cells, we attempted to add nearby unassigned cells based on their distance from the cluster’s centroid in a 10-dimensional UMAP space (McInnes et al., 2018). We used an iterative procedure for each small topic cluster as follows: First, we computed the cluster centroid in UMAP space by averaging 200 samples from a Gaussian kernel density estimate of the shape of the cluster. Next, we used a KDTree to identify a set of nearest neighbor cells to the centroid that was 25% larger than the cluster size. We detected whether any of these nearest neighbors were distance outliers (and thus more likely from a different cluster) by ranking the neighbors by distance from the centroid and convoluting the resulting distances with a step function to detect regions where the slope of the ranked distances dramatically increases. We added to the topic cluster the unassigned cell closest to the centroid as long as its value from the convolution was not greater than 1.5 times the interquartile range of all convolution values. Then we iterated this procedure, growing the small topic clusters one unassigned cell at a time. After this procedure, any topic clusters that still had fewer than 50 associated cells were removed from consideration as clusters and their cells were not assigned to any cluster.

To visualize the resulting topic clusters, we used UMAP (McInnes et al., 2018) to reduce the dimensions of the theta matrix to 2 (Fig. 3b). We first row-normalized the theta matrix with the L2-norm and then used the Python implementation of UMAP (`umap-learn`, v. 0.3.8) with default parameters. Finally, we plotted the cells as a scatter plot based on their coordinates in 2D UMAP space, and colored the cells by their topic cluster assignments.

### 4.11 Calling peaks for each topic cluster

The final step in the clustering pipeline is to call peaks for each cluster. For each topic cluster, we pooled the cut sites from the cells in that cluster, and used the pooled data as input to MACS2 (Zhang et al., 2008) (v. 2.2.5) with the same settings as in the alignment pipeline, including providing the bulk ATAC-seq data from naked genomic DNA as an input control. After calling peaks for each cluster, we merged the peak calls from all clusters using `bedtools merge` to make a master list of peak regions, and then used this master peak list to create a new cells-by-peaks matrix. We used our new cells-by-peaks matrix as input to a second round of LDA and cell clustering to refine our clusters and peak calls.

### 4.12 Overlapping peaks with other data sets

Supplementary data for figure 2 from Jänes, et al. 2018 (Jänes et al., 2018) was downloaded from the eLife website (filename: `janes2018_fig2_data1_v2.txt`). This file was parsed using Unix tools and `bedtools` (v. 2.25.0) (Quinlan and Hall, 2010) into three files: a file containing all of the peaks in BED format with overlapping sites merged (i.e. using `bedtools merge` with default parameters), a file containing promoter-annotated peaks (those annotated as “coding_promoter”, “unassigned_promoter”, or “pseudogene_promoter”) with overlapping peaks merged, and a file containing enhancer-associated peaks (those annotated as “putative_enhancer”) with overlapping peaks merged. These files were overlapped with the sci-ATAC-seq peaks, and the sci-ATAC-seq peaks were overlapped with these files, using `bedtools intersect`. The significance of the extent of the overlapping peaks was calculated using the Fisher’s exact test implementation in the `bedtools fisher` command.

Peak loci from the modERN project were downloaded from the EPIC website (http://epic.gs.washington.edu/modERN/) for reference WS245/ce11 using the “Download Aggregated Peaks” and “Download Clustered Peaks” buttons on the “Worm By LifeStage” tab of the user interface. Peaks were parsed into different files based on developmental stage, and any overlapping peak regions were merged in the final files. As above, these files were overlapped with the sci-ATAC-seq peaks, and the sci-ATAC-seq peaks were overlapped with these files using `bedtools intersect`. The significance of the extent of the overlapping peaks was calculated using the Fisher’s exact test implementation in the `bedtools fisher` command.

### 4.13 Cell-by-topic and peak-by-topic heatmaps

To generate the heatmaps as in Fig. 3c and Fig. 7, we selected only the columns corresponding to the topics used for clustering the cells. Next, we row-normalized the counts by dividing each row by its sum. Then we hierarchically clustered the rows and columns in Python (v. 3.6.10) with the `scipy` module (v. 1.4.1). We used the `scipy.spatial.distance.pdist` function with the `cosine` metric to compute the pairwise distance matrix, then `scipy.cluster.hierarchy.linkage` with method `average` to generate clusters, and finally `scipy.cluster.hierarchy.dendrogram` to order the rows and columns after clustering. The clusters were classified by tissue type based on the dominant tissue type for each topic in Fig. 5. The cell/peak coverage information was calculated as the log_2_-transformed sum of peaks found per cell for the cell-by-topic matrix or the log_2_-transformed sum of cells exhibiting a peak for the peak-by-topic matrix.

### 4.14 Identifying tissue-specific topics

We identified topic tissue-specificity in two ways: by overlapping peaks from each topic with peaks from ChIP-seq of cell type-specific factors (Fig. 4), and by assigning peaks to the nearest gene and assessing which tissues those genes tend to be expressed in (Fig. 5).

To compare the overlap of topic-associated sci-ATAC-seq peaks with TF ChIP-seq peaks, we created separate BED files for peaks from ChIP-seq experiments for HLH-1, ELT-1, or ELT-2, removed any peaks that had overlaps in 40 or more other ChIP experiments (i.e. high occupancy target, or HOT, sites), and merged any remaining overlapping peaks with `bedtools merge`. We intersected the peaks called from each topic cluster with the peaks from each transcription factor and recorded the number of overlapping peaks for each topic. In order to understand whether the observed overlaps per topic were surprising, we generated a null distribution by sampling a number of sci-ATAC-seq peaks equal to the number of observed overlaps, with each peak being drawn from a particular topic with a probability based on the total number of peaks called for that topic cluster. We then took the log_2_ ratio of the topic distribution of observed overlaps to the topic distribution of each sample of randomly-drawn peaks. We plot the mean log_2_ ratio, and use the samples to compute a 95% confidence interval around each bar (Fig. 4).

We also used previously published sci-RNA-seq data (Cao et al., 2017) to comprehensively identify cell types for our topic clusters by assessing the tissue expression patterns of genes near our topic-specific peaks. For each topic, we ranked the peaks by their topic-specificity, which we define for a given peak and topic as the fraction of cells with evidence for that peak that are members of the topic’s cluster. We wrote a BED file for each topic cluster that contained the coordinates of either the top 250 peaks by topic-specificity or all peaks with topic-specificity greater than 0.5, whichever was greater. Next, we associated these topic-specific peaks with their nearest downstream gene (within 1200 bp) by using `bedtools closest` with options -D b, -io, and -id). Similar to the cell type-specific TF ChIP-seq analysis above, we then asked whether the expression distribution of these top genes across tissues is enriched on average for particular tissues. Accordingly, we drew 100 samples of 250 genes from the null distribution of all genes with sci-RNA-seq data, and we compared the expression distribution of these gene sets with our sets of topic-specific genes by computing the log_2_ ratio of the mean topic-specific tissue expression distribution to the mean tissue expression distribution of each random sample. We again reported the mean log_2_ ratio and computed the 95% confidence interval around the mean. See Figure S7 for an example illustrating this procedure, and Figure S9 for the tissue expression distributions associated with all 37 topic clusters.

### 4.15 Identifying tissue subtypes using marker genes

In order to identify fine-grained cell types, we conducted a sub-clustering analysis for each of the following major worm tissues that we identified: coelomocyte (Figs. S17, S18), glia (Figs. S19, S20), gonad (Figs. S21, S22), hypodermis (Figs. S23, S24), intestine (Figs. 8, S11), muscle (Figs. 8, S10), neuron (Figs. 9, S12, S13, S14, S15, S16), and pharynx (Figs. S25, S26). There were too few cells identified as sex myoblast to conduct a sub-clustering on that tissue. First, we pooled the data for the cells in the topic clusters corresponding to each tissue, and also merged the peaks called from these topic clusters, to create a data set per tissue (Table 1).

Next, we ran the same iterative LDA-modeling and cell-clustering procedure as detailed above for the whole worm. We identified subclusters by coloring the UMAP scatter plots by the cells with peaks near cell type-specific marker genes (see Figs. 8, 9). We colored a cell for a marker gene if it showed evidence for any peak that either overlapped the gene body or overlapped the region from 1200 bp upstream (5’) of the gene to 100 bp downstream (3’) of the gene. The marker genes we plot were used to identify fine-grained cell types in embryonic and L2 *C. elegans* scRNA-seq data (Packer et al., 2019).

### 4.16 Generating UCSC Genome Browser tracks for the topic clusters

To generate signal files in bigWig format for display in the UCSC Genome Browser, we used the MACS2 output from our peak calling steps. The data from the input control BEDGraph file was subtracted from the data in the treatment BEDGraph using `macs2 bdgcmp -m subtract`, and the resulting BEDGraph file was converted to bigWig format using the `bedGraphToBigWig` utility from UCSC (Kent et al., 2010). These bigWig tracks are displayed along with our peak calls in two track hubs on the UCSC Genome Browser:

sci-ATAC-seq Main Track Hub URL: http://genome.ucsc.edu/cgi-bin/hgTracks?db=ce11&hubClear=http://waterston.gs.washington.edu/atacTissue/Durham_hub.txt sci-ATAC-seq Subclustering Track Hub URL: http://genome.ucsc.edu/cgi-bin/hgTracks?db=ce11_hubClear=http://waterston.gs.washington.edu/atacCellType/Durham_hub.txt

## 5 Data and code availability

All raw and processed sequencing data generated in this study have been submitted to the NCBI Gene Expression Omnibus (GEO; https://www.ncbi.nlm.nih.gov/geo/). The parallel LDA code is available through GitHub at https://github.com/gevirl/LDA and is released under the GNU GPLv3 license. The data pipeline, scripts for performing the LDA clustering analysis, and Jupyter notebooks for generating figures are available at the following GitHub repository: https://github.com/tdurham86/L2_sci-ATAC-seq and is released under the MIT license.

## 6 Acknowledgements

We want to thank Anh Leith for sharing her expertise in FACS and helping us to collect samples and run the sci-ATAC protocol. Chau Huynh for guidance when developing the worm nuclei isolation protocol and when optimizing our early experiments on bulk ATAC-seq. Olubusayo Bolonduro, as a summer intern in the Waterston Lab, also helped optimize conditions for ATAC-seq in worm and performed the input control experiment to collect ATAC-seq data from naked *C. elegans* genomic DNA. We would like to thank Illumina for providing the indexed Tn5, and also PCR reagents for the preparation of some of our sci-ATAC-seq libraries. This work was funded by NIH grants U41HG007355, R01GM072675, and R01HG010478 to RHW. JS is a Howard Hughes Medical Institute investigator.

## 7 Author contributions

TJD and RHW conceived of the experiment. TJD grew the worms and isolated the nuclei. TJD and RMD performed the sci-ATAC-seq experiments. DAC helped perform the pilot sci-ATAC-seq experiment, and shared his code for the initial processing and QC of the sequencing data. JS provided important reagents and lab equipment for the sci-ATAC-seq protocol. TJD analyzed the data and wrote the paper with input from RHW and WSN. LG wrote the parallelized LDA Java program. RHW, LG, and WSN contributed to discussions of the data analysis.

## 9 Supplementary Figures

**Supplementary Figure S1:**
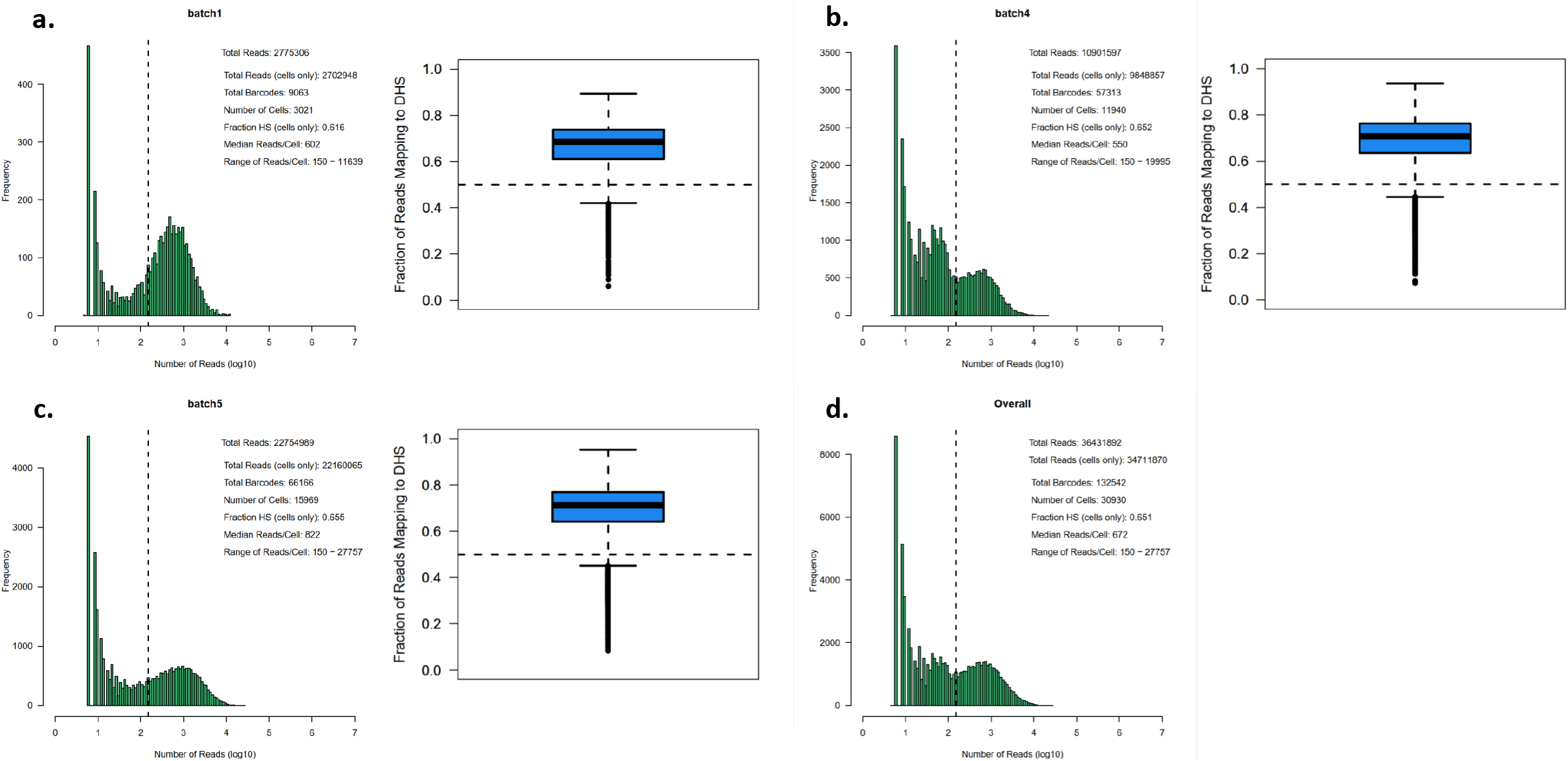
After thresholding cell coverage distribution, we recover a total of 30,930 cells from three sequencing batches. We show the histograms of unique reads per cell barcode and the distribution of the fraction of reads mapping in a peak region for each sequencing batch, including **a.** the initial pilot batch (MiSeq sequencing), **b.** large-scale batch number 1 (NextSeq), and **c.** large-scale batch number 2 (NextSeq). The final panel **d.** shows the aggregated read coverage statistics for all three batches.

**Supplementary Figure S2:**
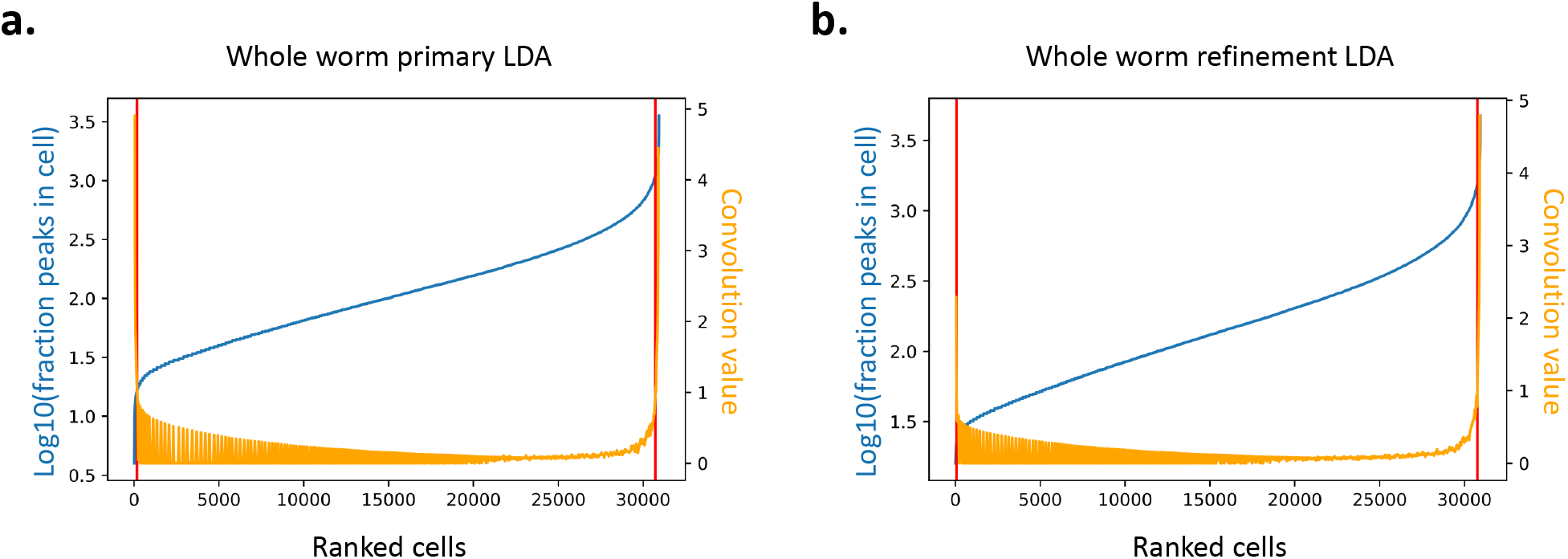
Filtering cells with too few peaks. Cells were ranked by the number of peaks detected (blue line), and cells with too few peaks were filtered out. The threshold (lefthand red vertical line) was determined by automatically finding the inflection point in the ranking curve (orange line). **a.** Filtering cells before the whole-worm primary LDA iteration. **b.** Filtering cells before the whole-worm refinement LDA iteration.

**Supplementary Figure S3:**
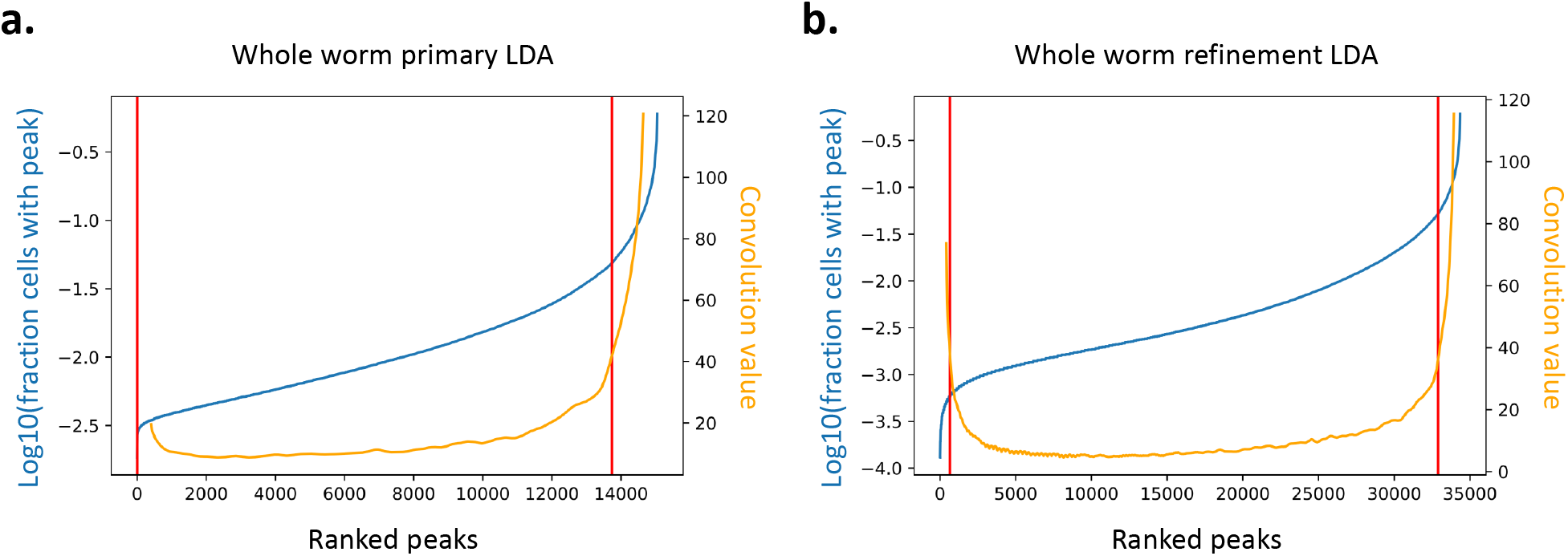
Filtering peaks found in too many or too few cells. Peaks were ranked by the fraction of cells in which they were detected (blue line), and outlier peaks were filtered out. The thresholds (red vertical lines) were determined by automatically finding the inflection points in the ranking curve (orange line). **a.** Filtering peaks before the whole-worm primary LDA iteration. **b.** Filtering peaks before the whole-worm refinement LDA iteration.

**Supplementary Figure S4:**
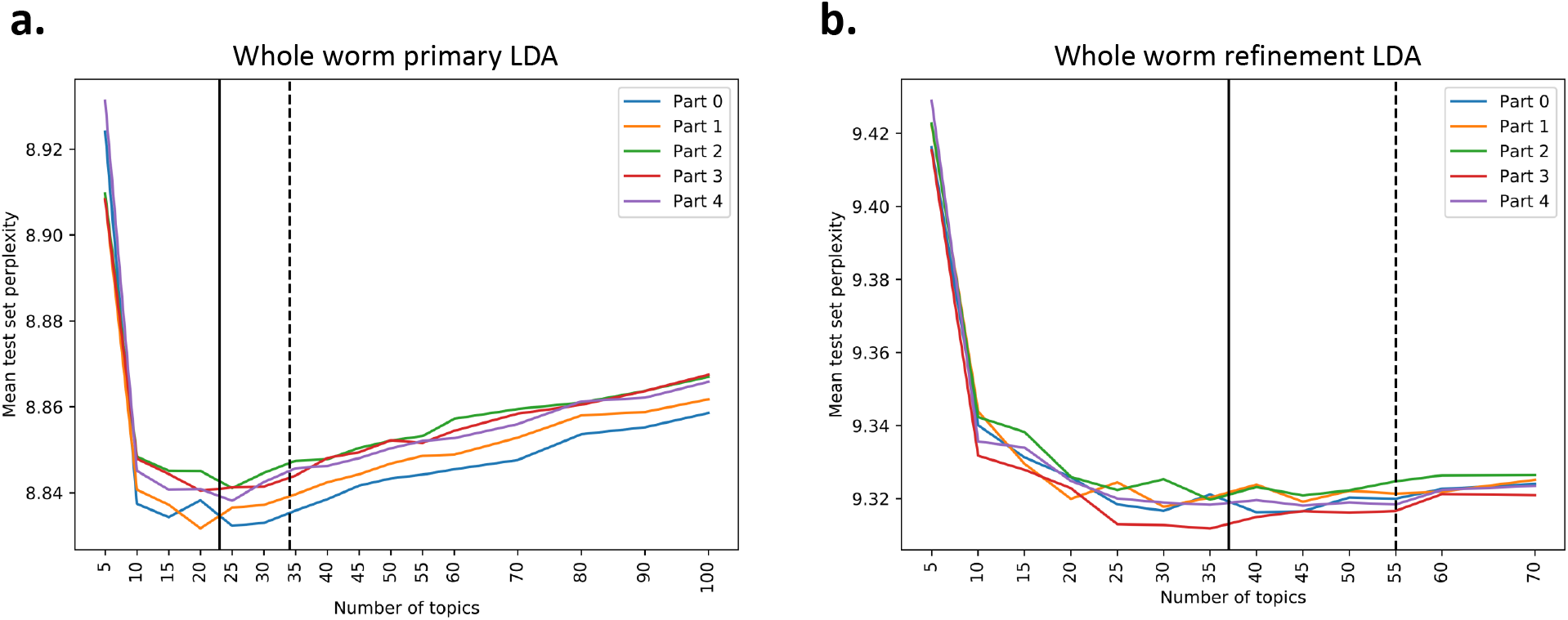
Tuning the number of topics using 5-fold cross validation. Models were trained on 4 folds and tested on a held-out fold for varying numbers of topics. The average minimum topic number (solid line) was calculated, and used as the basis to pick a number of topics 1.5 times greater for use in training the full LDA model (dotted line). **a.** Topic number search for the whole-worm primary LDA. **b.** Topic number search for the whole-worm refinement LDA.

**Supplementary Figure S5:**
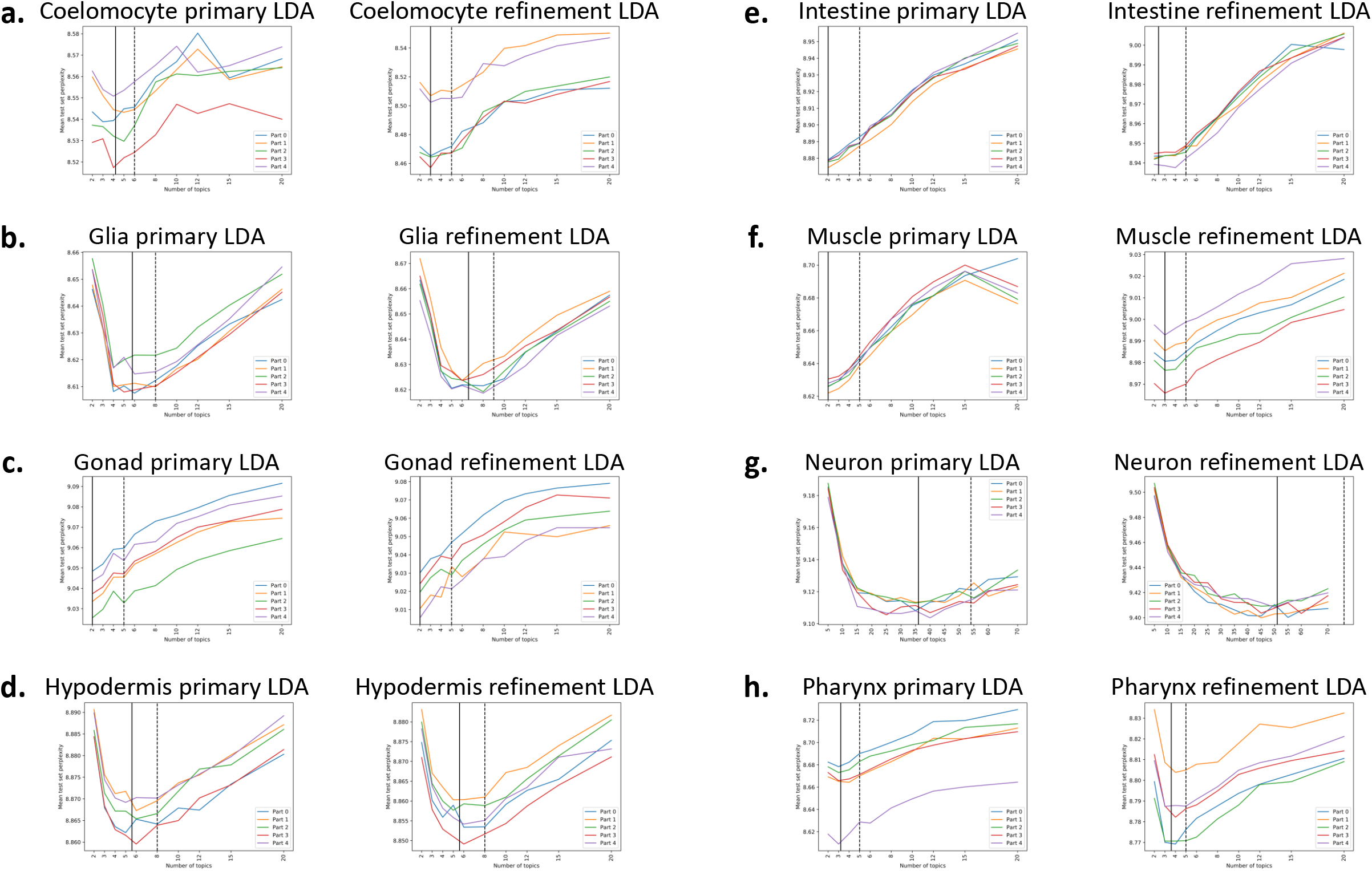
Tuning the number of topics using 5-fold cross validation. Models were trained on 4 folds and tested on a held-out fold for varying numbers of topics. The average minimum topic number (solid line) was calculated, and used as the basis to pick a number of topics 1.5 times greater for use in training the full LDA model (dotted line). Pairs of plots show the topic number search for the tissue-specific primary LDA (left plot) and tissue-specific refinement LDA (right plot) for **a.** coelomocyte, **b.** glia, **c.** gonad, **d.** hypodermis, **e.** intestine, **f.** muscle, **g.** neuron, **h.** pharynx.

**Supplementary Figure S6:**
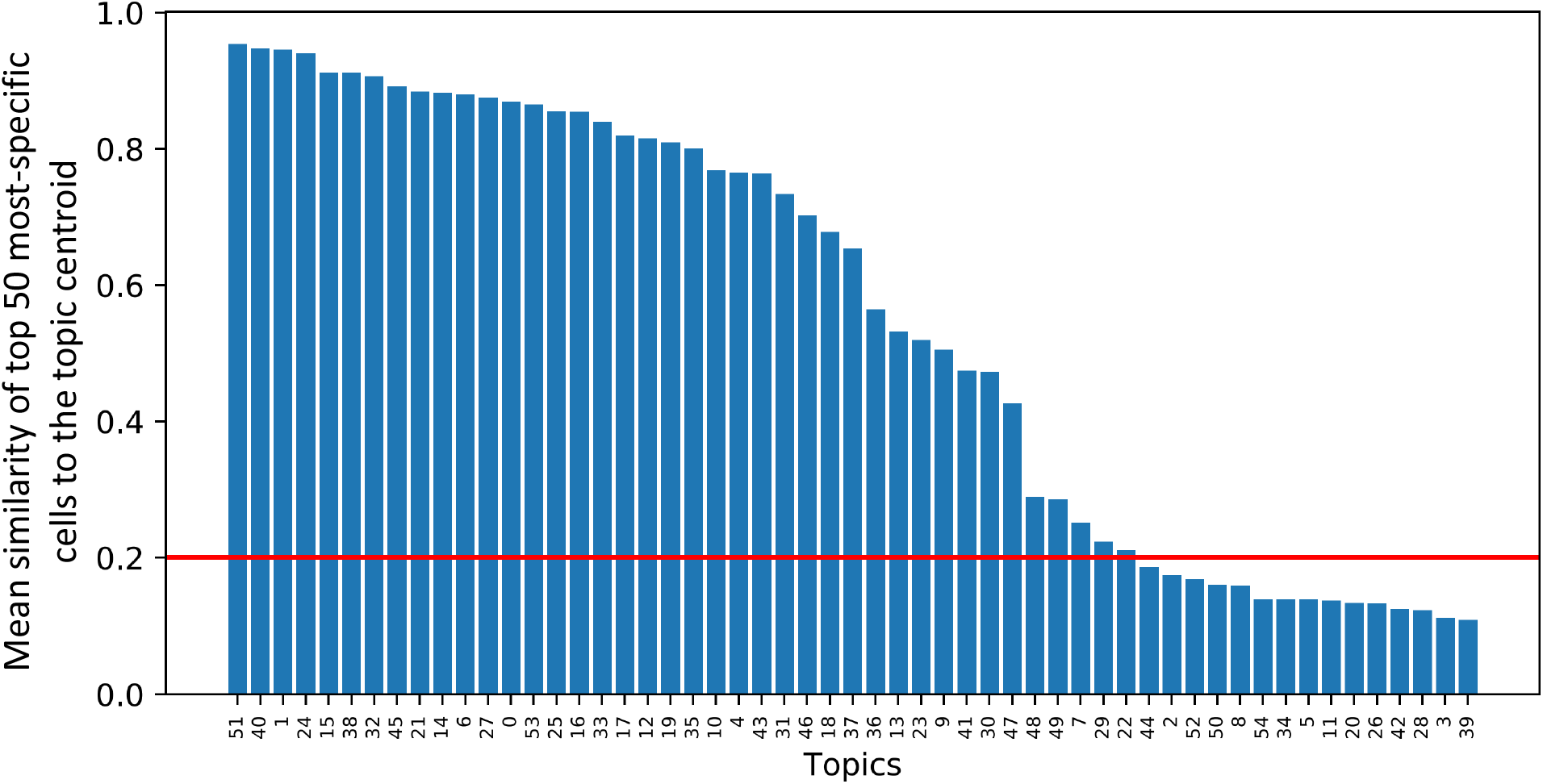
Identifying topics that capture clusters of cells. Topics were ranked by the mean similarity of their top 50 most-specific cells to the average topic distribution of those same 50 cells. Topics were considered for further analysis if their mean similarity exceeded the 0.2 threshold (red line).

**Supplementary Figure S7:**
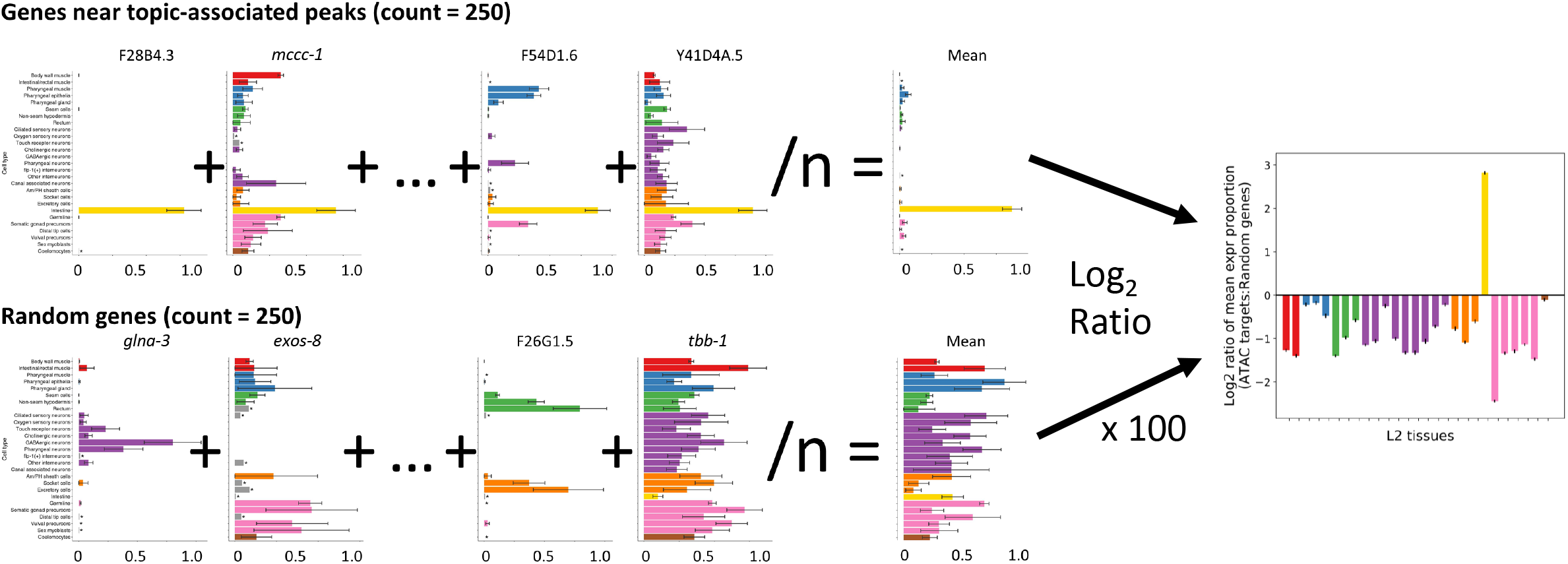
Schematic describing how to compute tissue enrichment. For Fig. 5, we computed the tissue enrichment values by normalizing the tissue expression values for each gene to sum to one, then calculating the log_2_ ratio of the mean expression distribution for the top 250 genes by peak topic-specificity to the mean tissue expression distribution of 250 randomly-selected genes. This was repeated for 100 random samples of 250 genes, and the mean log_2_ ratio was plotted with error bars indicating the 95% confidence interval for the enrichment of each tissue.

**Supplementary Figure S8:**
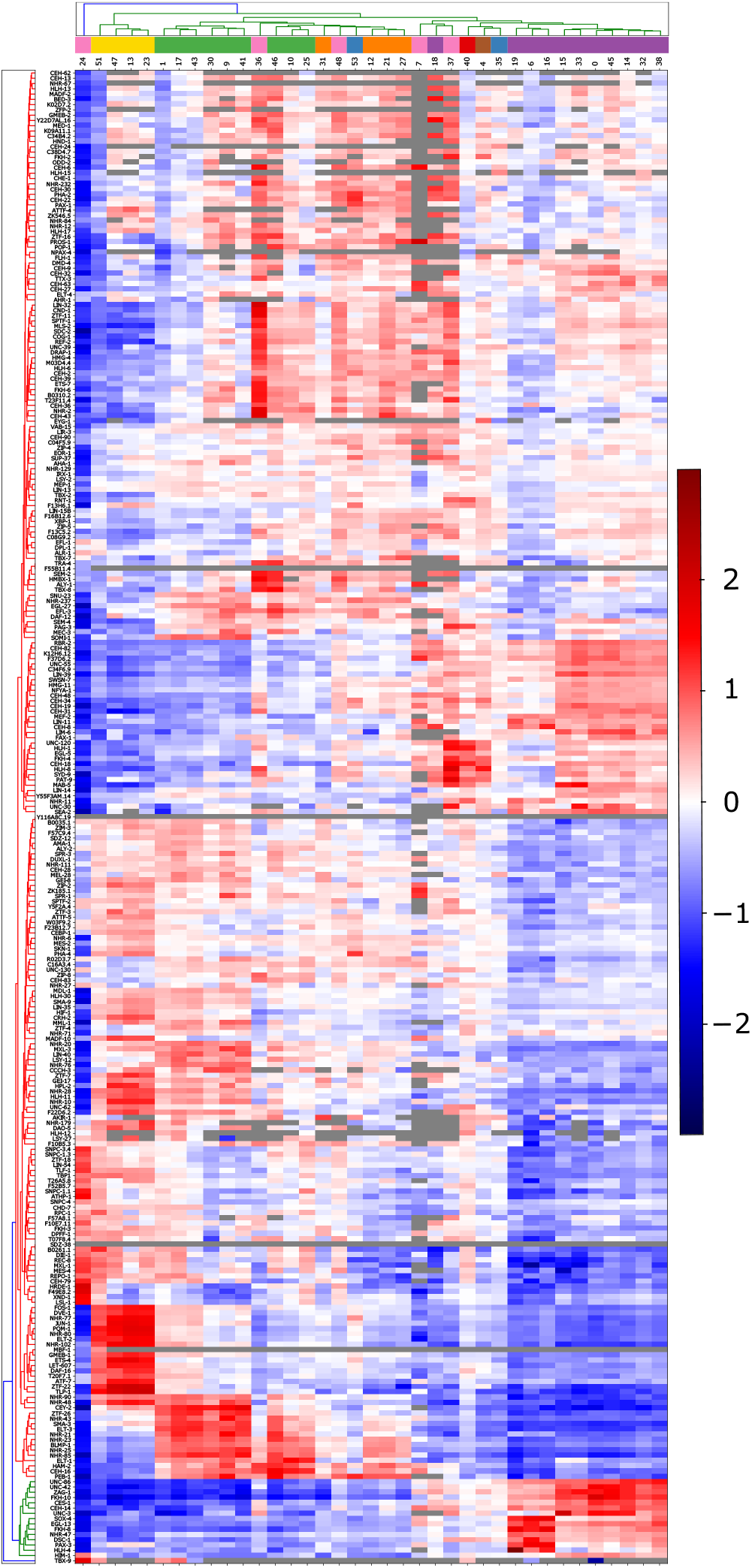
Sci-ATAC-seq peaks called from the whole-worm refinement LDA analysis show enrichment and depletion for overlaps with TF ChIP-seq peaks that is consistent with the TF tissue-specificity. We extended the analysis shown in Fig. 7 to all TFs with ChIP-seq data. The heatmap shows the mean *log_2_* ratio of the overlap counts for the peaks called based on each LDA topic to 500 randomly sampled sets of matched size from each topic. Each row displays the results for a single TF across all 37 topics used for clustering in the refinement LDA analysis. The rows and columns are hierarchically clustered based on Euclidean distance, using `sklearn.metrics.pairwise.nan_euclidean_distances` to ignore any TF/topic pairs with no overlaps (grey in the heatmap). TFs with similar tissue-specific expression patterns tend to show similar enrichment and depletion patterns for overlaps with peaks from different topics, and these patterns are consistent with our inferred tissue identities for the topics (Figs. 5, S9).

**Supplementary Figure S9:**
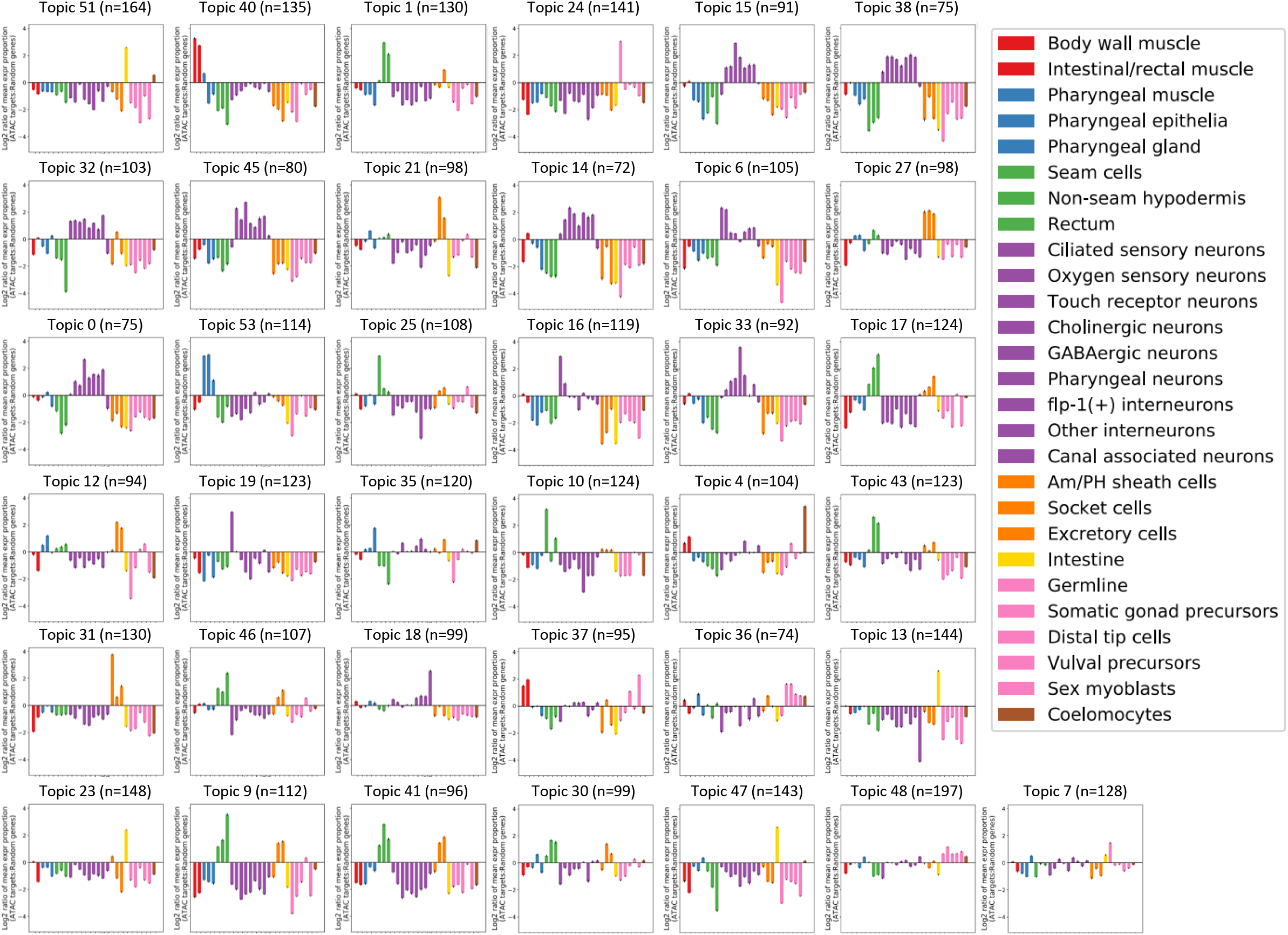
RNA-seq tissue expression enrichment for all 37 topic clusters.

**Supplementary Figure S10:**
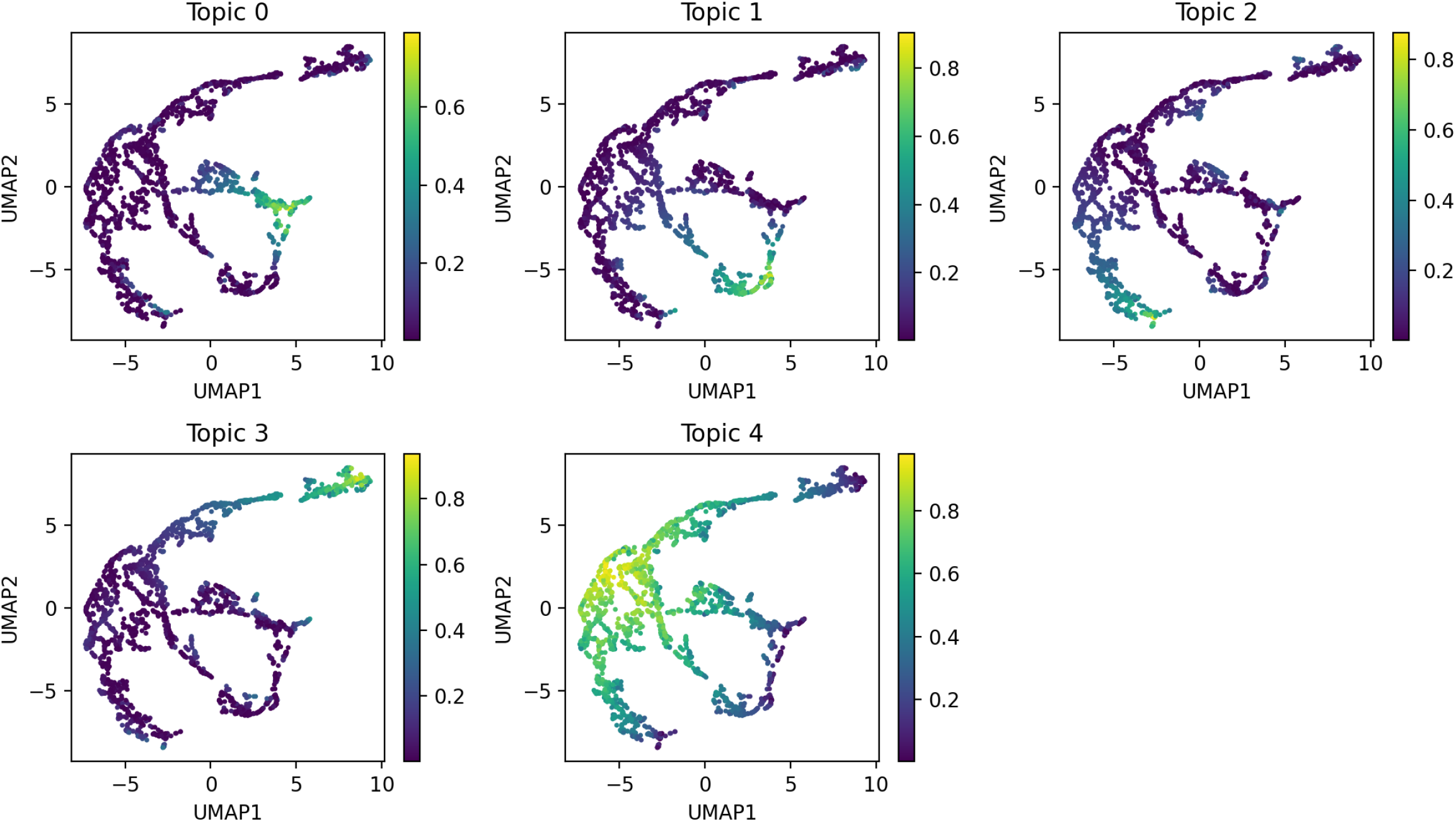
Topic probabilities for muscle subclustering LDA analysis. UMAP plots displaying the results of performing our iterative LDA procedure on only muscle cells (topic 40 in the whole worm refinement LDA, see Fig. S9). Each dot in the scatterplot represents one cell, and in each plot the cells are colored by their probability for each LDA topic.

**Supplementary Figure S11:**
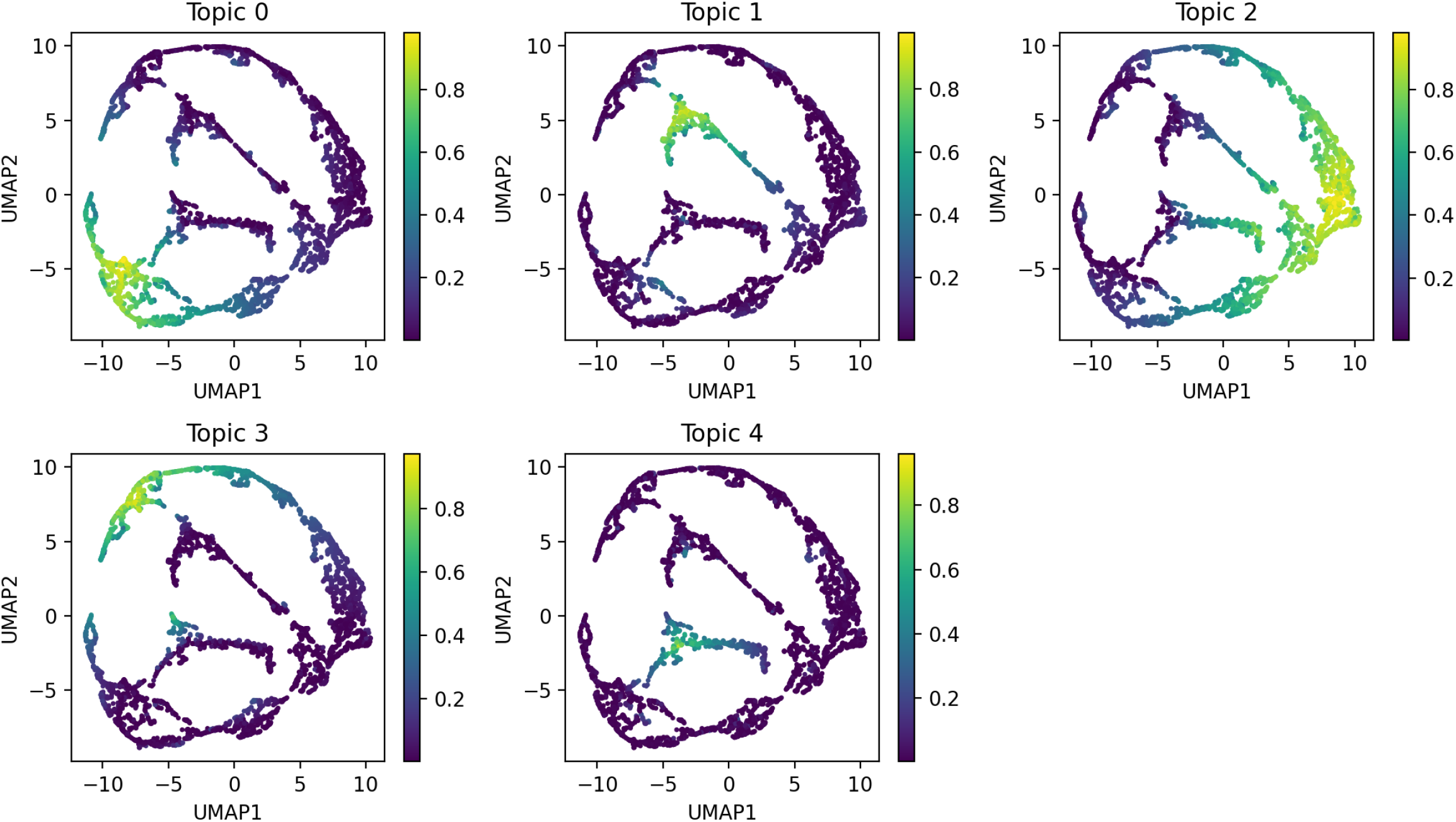
Topic probabilities for intestine subclustering LDA analysis. UMAP plots displaying the results of performing our iterative LDA procedure on only intestine cells (topics 13, 23, 47, and 51 in the whole worm refinement LDA, see Fig. S9). Each dot in the scatterplot represents one cell, and in each plot the cells are colored by their probability for each LDA topic.

**Supplementary Figure S12:**
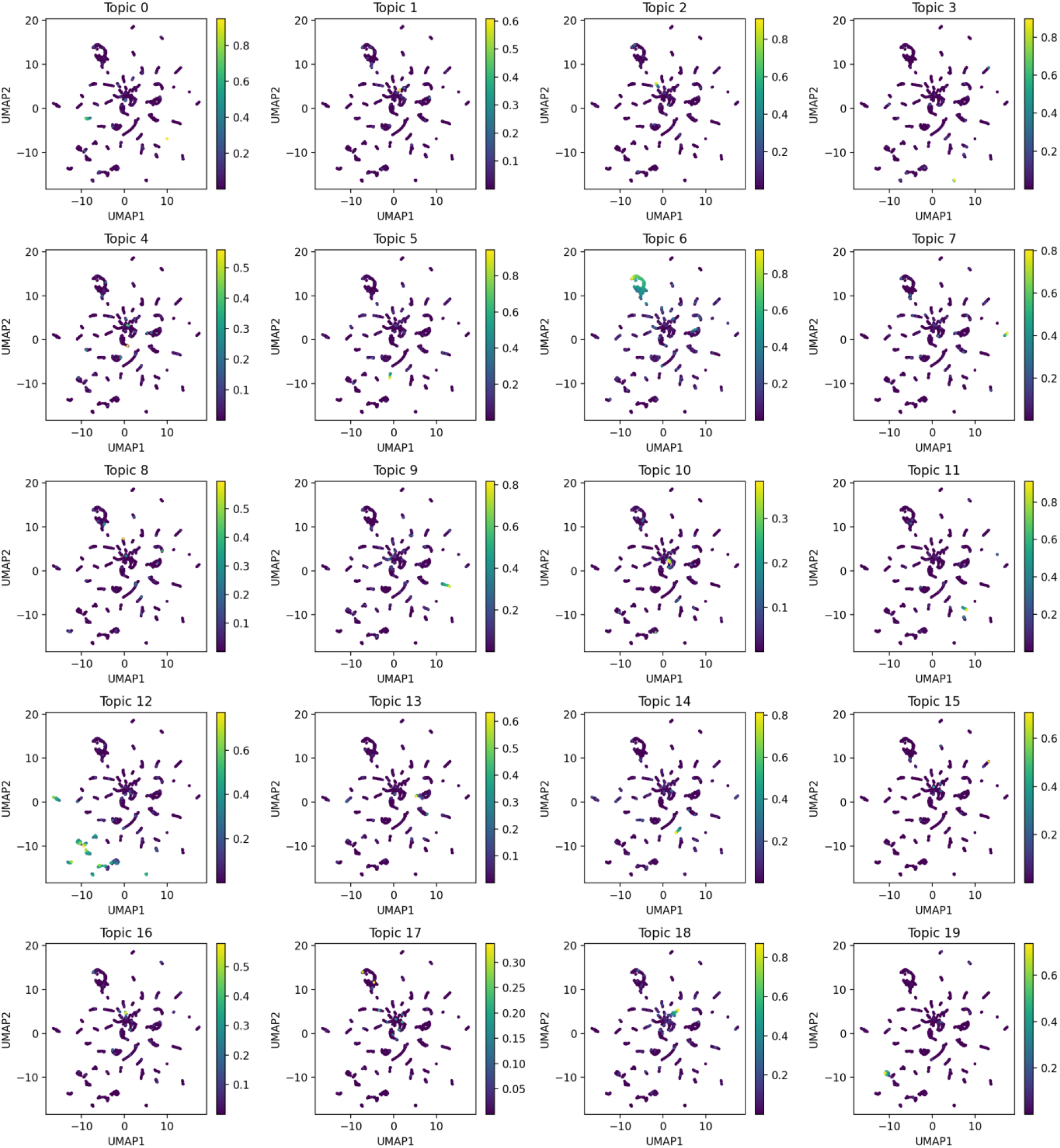
Topic probabilities for neuron subclustering LDA analysis. Part 1. UMAP plots displaying the results of performing our iterative LDA procedure on only neuron cells (topics 0, 6, 14, 15, 16, 18, 19, 32, 33, 38, and 45 in the whole worm refinement LDA, see Fig. S9). Each dot in the scatterplot represents one cell, and in each plot the cells are colored by their probability for each LDA topic.

**Supplementary Figure S13:**
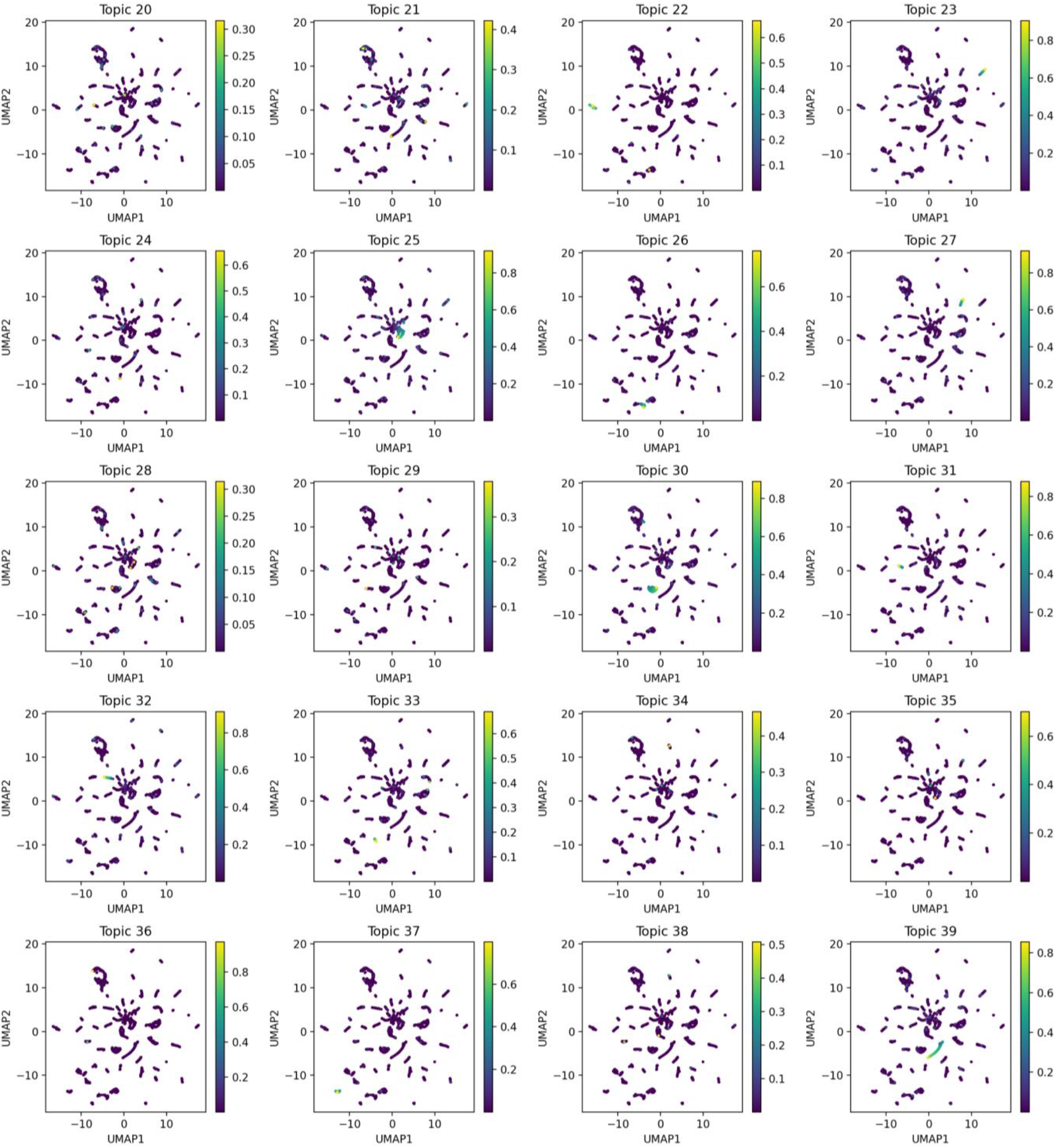
Topic probabilities for neuron subclustering LDA analysis. Part 2. UMAP plots displaying the results of performing our iterative LDA procedure on only neuron cells (topics 0, 6, 14, 15, 16, 18, 19, 32, 33, 38, and 45 in the whole worm refinement LDA, see Fig. S9). Each dot in the scatterplot represents one cell, and in each plot the cells are colored by their probability for each LDA topic.

**Supplementary Figure S14:**
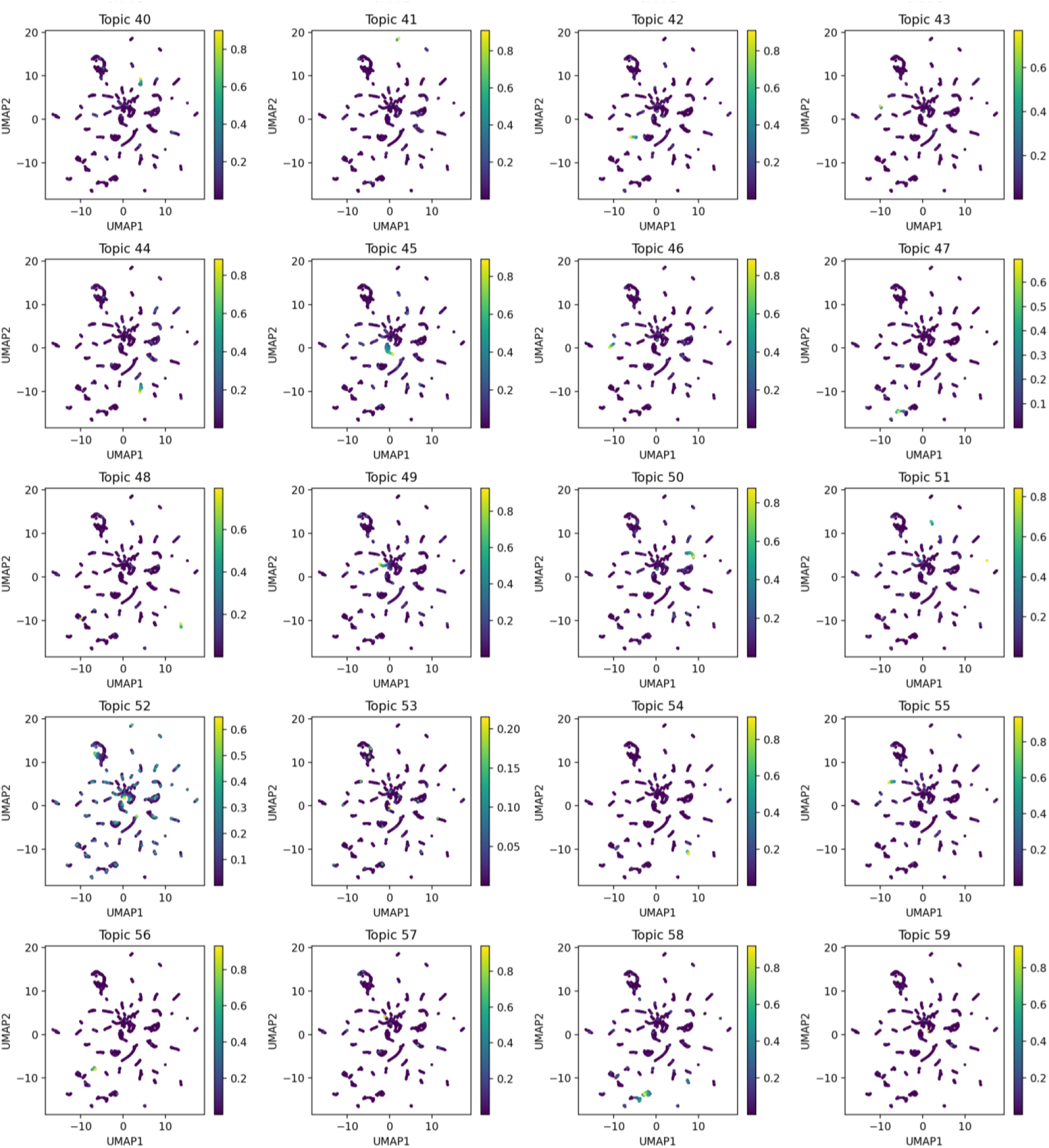
Topic probabilities for neuron subclustering LDA analysis. Part 3. UMAP plots displaying the results of performing our iterative LDA procedure on only neuron cells (topics 0, 6, 14, 15, 16, 18, 19, 32, 33, 38, and 45 in the whole worm refinement LDA, see Fig. S9). Each dot in the scatterplot represents one cell, and in each plot the cells are colored by their probability for each LDA topic.

**Supplementary Figure S15:**
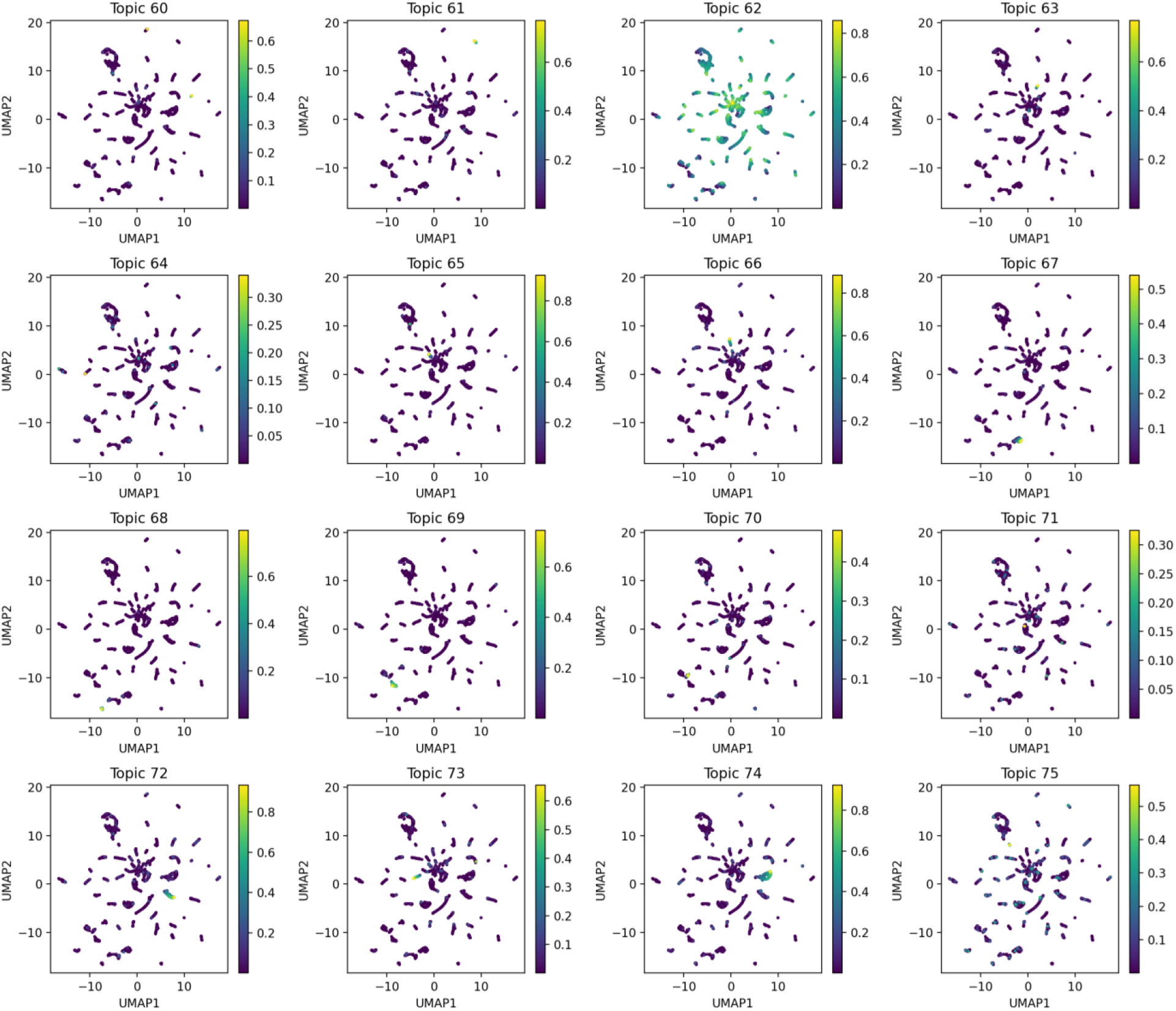
Topic probabilities for neuron subclustering LDA analysis. Part 4. UMAP plots displaying the results of performing our iterative LDA procedure on only neuron cells (topics 0, 6, 14, 15, 16, 18, 19, 32, 33, 38, and 45 in the whole worm refinement LDA, see Fig. S9). Each dot in the scatterplot represents one cell, and in each plot the cells are colored by their probability for each LDA topic.

**Supplementary Figure S16:**
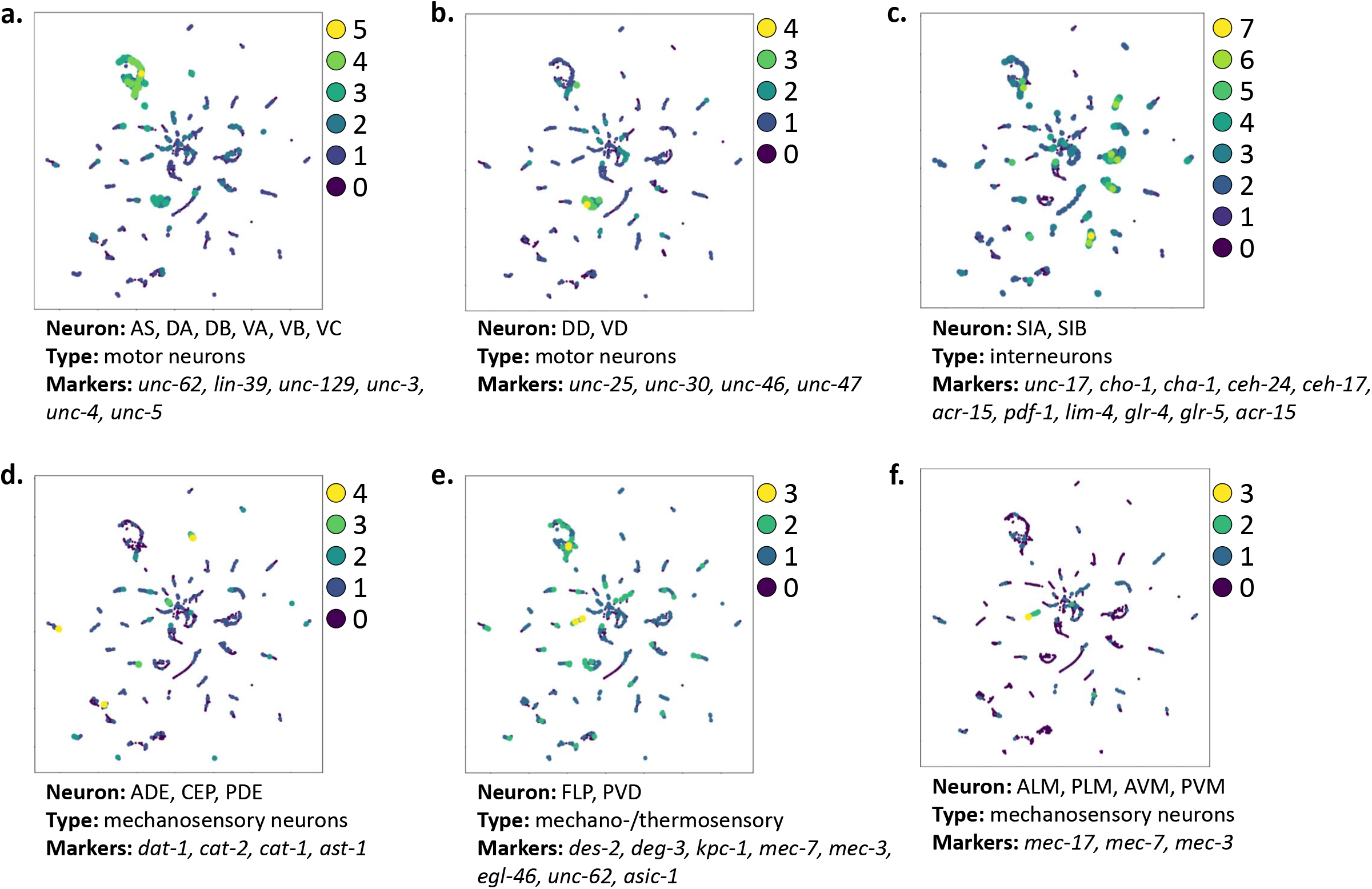
Marker genes identify neuronal types at high resolution. We identified additional neurons in the UMAP plot by plotting the distribution of cells showing peaks of accessibility within 1200 bp upstream or 100 bp downstream of sets of marker genes from Packer, et al. 2019 (Packer et al., 2019). Each scatter plot dot represents a cell, and the number of genes with nearby accessibility in a given cell is shown by the color and size of its dot on the scatter plot, with cells showing accessibility near more marker genes having dots that are larger and more yellow. Information below each plot details the names of the neurons being highlighted, the type of neuron, and the marker genes used to generate the plot.

**Supplementary Figure S17:**
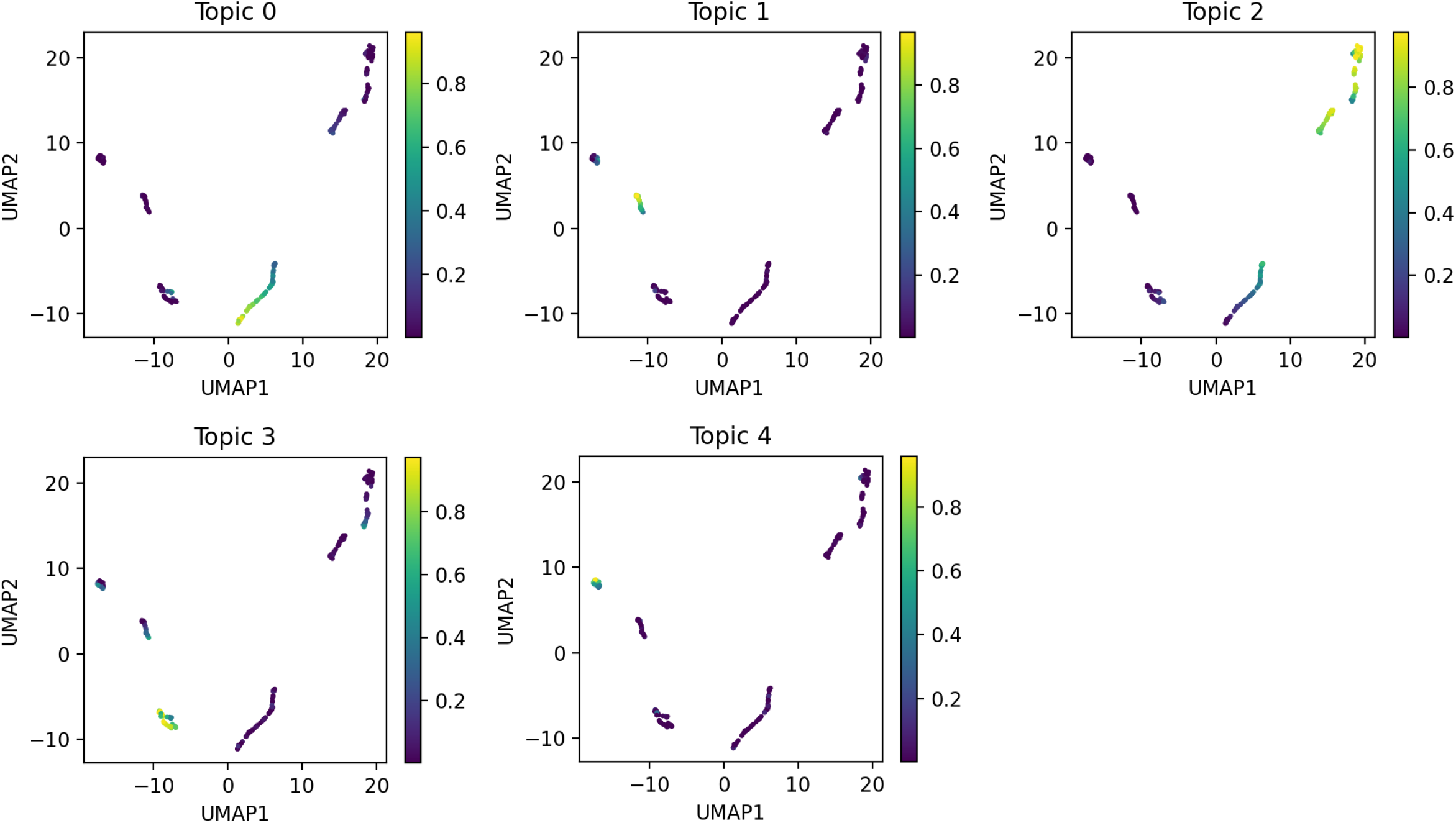
Topic probabilities for coelomocyte subclustering LDA analysis. UMAP plots displaying the results of performing our iterative LDA procedure on only coelomocyte cells (topic 4 in the whole worm refinement LDA, see Fig. S9). Each dot in the scatterplot represents one cell, and in each plot the cells are colored by their probability for each LDA topic.

**Supplementary Figure S18:**
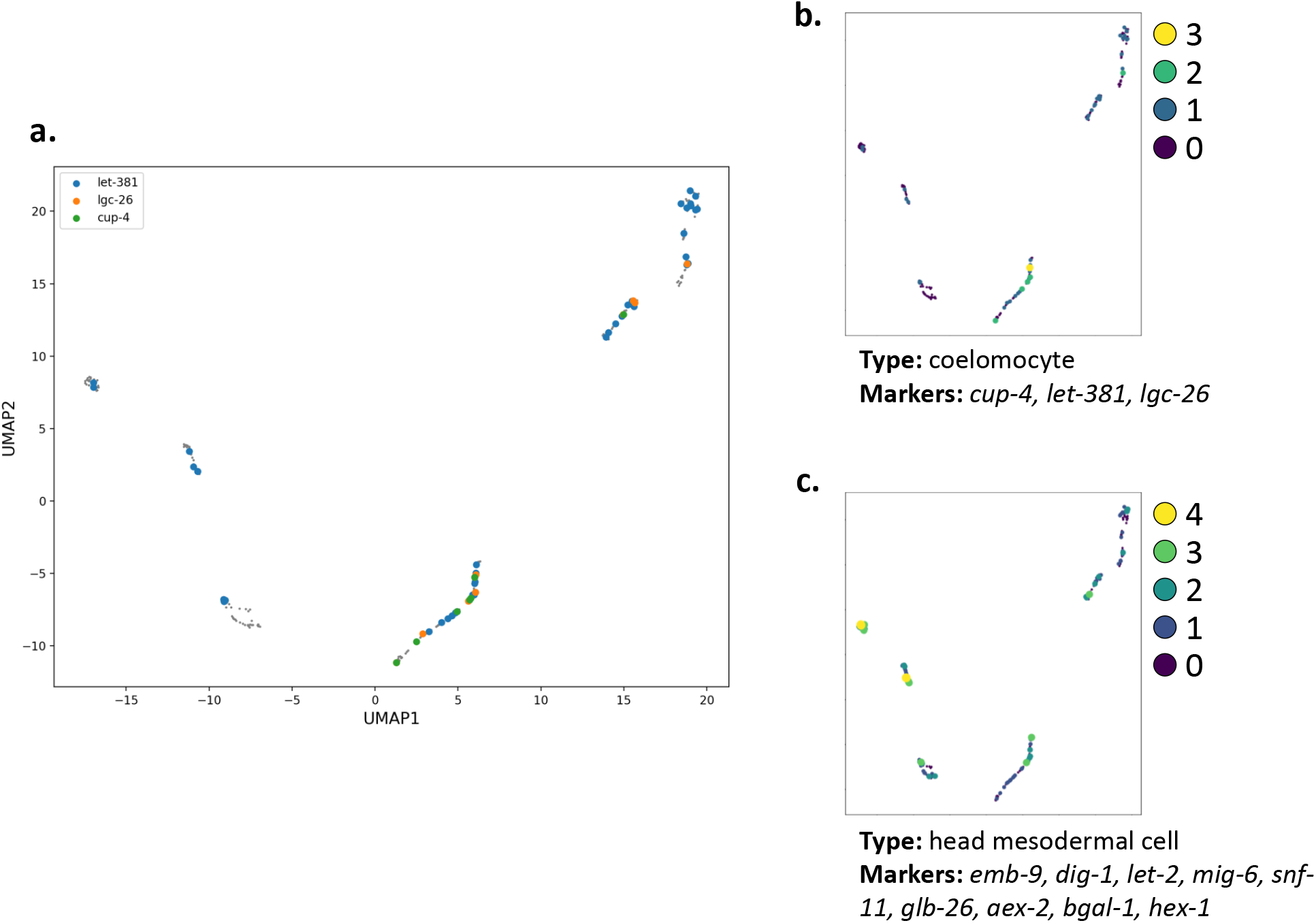
Marker genes identify coelomocyte subclusters. We identified coelomocyte subclusters in the UMAP plot by plotting the distribution of cells showing peaks of accessibility within 1200 bp upstream or 100 bp downstream of sets of marker genes from Packer, et al. 2019 (Packer et al., 2019). **a.** Scatter plot of the UMAP embedding with the cells colored by which of three coelomocyte marker genes show nearby accessibility. **b.** The same coelomocyte marker genes are plotted, but in this case each cell is colored based on how many of the marker genes show nearby accessibility in each cell. **c.** Plotting the number of head mesodermal cell marker genes with nearby accessibility identifies the clusters enriched for topics 1 and 4 (Fig. S17) as candidate head mesodermal cells.

**Supplementary Figure S19:**
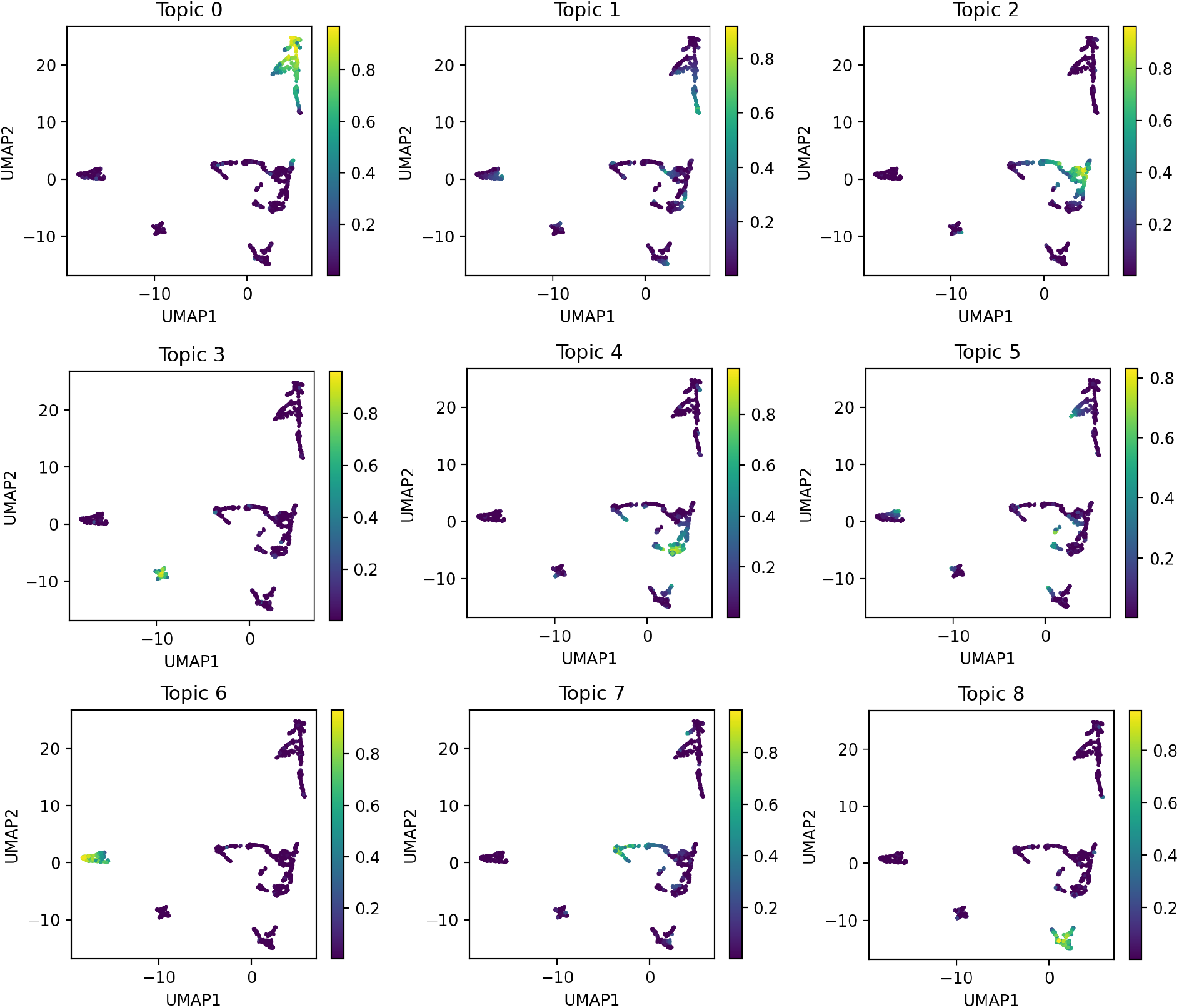
Topic probabilities for glia subclustering LDA analysis. UMAP plots displaying the results of performing our iterative LDA procedure on only glial cells (topics 12, 21, 27, and 31 in the whole worm refinement LDA, see Fig. S9). Each dot in the scatterplot represents one cell, and in each plot the cells are colored by their probability for each LDA topic.

**Supplementary Figure S20:**
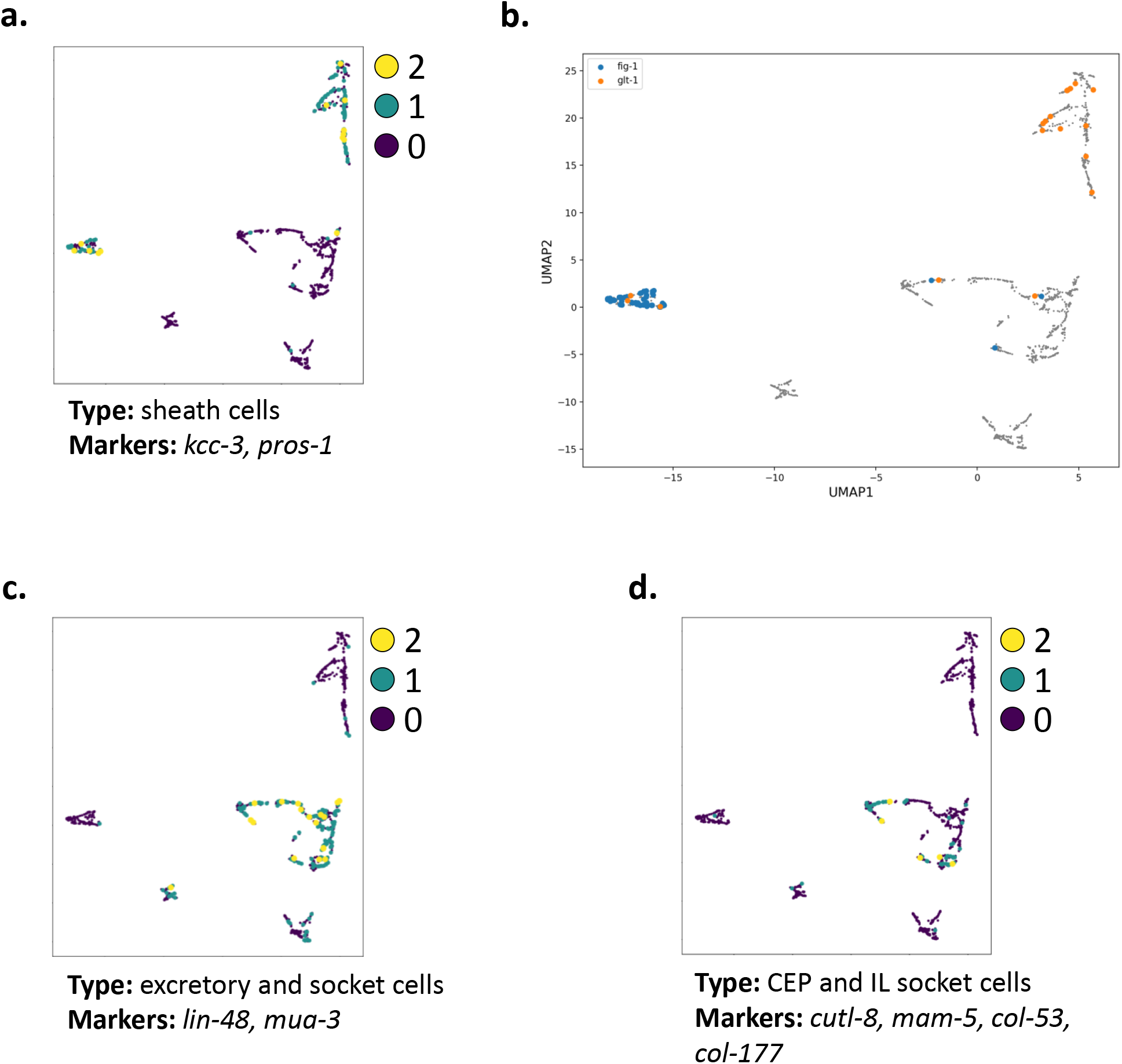
Marker genes identify glia subclusters. We identified glia subclusters by plotting the distribution of cells showing accessibility for sets of marker genes from Packer, et al. 2019 (Packer et al., 2019). **a.** Sheath cells are characterized by expression of marker genes *kcc-3* and *pros-1*. Cells that show accessibility near these genes are predominantly those with high probabilities for topics 0 and 6 (Fig. S19). **b.** The sheath cells can be further subdivided based on the expression of *fig-1*, which marks amphid and phasmid sheath cells, and *glt-1*, which marks cephalic sheath cells. The cells with high probability for topic 0 (Fig. S19) have low accessibility near *fig-1*, but do show accessibility near *glt-1*, identifying them as candidate cephalic sheath cells, while cells with high probability in topic 6 (Fig. S19) show the reverse and are candidate amphid and phasmid sheath cells. **c.** Similarly, coloring the cells by accessibility nearby *lin-48* and *mua-3* show the other cells in the plot are candidate excretory and socket cells, while accessibility near marker genes *cutl-8, mam-5, col-53*, and *col-177* suggest that the cells with high topic 4 and topic 7 probability (Fig. S19) are candidate cephalic and inner labial socket cells (**d.**).

**Supplementary Figure S21:**
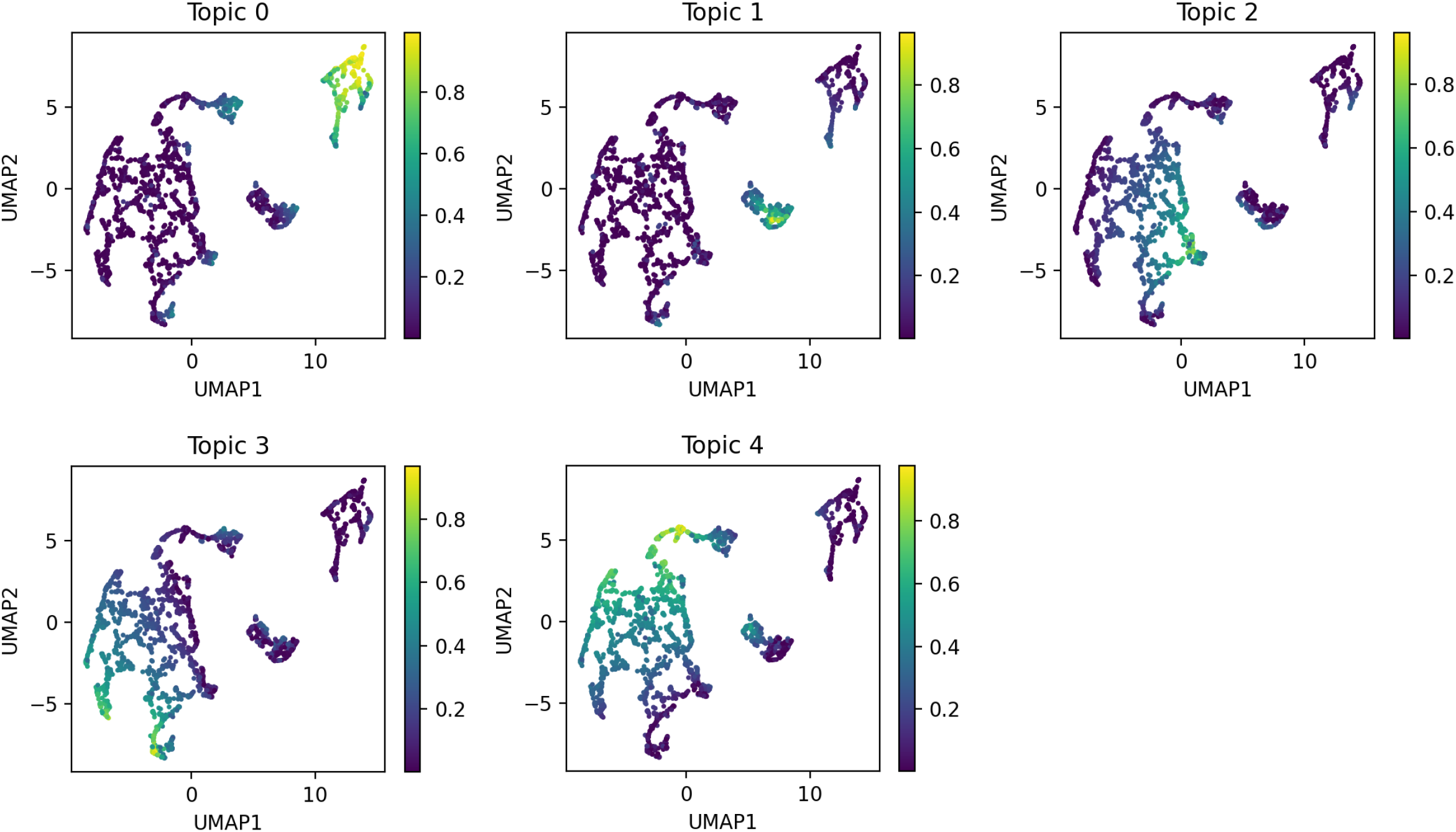
Topic probabilities for gonad subclustering LDA analysis. UMAP plots displaying the results of performing our iterative LDA procedure on only gonad cells (topics 7, 24, 36, and 48 in the whole worm refinement LDA, see Fig. S9). Each dot in the scatterplot represents one cell, and in each plot the cells are colored by their probability for each LDA topic.

**Supplementary Figure S22:**
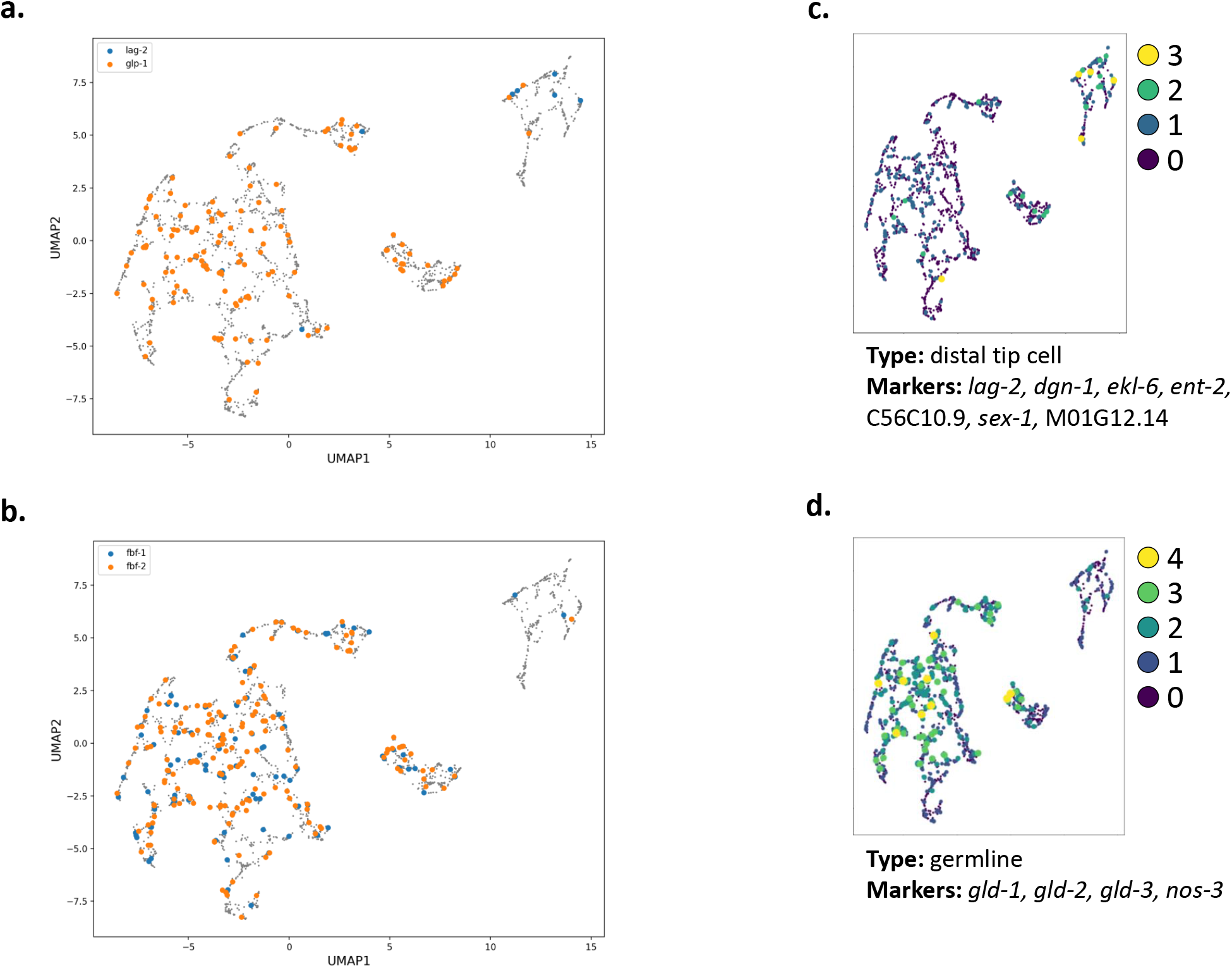
Marker genes identify gonad subclusters. We identified gonad subclusters by plotting the distribution of cells showing accessibility for sets of marker genes from Packer, et al. 2019 (Packer et al., 2019) and Wormbook (Kimble, 2005). The gonad forms with a stem cell niche maintained by the distal tip cells that maintain stemness in the germline by Notch signaling. The distal tip cells produce the Notch ligand LAG-2, while the mitotic germline cells express the receptor, GLP-1. Here, the gonad LDA analysis largely separates the cells with accessibility near these two genes **a.**, suggesting that the cells with high topic 0 probability (Fig. S21) are candidate distal tip cells, while most of the others are candidate germline cells. This observation is also supported by looking for accessibility near the *fbf-1* and *fbf-2* genes **b.**, which encode RNA binding proteins that function downstream of GLP-1 to maintain germ cells in the mitotic state. The candidate distal tip cells also show coaccessibility near other distal tip cell marker genes identified from the single cell RNA-seq data **c.**, and similarly, additional germline marker genes show nearby coaccessibility in the same cells that have accessible sites near the *fbf* genes.

**Supplementary Figure S23:**
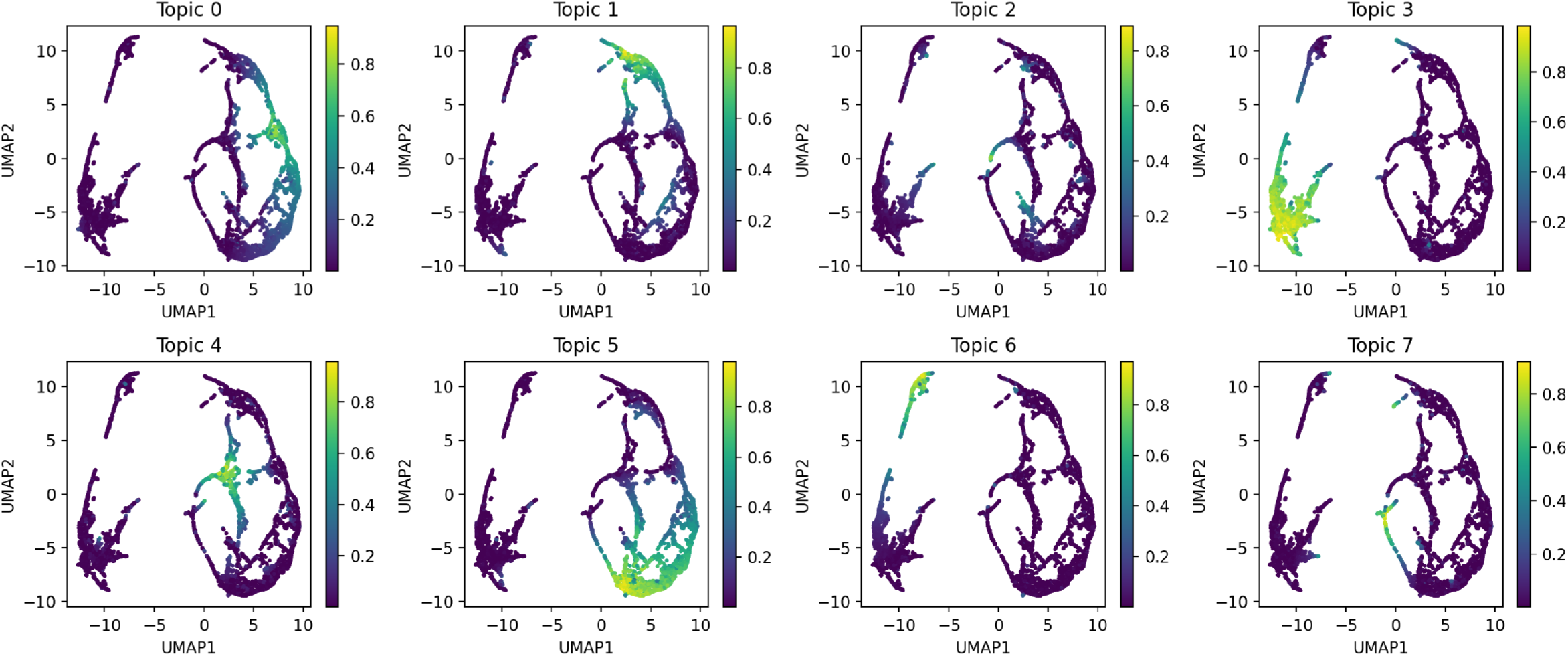
Topic probabilities for hypodermis subclustering LDA analysis. UMAP plots displaying the results of performing our iterative LDA procedure on only hypodermal cells (topics 1, 9, 10, 17, 25, 30, 41, 43, and 46 in the whole worm refinement LDA, see Fig. S9). Each dot in the scatterplot represents one cell, and in each plot the cells are colored by their probability for each LDA topic.

**Supplementary Figure S24:**
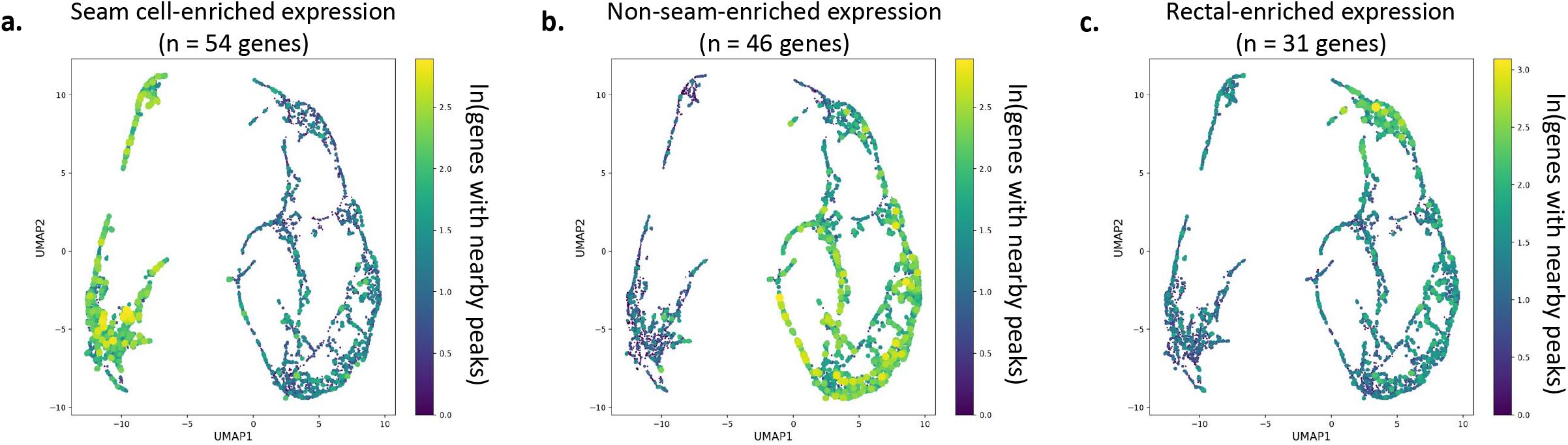
Marker genes identify hypodermis subclusters. To identify hypodermis subclusters, we assessed co-accessibility of sites near genes enriched in expression for the hypodermal tissues identified in Cao et al. 2017 (Cao et al., 2017). The genes that we selected have greater than five-fold enrichment in the specified hypodermis tissue compared to all other tissues, as reported by the GExplore website (http://genome.sfu.ca/gexplore/gexplore_search_tissues.html). **a.** The cells with high probability in topics 3 and 6 (Fig. S23) show high co-accessibility of regions near genes with enriched expression in seam cells. **b.** Genes with enriched expression in non-seam hypodermis have nearby co-accessible sites in cells with high probability in topics 0, 4, 5, and 7 (Fig. S23). **c.** Last, cells with high probability for topic 1 (Fig. S23) tend to have co-accessible sites near genes with enriched expression in rectum.

**Supplementary Figure S25:**
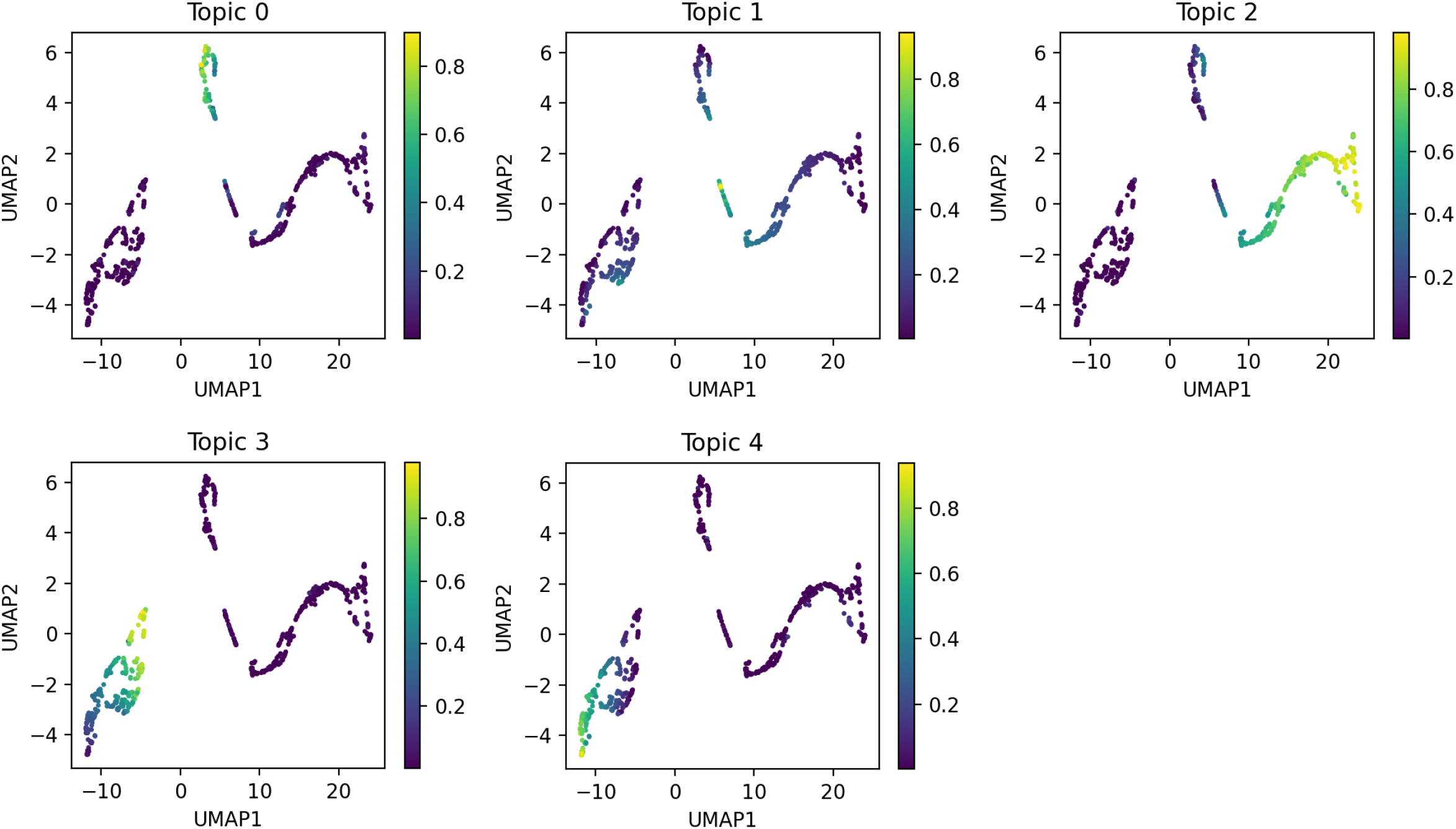
Topic probabilities for pharynx subclustering LDA analysis. UMAP plots displaying the results of performing our iterative LDA procedure on only pharyngeal cells (topics 35 and 53 in the whole worm refinement LDA, see Fig. S9). Each dot in the scatterplot represents one cell, and in each plot the cells are colored by their probability for each LDA topic.

**Supplementary Figure S26:**
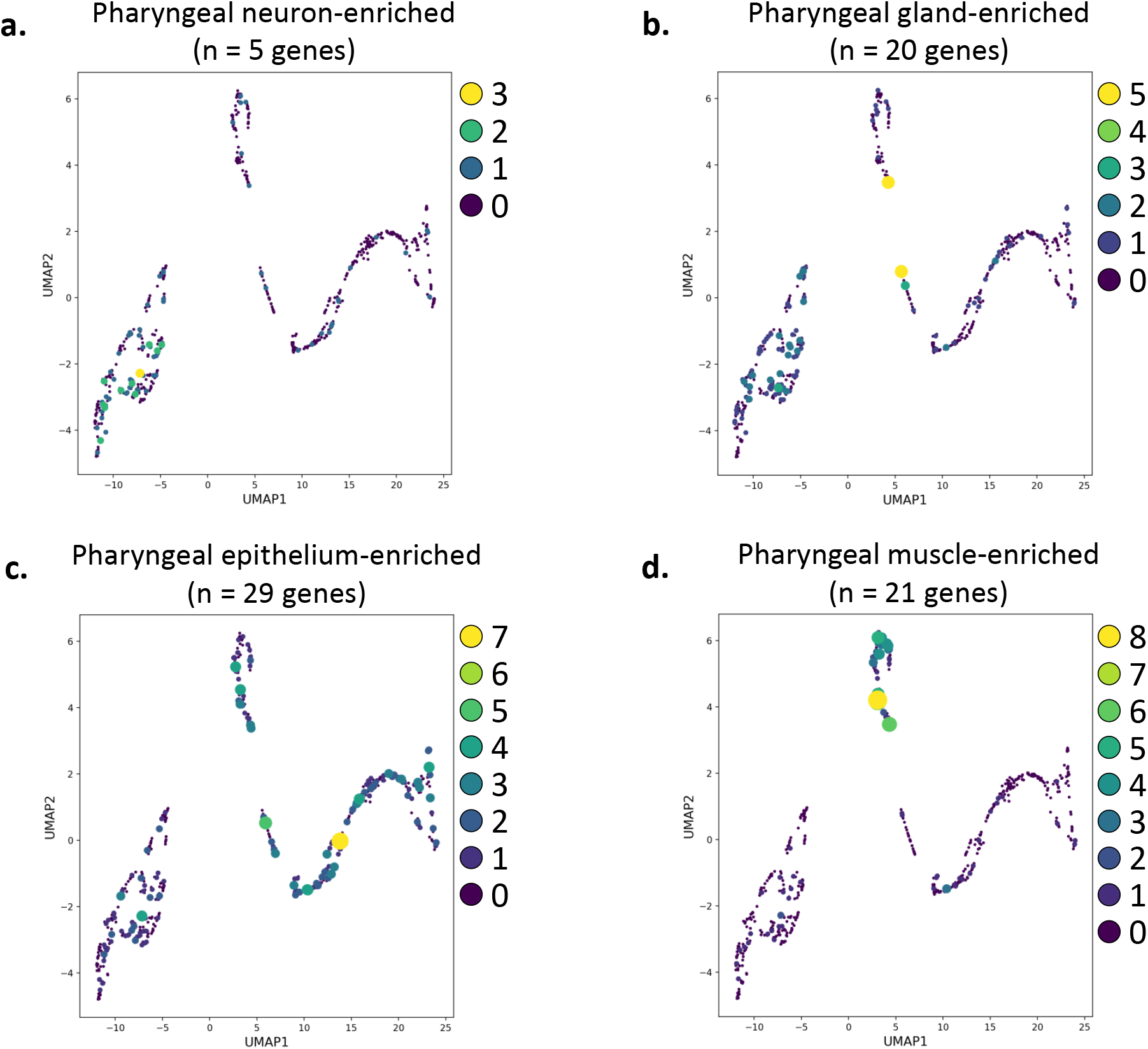
Marker genes identify pharynx subclusters. To identify pharynx subclusters, we assessed co-accessibility of sites near genes enriched in expression in the pharyngeal tissues identified in Cao et al. 2017 (Cao et al., 2017). The genes that we selected have greater than five-fold enrichment in the specified pharyngeal tissue compared to all other tissues, as reported by the GExplore website (http://genome.sfu.ca/gexplore/gexplore_search_tissues.html). Note that a relatively small subset of the genes matching the expression criteria in these tissues have nearby peaks, probably because our experiment recovered relatively few pharyngeal cells, reducing our power to detect pharynx-specific peaks. Nevertheless, we find that the genes with enriched expression in different pharyngeal tissues show nearby co-accessibility in cells with high probability in different topics. In particular, cells with high probability for topics 3 and 4 (Fig. S25) have more accessibility near genes expressed in pharyngeal neurons (**a.**), cells with high probability in topic 1 (Fig. S25) have more accessibility near genes expressed in pharyngeal gland (**b.**), cells with high probability in topic 2 (Fig. S25) have more accessibility near genes expressed in pharyngeal epithelium (**c.**), and cells with high probability in topic 0 (Fig. S25) have more accessibility near genes expressed in pharyngeal muscle (**d.**).

